# How we learn things we don’t know already: A theory of learning structured representations from experience

**DOI:** 10.1101/198804

**Authors:** Leonidas A. A. Doumas, Guillermo Puebla, Andrea E. Martin

**Affiliations:** University of Edinburgh; Max Planck Institute for Psycholinguistics Psychology of Language Department

**Keywords:** relation learning, predicate learning, neural networks, similarity, relative magnitude, invariance, learning structured representations

## Abstract

How a system represents information tightly constrains the kinds of problems it can solve. Humans routinely solve problems that appear to require structured representations of stimulus properties and relations. Answering the question of how we acquire these representations has central importance in an account of human cognition. We propose a theory of how a system can learn invariant responses to instances of similarity and relative magnitude, and how structured relational representations can be learned from initially unstructured inputs. We instantiate that theory in the DORA (*Discovery of Relations by Analogy*) computational framework. The result is a system that learns structured representations of relations from unstructured flat feature vector representations of objects with absolute properties. The resulting representations meet the requirements of human structured relational representations, and the model captures several specific phenomena from the literature on cognitive development. In doing so, we address a major limitation of current accounts of cognition, and provide an existence proof for how structured representations might be learned from experience.

---

To reason relationally is to reason about objects based on the relations that those objects play, rather than based on the literal features of those objects (see, e.g., Holyoak, 2012; Holyoak & Thagard, 1995). For example, when we make an analogy between the nucleus of an atom and the sun, we do so based on a common relation—e.g., that both nuclei and suns are *larger* than their orbiting bodies (planets and electrons respectively)—despite the fact that nuclei and suns are otherwise not particularly similar. Humans routinely draw inferences based on relations, from the mundane (“my kid won’t eat a portion that big”), to the sublime (“the cardinal number of the reals between 0-1 is larger than the cardinal number of the positive integers”), and relational reasoning has been shown to importantly contribute to abilities such as analogy (e.g., Holyoak & Thagard, 1995), categorisation (e.g., Medin, Goldstone, & Gentner, 1993), concept learning (e.g., Doumas & Hummel, 2004, Doumas & Hummel 2013), and visual cognition (e.g., Biederman, 1987, Hummel, 2013). In fact, the capacity to represent and reason about relations has been posited as the key difference in human and non-human animal cognition (Penn, Holyoak, & Povinelli, 2008).

Perhaps the most plausible explanation of how humans are able to reason relationally is that we can represent relations as abstract structures that take arguments—i.e., as predicates (see, e.g., Holyoak, 2012; Holyoak & Hummel, 2000). A predicate is a symbolic representational structure that can be dynamically bound to an argument, specifying some property about that argument (see e.g., Doumas & Hummel, 2005). Systems based on predicates or formally identical representations (i.e., structured or symbolic representations) are powerful and successfully account for many aspects of cognition (see Bringsjord, 2008 for a review). Providing an account of how these representations are learned from experience has proven difficult, however (e.g., Kriete et al., 2013; Leech, Mareschal, & Cooper, 2008; Rogers & McClelland, 2008). Models that rely on structured representations either make strong nativist claims, positing that a large set of representational elements and rules for building compositions of these elements are innate, or, at the very least, require that the powerful representations that they use are hand-coded by the modeller (e.g., Goodman et al., 2011; Kemp & Tenenbaum, 2009; Lake et al., 2015). The lack of an account of how structured representations might be learned from experience has been levied as one of the fundamental limitations of the symbolic approach to understanding cognition (e.g., Leech et al., 2008; McClelland, 2012; O’Reilly & Busby, 2002; O’Reilly, Busby, & Soto, 2003; Rogers & McClelland, 2008; Rummelhart, McClelland, & the PDP Research Group, 1986).

The goal of the present paper is to provide a theory of how the human cognitive system learns structured relational representations or predicates from experience. In the following we review the properties of predicate representations that make them hard to learn, and give an overview of the previous work in the domain of predicate learning. Next, we outline the set of three problems that must be solved in the service of learning abstract predicate representations from examples. We then describe a solution to these three problems instantiated in a computational model. Finally, we provide several simulations illustrating the success of the resulting system. In doing so, we address a major limitation of symbolic accounts of cognition, and provide, at the very least, an existence proof that the problem of learning structured predicate representations from unstructured experience is not fundamentally insurmountable.

## Why are predicates hard to learn?

Predicates—and other structured representations—as they are instantiated in theories of human mental representations, have two fundamental attributes (e.g., Doumas & Hummel, 2005, Doumas & Hummel 2012). First, a predicate specifies a property (or set of properties) about its argument in a manner that is invariant with that argument For example, the predicate *hairy* (*x*) can take (i.e., be bound to) any argument, and in doing so specify the truth of the property of ‘hairiness’ about that object. Binding *hairy* to dog (i.e., *hairy*(dog)) specifies the truth of ‘hairiness’ about the dog, and binding *hairy* to bear (i.e., *hairy*(bear)), specifies the truth of ‘hairiness’ about the bear.

Second, bindings between predicates and their arguments are dynamic—that is, they can be created and destroyed on the fly. While a predicate can be bound to an object to specify some property about the object, the binding can just as easily be broken. For example, the property *hairy* can be bound to dog, but if the dog is subsequently shaved, the binding between *hairy* and dog can be broken, and the dog can be bound to some other property (e.g., *bald*).

As a result of these properties, models that use structured representations are successful at accounting for a wide range of human cognitive phenomena. However, these properties that make relations so powerful, are also, in large part, what make accounting for how predicates are learned so difficult: While our representation of a relation like *taller* (*x,y*) is completely independent of any specific *x* or *y*, the instances from which we learn that relation in the world are exquisitely tied to very specific objects—that is, we do not get to experience instances of disembodied *taller*-ness in our environment. Every instance of *taller* that we experience in the world involves some specific object of a greater height than some other specific object. Predicates in the form that we end up representing them (abstract and invariant structures) just aren’t “out there” in the world of concrete and context-laden instances.

## Current approaches to the problem of learning predicates

While the symbolic approach to cognition dominated the field for decades (see Franklin, 1999), the difficulty in providing an account of how structured representations are learned has led to a tension in the field. Approaches to modelling human cognition now tend to fall into one of two camps. On the one hand, the connectionist (or eliminative) approach explicitly eschews predicates, solving the problem of where predicates come from by rendering it moot. On the other hand, current symbolist approaches make—either explicitly or implicitly—strong nativist claims about the origins of knowledge. We now outline these two approaches as well as a third more recent approach focused on attempting to address the problem of learning structured representations from experience.

### Eliminative connectionism: “We do not represent predicates”

The lack of an account of where symbols come from was one of the initial impetuses of the development of the connectionist approach, and it remains one of the primary motivations of more modern connectionist approaches (Leech et al., 2008; O’Reilly & Busby, 2002; O’Reilly, Busby, & Soto, 2003; Rogers & McClelland, 2008). In response to the question of how symbolic representations are learned to begin with, the prevailing answer in the connectionist tradition has been: They aren’t (e.g., McClelland, 2012).

Traditional connectionist models operate at the so-called sub-symbolic level. Connectionist representations are holistic, realized in patterns of activation in the system rather than structured. In fact, connectionist models explicitly eschew structured representations, and the lack of structured representations in these models was one of the core principles of the connectionist approach from its inception (see Rummelhart et al., 1986).

The persistence of the traditional connectionist approach is due, at least in part, to its wild successes (including those of its more recent offshoots, e.g., deep learning; e.g., LeCun et al., 2015). Connectionist models have been used to simulate a wide range of human cognitive phenomena from perceptual inference (Usher & McClelland, 2001) to language processing (Christiansen & Chater, 2001) to strategic video-game playing (Silver et al., 2016).

However, connectionist models have had limited success in domains like analogy-making and relational reasoning (Gentner & Forbus, 2011). The lack of symbolic representations in traditional connectionist systems may impose important restrictions in their capacity to simulate human level cognition. Indeed, some aspects of human cognition seem to require symbolic representations (e.g., solving cross-mappings [analogies where the relations point to one mapping, and the literal features of objects point to an orthogonal mapping]; e.g., Holyoak & Thagard, 1995; or integrating multiple relations when making an analogy; e.g., Spellman & Holyoak, 1992, 1996; or learning and using human language; e.g., Marcus, 1998, Pinker & Prince, 1988). Models without symbolic representations have repeatedly and systematically failed on these kinds of tasks, leading to arguments that systems without symbolic capacities are patently insufficient to account for the entirety of human cognition (e.g., Fodor & Pylyshyn, 1988; Holyoak & Hummel, 2000; Lake et al., 2017; Marcus, 1998; Pinker & Prince, 1988).

### Classic and neo-classic symbolism: “Necessary predicates are innate”

Unsurprisingly, models that use structured (i.e., symbolic) representations have had success in accounting for aspects of human cognition that traditional connectionist models have struggled with. For example, models with production system architectures like ACT (e.g., Taatgen & Anderson, 2008) have successfully accounted for a range of phenomena including problem solving, memory retrieval, and parsing of long-distance dependencies in sentence processing, while models of such as SME (Falkenhainer, Forbus, & Gentner, 1989), STAR (Halford, Wilson, & Phillips, 1998), and LISA (Hummel & Holyoak, 1997, 2003), successfully account for many phenomena from the literature on human relational reasoning.

More recently, Bayesian models have come to the fore as accounts of human cognition. Bayesian models of concept learning generally follow a learning-by-hypothesis-testing framework (e.g., Goodman et al., 2011; Kemp & Tenenbaum, 2009; Lake et al., 2015; for a notable counterexample, however, see Lu, Chen, & Holyoak’s, 2012, BART model). In these models, the system starts with a large set of representations and rules for combining them, and then learns combinations of these elements that best fit a given set of data. For example, in Lake et al.’s (2015) model of letter recognition, the system starts with representations of all possible line segments that could be used to construct a letter in any possible alphabet set, along with predicate representations for *connected-at-top* (*x*, *y*), *connected-at-bottom* (*x*, *y*), and *connected-at-middle* (*x*, *y*). The model might learn that the representation *connected-at-top* (squiggle1, squiggle4) is a legal letter in an alphabet, but all of the elements of the representation are present in the model before any learning occurs. Similarly, in Kemp & Tenenbaum’s (2009) account of relational concept learning, the model begins with all possible graph primitives and rules for combining them. The model might learn that a line of nodes is the best way to represent the legal leanings of supreme court justices, or that a hierarchical tree best describes inheritance in object-oriented programming languages, but, again, all those structures are represented by the model before any actual learning takes place.

These models do not provide an account of how the representations that they use might be learned in the first place. They require that the modeller hand code or pre-specify the representational structures that the model exploits, and, as a consequence, make very strong, though often implicit, nativist assertions.

### Learning structured representations from experience

Some models have attempted to account for the origins of abstract symbolic representations without positing innate sets of structured representations. In particular Lu et al.’s (2012) BART model, and Doumas, Hummel, and Sandhofer’s (2008) DORA model have been proposed as accounts of how relational representations might be learned from experience, rather than pre-specified.

BART (Lu et al., 2012) begins with feature lists generated by human subjects or via crawls of corpora. BART is then given pairs of items representing a particular relation (e.g., *bigger*). BART learns what amounts to second-order probability distributions over the features of the objects involved in particular roles (e.g., the *larger* and *smaller* roles of *bigger* (*x*,*y*)) using Bayes rule and some very general learning priors. Essentially, BART find properties associated with items in the world that embody particular relations.

The representations that BART learns are sufficient to solve a wide range of analogy problems, but the system struggles with some important edge cases. For example, it has difficulty with full relational reasoning (e.g., predicating that an atom can be *bigger* when the system has never experienced an instance in which an atom was bigger than something else), reasoning about counterfactuals, and reasoning outside of training range. However, the model makes a serious effort to account for the development of analogy making with minimal assumptions about the starting representations of the learning system.

In a similar vein, the DORA model (Doumas et al., 2008) provides an account for how structured representations (i.e., predicates) are acquired from unstructured representations (i.e., feature vectors). DORA begins with representations of objects as flat feature vectors. DORA uses a process of comparison-based feature extraction and self-supervised learning routines to learn structured representations of any invariant properties from the learning instances that it receives. For example, given a number of examples of instances where one object is larger than another object, DORA can extract and learn a structured representation of the invariant properties of all those larger things (e.g., it learns a predicate for *larger* from the properties that are invariant across instances of one larger and one smaller object). The resulting representations support successful reasoning in a wide range of relational tasks (see, e.g., Doumas & Hummel, 2010; Doumas et al., 2008; Hamer & Doumas, 2013; Martin & Doumas, 2017; Morrison et al., 2012; Son et al, 2010; a more detailed description of the model is also given below).

While the representations that DORA learns do not have the limitations of BART’s representations (Doumas et al., 2008), DORA does have the important limitation that it simply assumes a capacity to detect a set of invariant features that underlie the abstract concepts that it learns. The model, as it stands, does not provide much insight into what those invariant features are, or any insight into how they are acquired. Overcoming for this limitation and providing a more complete account of how structured representations can be learned from experience is a central focus of the current paper.

## What must be done to learn structured representations

A complete account of how we learn structured representations of abstract relations from experience entails solving three problems. First, the perceptual/cognitive system must learn to detect the basic featural *invariants* that remain constant across instances of the relation. That is, the perceptual system must deliver, or learn to deliver, an invariant response to instances of the to be learned relation. Second, the system must isolate those invariants from the other properties of the objects engaged in the relation to be learned. That is, given an activation vector delivered by the perceptual system in response to a stimulus (e.g., a visual scene wherein a table is larger than a hammer), the system must isolate those properties that code for the to be learned relation (e.g., the system must be able to separate the invariant features of *larger* from those encoding the rest of the scene). Third, the system must learn a *predicate* representation of the invariant relational properties—that is, it must come to represent them as an explicit entity that can be bound to arbitrary and novel arguments while remaining independent of those arguments.

The DORA model solves the second and third of these problems (see Doumas et al., 2008). We now present a model that solves the first problem for instances of similarity and relative magnitude (SRM), which form the basis of spatial relations, and we integrate this system with DORA. Below we give a very broad overview of our proposal for how the human cognitive system learns to respond in an invariant manner to instances of SRM. We then describe the DORA model, our instantiation of the SRM detection model, their integration, and provide an example of how they combine to learn structured representations of SRM. Next, we present simulations providing a proof of concept that SRM relations can be learned without hardcoding relational primitives. Subsequently, we show that the DORA model accounts for several developmental trends observed in empirical studies. Finally, we discuss implications and extensions of the current model, and suggest directions for future research.

## A theoretical proposal for detection and invariant responding to similarity and relative magnitude

Below we present a theory of how the cognitive system learns invariant implicit responses to similarity and relative magnitude (SRM). The core theoretical claims of our proposal and their instantiations in the current model are outlined in Table 1. These core theoretical claims are as follows: (1) The invariant codes for SRM are a property of invariant neural responses that arise as a function of comparison. (2) During comparison compared items are co-activated, and co-activate their constituent distributed representations. (3) The similarity of the two compared items can be established by a measure of the match in the resulting firing pattern of the semantic properties. (4) The cognitive system learns invariant codes of basic similarity as well as relative “same” and “different” by exploiting the invariant patterns of firing in (3). (5) The cognitive system detects relative magnitude by directly comparing absolute neural response in the neural system. (6) A specific and invariant firing pattern emerges when two items of greater and lesser magnitude are compared, and a different specific and invariant pattern emerges when two items of the same magnitude are compared. (7) The cognitive system learns invariant codes for “same”/“different”, “more”/“less” by exploiting the invariant patterns of firing in (6).

**Table 1.** Core theoretical claims of the model for similarity and relative magnitude detection, and their instantiations in the current model.

| **Core theoretical claim** | **Instantiation in the current model** |
| --- | --- |
| 1. The invariant codes for SRM are a property of invariant neural responses that arise as a function of comparison. | 1. The basic mechanisms for learning invariant implicit responses to relative SRM emerge from comparing representations. |
| 2. During comparison compared items are co-activated, and co-activate their constituent distributed representations. | 2. Comparison is a process of two units becoming active together. |
| 3. The similarity of the two compared items can be established by a measure of the match in the resulting firing pattern of the semantic properties. | 3. Similarity is computed directly the overlap of compared sets of distributed representations. |
| 4. The cognitive system learns invariant implicit codes (features) of basic similarity as well as relative “same” and “different” by exploiting the invariant patterns of firing in (3). | 4. DORA (see below) will learns to associate the response of very high overlap with “same”, other overlap levels with “different”. |
| 5. The cognitive system detects relative magnitude by directly comparing absolute neural response in the neural system. | 5. During magnitude comparison, two POs compete to stay active after activating the units that code absolute dimensional magnitude. |
| 6. A specific and invariant firing pattern emerges when two items of greater and | 6. When two items that code for different absolute magnitudes are compared and |
| lesser magnitude are compared, and a different specific and invariant pattern emerges when two items of the same magnitude are compared. | compete to become active, the item coding for the greater magnitude (and thus receiving input from more units) will win the competition and remain active; complementarily, the item coding for the lesser magnitude (and thus receiving input from fewer units) will lose the competition and become active later. By contrast, when two items that code the same absolute magnitude are compared and compete to become active, the competing units will receive the same input, and settle into a state of mutual low activation. These patterns are a direct consequence of how distributed networks behave. They are not unique to the current model. |
| 7. The cognitive system learns invariant implicit codes (features) for “same”/“different”, “more”/“less” by exploiting the invariant patterns of firing in (6). | 7. DORA (see below) exploits the invariant responses described in (7). Features learn to respond to the winner, the loser, and simultaneously active competing POs. DORA learns excitatory connections between the POs and the features that respond to them by Hebbian learning. These features then |
|  | serve as the invariant semantic codes for<br>“more”, “less”, and “same”. |

### Invariant responses to similarity and difference

Diagnostic and information rich patterns emerge when a system based on distributed representations compares items. Consider a very simple case where the system compares a distributed representation of an apple to a distributed representation of a fire-engine. A node in the network is connected to a set of features coding for the apple in a distributed fashion, and another node in the network is connected to a set of features coding for the fire-engine in a distributed fashion (Figure 1a). When the two objects are compared, the units coding for the apple and those coding for the fire-engine are co-activated (Figure 1b). When both the apple and fire-engine representations are co-active, any features common to both the apple and the fire-engine will receive input from two sources and thus become roughly twice as active as any features unique to either the apple or the fire-engine (Figure 1c). This process of comparison highlights the intersection of the apple and the fire-engine (i.e., their shared features). Comparison-based intersection discovery has been exploited in models like LISA (Hummel & Holyoak, 1997, 2003) and DORA to support processes such as analogical mapping, schema induction, and learning structured representations from unstructured representations of simple objects.

**Figure 1.**
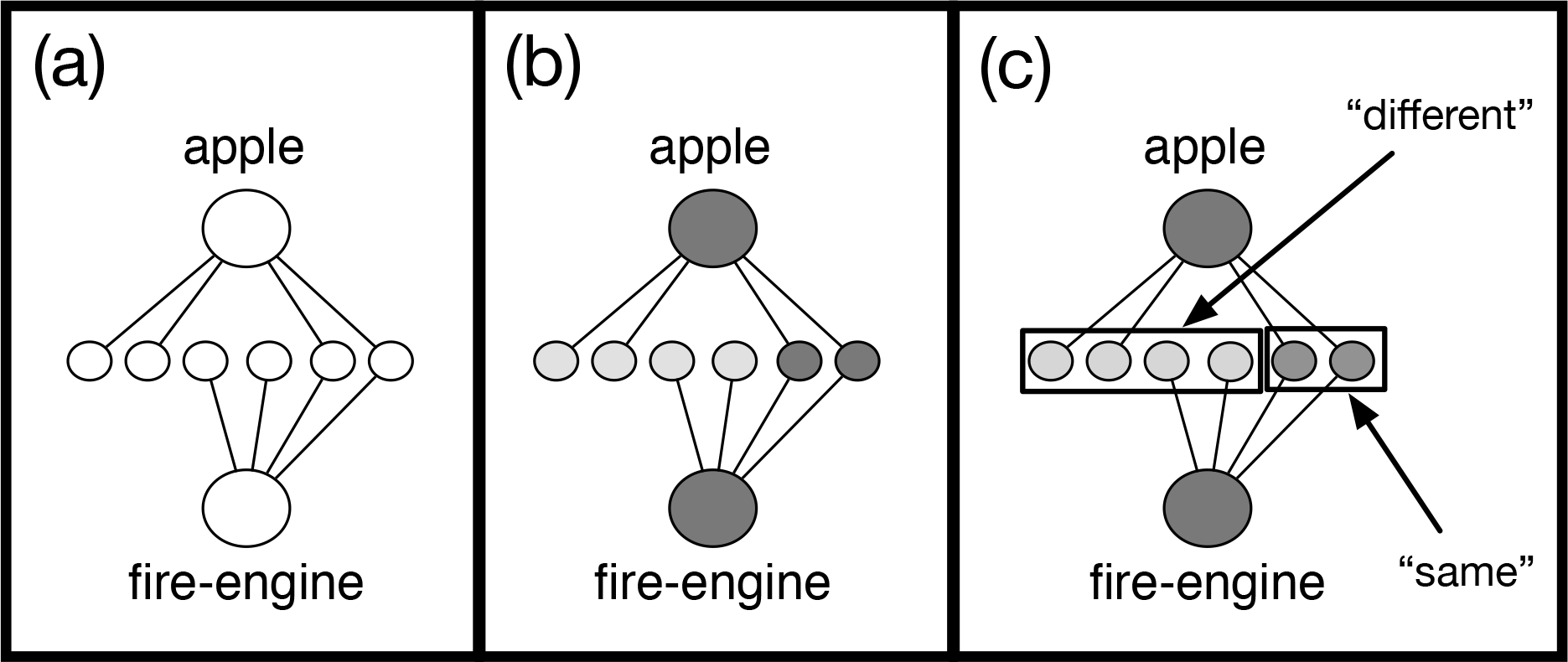
An illustration of comparison-based intersection highlighting. (a) When objects coded by distributed feature sets are compared (i.e., are co-activated), shared features receive input from multiple sources, while features unique to only one of the objects receive input only from a single source. (b) Shared features, therefore, become more active than unshared features. (c) This difference in activation marks what is similar and different about the compared objects: More active features are properties that are similar, while less active features are properties on which the two objects are different.

Crucially, the pattern of differentially active units that arises during comparison—with similarities marked by more active units and differences by less active units (Figure 1c)—will arise whenever two distributed representations are co-activated. As a consequence, the signal serves as an invariant indicator for basic similarity (or sameness) and difference. Learning an invariant response for sameness or difference, therefore, reduces to learning to respond differentially to these two patterns. As we show below, an invariant feature for “same” and “different” learned in this manner supports solving very simple similarity and difference problems, and also serves as the basis for learning structured (i.e., symbolic) representations of abstract *same* and *different* that support more complex problem solving.

### Invariant responses to relative magnitude

Just as basic comparison of distributed representations serves to mark basic “sameness” and “difference”, so comparison also serves as the basis for responding in an invariant manner to relative magnitude information (i.e., “moreness”, “lessness”, and “sameness”).

A comparison-based solution to the problem of learning an invariant feature response for “more”, “less”, and “same” requires the assumption that initial absolute magnitude information is coded by a direct neural proxy: All else being equal, higher magnitude items are coded by more neurons—or with a higher rate of firing, or signal amplitude, etc.—than comparatively lower magnitude items. There is a large amount of evidence for this assumption. For example, in visual processing, larger items take up more space on the retina, and are coded by larger swaths of the visual cortex (e.g., Engel et al., 1994; Wandell, 1995).

Basic magnitude computation follows naturally from this coding mechanism. To illustrate, consider a case where two objects of different size are compared (Figure 2a). If the two representations of absolute size are co-activated, and the units coding the two objects then compete to become active, or to stay active (e.g., via lateral inhibition; Figure 2b), then the larger object will tend to become active first (Figure 2c). When that object becomes inactive (e.g., via a yoked inhibitor unit—see, Doumas et al., 2008; Hummel & Holyoak, 1997, 2003; von der Malsburg, 1981, 1999), the smaller object can become active (Figure 2d). In brief, the process of attempting to co-activate two competing representations of objects that code for some form of absolute magnitude information will result in a pattern of firing wherein the item coding the larger magnitude becomes active at a different time scale than the item coding the smaller magnitude. By contrast, when two items of the same magnitude are compared, the two items will settle into a state of mutual co-activation (i.e., both units will remain somewhat active until their inhibitors fire).

**Figure 2.**
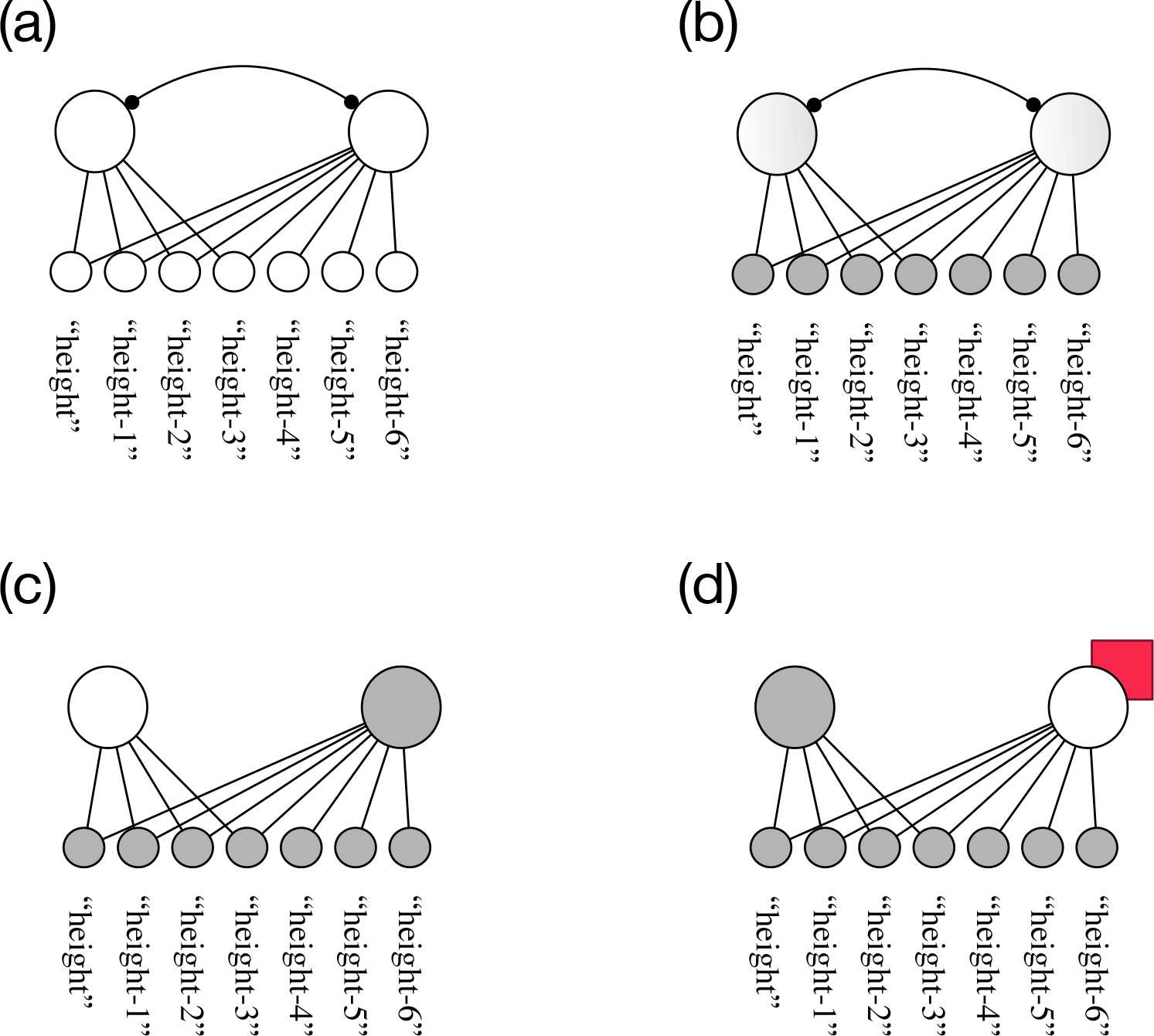
An illustration of the basic invariant pattern that emerges when two items coding basic absolute magnitude information compete to become active. (a) Two units (large circles) coding for absolute magnitude. (b) The semantics (small circles) coding for absolute magnitude are activated, and the two units compete (via lateral inhibition; line with circle endpoint) to become active. (c) The unit attached to the greater magnitude wins the competition and becomes active first. (d) After the winning unit is inhibited to inactivity by a yoked inhibitor (represented here by the small square behind large circle), the unit attached to the lesser magnitude becomes active.

Consequently, comparing items with values on dimensions or extents result in invariant signals for basic relative magnitude information (e.g., basic “moreness”, “lessness”, and “sameness”). Learning invariant features to code for “more”, “less”, and “same”, therefore, reduces to learning to respond differentially to these patterns. For example, the invariant feature for “more” might simply be the node (or nodes) that respond to the compared item that becomes active first; the invariant feature for “less” is then the node (or nodes) that respond to the compared item that that becomes active next; the invariant feature “same” is the node or (nodes) that respond to two mutually co-active items. As we show below, invariant features for “more”, “less”, and “same” learned in this manner serve as the basis for learning structured (i.e., symbolic) representations of abstract *more*, *less*, and *same* that support more complex problem solving.

In the next section we describe a theory of how structured (i.e., symbolic) representations of SRM can be learned from unstructured representations of extent. Our theory is embodied in a computational model called DORA.

## A theory and computational model for learning structured representations of similarity and relative magnitude

As noted above, it is one thing to respond in an implicit manner to SRM, however, to achieve human level cognitive performance, a system must be able to learn explicit structured representations of these invariant implicit responses. A solution to the problem of how the implicit (featural) invariant responses to SRM are generated and then support learning structured (i.e., symbolic) representations of *same*, *different*, *more*, and *less*, is given by combining a system that learns to respond to the invariant neural responses to SRM with the DORA model (Doumas et al., 2008) that learns structured representations from unstructured representations.

DORA is based on a set of four (additional) core theoretical claims, which we outline along with their instantiations in the current model in Table 2. (1) Mature human mental representations are formally similar to a role-filler binding system (see Doumas & Hummel, 2005; Doumas et al., 2008; and below), which reduces the problem of learning structured representations of multi-place relations to the comparatively simpler problems of learning single-place predicates coding object properties, and then linking sets of corresponding single-place predicates to form multi-place relational representations. (2) Comparison can lead to the discovery and predication of shared object properties. (3) A common vocabulary of representational elements forms the basis for both predicates and their arguments. (4) Mapping sets of predicates of smaller arity can lead to the formation of higher arity (i.e, multi-place relational) structures.

**Table 2.** Core theoretical claims of the DORA theory, and their instantiations in the DORA model.

| Core theoretical claim | Instantiation in DORA |
| --- | --- |
| Mature human mental representations are formally similar to a role-filler binding system, which reduces the problem of learning structured representations of multi- | DORA represents whole relational structures using LISAese, or as linked sets of role-filler pairs. |
| place relations to the comparatively simpler problems of learning single-place predicates coding single object properties, and then linking sets of corresponding single-place predicates to form multi-place relational representations. |  |
| Comparison can lead to the discovery and predication of shared object properties. | DORA uses intersection discovery based on comparison (co-activation) of representations, along with asynchrony based binding to learn and refine single-place predicate representations of the shared properties of compared representations. |
| A common vocabulary of representational elements forms the basis for both predicates and their arguments. | There is a single pool of semantic features shared by all predicates and objects in DORA. |
| Mapping sets of predicates of smaller arity can lead to the formation of higher arity structures. | DORA learns higher arity (i.e., multi-place) relational structures by linking sets of role-filler pairs that map to structurally similar sets of role-filler pairs. |

Below we describe the DORA model and its operation along with a model for generating and learning invariant SRM responses. We then provide an example of learning structured representations with an integrated DORA and SRM detection procedure. In the simulations section that follows, we show how DORA and the SRM detection procedure provide a solution to the problem of learning structured representations of similarity and relative magnitude functions from unstructured inputs with very minimal assumptions about innate knowledge, and account for a number of general and specific phenomena from the literature on human learning and development.

### Overview of the DORA model

DORA is a model of how structured (i.e., functional predicate) representations can be learned from unstructured representations without assuming an a priori set of structured predicates, or an innate explicit formal language. That is, DORA provides an account of how structured, symbolic, representations can be learned from scratch.

DORA is descended from Hummel and Holyoak’s (1997, 2003) LISA model. LISA is a highly successful model of many high-level cognitive phenomena, and DORA is a model of how the powerful representational currency upon which LISA’s reasoning is based (LISAese), can be learned in the first place. As DORA learns, it eventually becomes a version of the LISA model (i.e., once it has learned a set of representations, it operates like a version of LISA with those representations). To date, LISA and DORA have been used to account for over 50 empirical phenomena in domains such as analogy, memory retrieval, inductive generalisation, structured representation learning, concept development, the development of analogical reasoning, and the development of object recognition (e.g., Doumas & Hummel, 2010; Doumas et al., 2008; Hummel & Holyoak, 1997, 2003; Livins, Spivey, & Doumas, 2015; Martin & Doumas, 2017; Morrison et al., 2004; Morrison et al., 2012; Son et al., 2012).

As noted above, DORA is based on the set of core theoretical principles laid out in Table 2 (see also Doumas et al., 2008). Below we describe DORA and its basic operations. Full details of the model as well as the majority of equations are given in Appendix A. We begin by discussing representations in DORA—both the representations the model starts with, and the representations it learns (Table 2, theoretical points 1 and 3). We then describe a model for SRM detection based on the theoretical principles laid out in Table 1. Next, we describe how DORA learns and then refines representations of single place predicates (Table 2, theoretical point 2). Finally, we describe how DORA learns multi-place relational structures from sets of lower arity predicates (Table 2, theoretical point 4).

### Representation in DORA

DORA begins with representations of objects coded as flat feature vectors (see Figure 3a), These representations are similar to representations in traditional connectionist architectures where elements are coded by distributed collections of units. For example, DORA might represent a toy ball with a node connected to a set of features (Figure 3a).^1^ In short, DORA begins with objects coded conjunctively by flat feature vectors. (In terms of cortical computation, feature nodes can be thought of as aggregate units, perceptual representations, or activation states over networks.)

**Figure 3.**
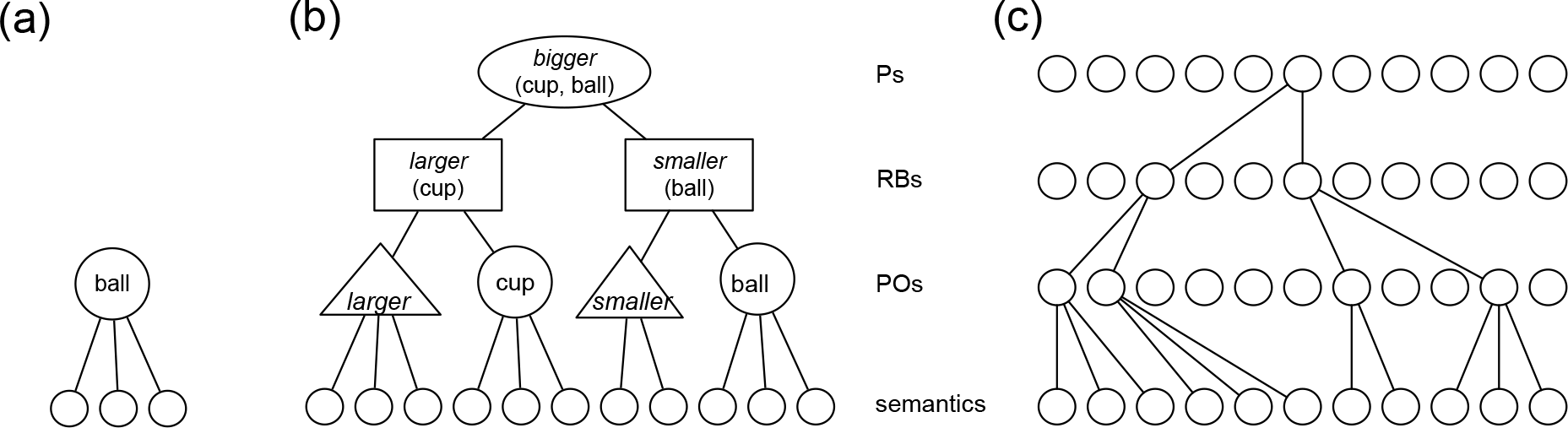
Representations in DORA. (a) DORA’s starting state. DORA begins with representations of objects connected to lists of their features. (b) LISAese representation of the proposition *bigger* (cup, ball). DORA learns full LISEese representations from examples of representations like those in (a). We use larger circles, triangles, rectangles, and ovals for the purposes clearly differentiating units in different layers of the network. Importantly. (c) More conventional depiction of a LISAese proposition. The proposition is instantiated in layers of bidirectionally connected nodes.

DORA learns representations of a form we call LISAese (Figure 3b-c). Full propositions in LISAese are coded by layers of units in a connectionist computing framework (Figure 3b). At the bottom of the hierarchy, semantic (or feature) nodes (small circles in Figure 3b) code for the featural properties of represented instances in a distributed manner. At the next layer, localist predicate and object units (POs; triangles and large circles in Figure 3b) conjunctively code collections of semantic units into representations of objects and roles. At the next layer localist role-binding units (RBs; rectangles in Figure 3b) conjunctively bind object and role POs into linked role-filler pairs. Finally, proposition units (Ps; ovals in Figure 3b) link RBs to form whole relational structures.

As an example, consider a LISAese representation of the proposition *bigger* (cup, ball), as depicted in Figure 3b. PO units representing the relational roles *larger* and *smaller*, and the fillers cup and ball, are connected to semantic units coding their semantic features. At the next layer of the network, RB units conjunctively connect a specific role to a specific filler. Specifically, one RB unit conjunctively connects cup to *bigger*, and one conjunctively connects ball to *smaller*. At the top of the hierarchy, a P unit links the RBs representing *larger*+cup and the RB representing *smaller*+ball to form a whole relational structure. The entire hierarchy of units then encodes the relational proposition *bigger* (cup, ball).

While we use different shapes (e.g., large circles, triangles, rectangles, and ovals) to indicate nodes at different layers, these are not different types of nodes. Rather, we use different shapes solely for the purpose of clarifying different layers of nodes. The same proposition can be represented in a more traditional format as layers of bidrectionally connected nodes (Figure 3c).

### Computational macrostructure

Propositions in DORA are divided into four mutually exclusive sets (Figure 4): the *driver*, the *recipients*, *long-term-memory* (LTM), and the *emerging recipient* (EM). Each set consists of a layered network coding for POs, RBs, and Ps (i.e., there are specific layers coding for POs, RBs, and Ps in the driver, and another set of layers coding for POs, RBs, and Ps in the recipient). Semantic units are common across all networks (i.e., driver, recipient, LTM, and EM units are connected to the same pool of semantic units).

**Figure 4.**
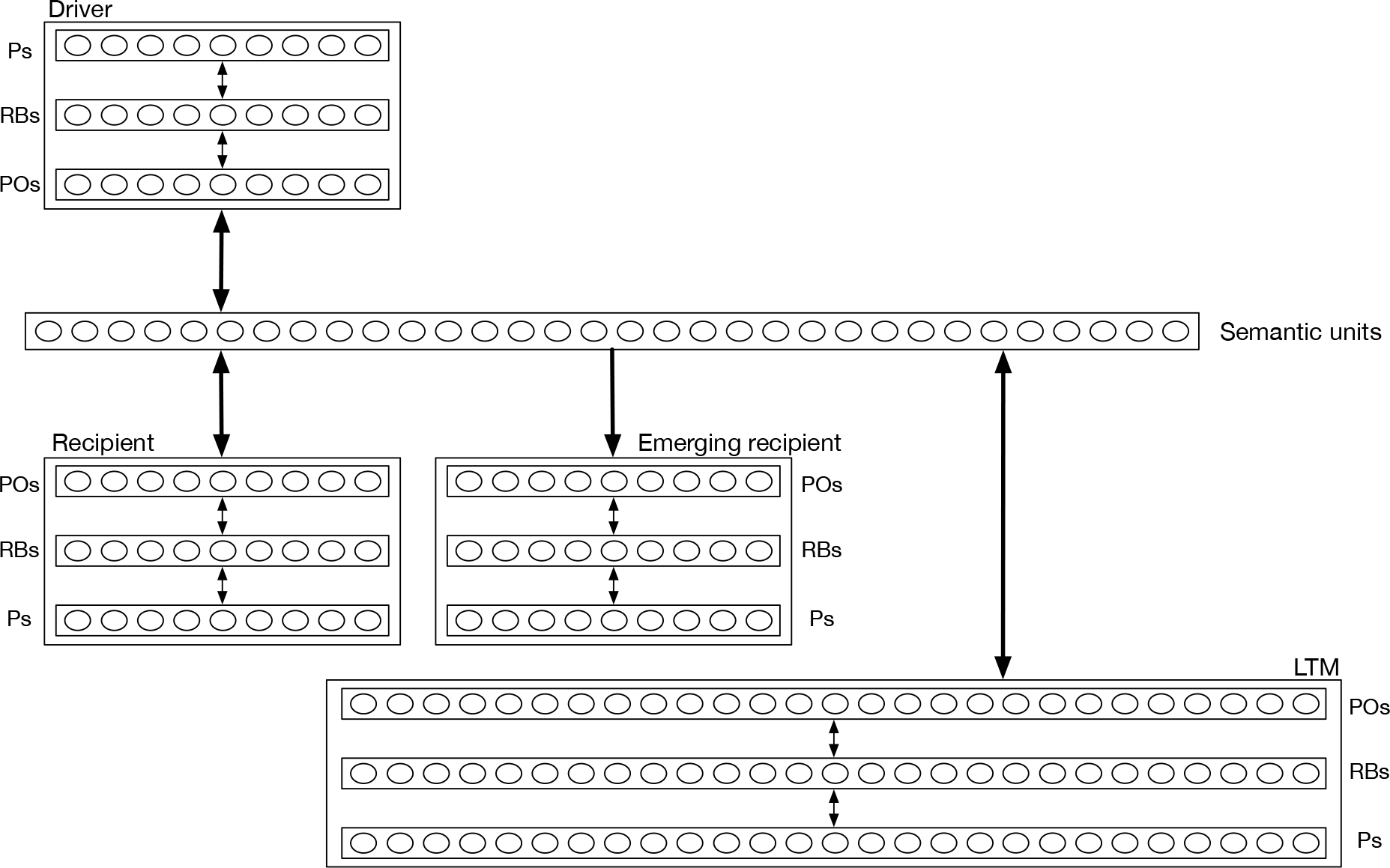
DORA’s computational macrostructure.

An *analog* in DORA is a complete story, event, or situation. Analogs are represented by a collection of token (P, RB and PO) units that together represent the propositions in that analog. While token units are not duplicated within an analog (e.g., within an analog, each proposition that refers to Don connects to the same “Don” unit), separate analogs have nonidentical token units (e.g., Don will be represented by one PO unit in one analog and by a different PO in another analog). All analogs are connected to the same pool of semantic units. The semantic units thus represent general type information and token units represent instantiations of those things in specific analogs (Hummel & Holyoak, 1997, 2003). For example, if in some analog, the token (PO) unit “Fido” is connected to the semantics “animal”, “dog”, “furry” and “Fido”, then it is a token of an animal, a dog, a furry thing and of the particular dog Fido.

### Basic processing

The driver controls the flow of activation in DORA, and corresponds to the DORA’s current focus of attention. Units in the driver pass activation to the semantic units. Because the semantic units are shared by propositions in all sets, activation flows from the driver to propositions in the other three sets. All of DORA’s operations (i.e., *retrieval*, *mapping*, *predicate learning*, *relation formation*, *schema induction*, and *generalisation*) proceed as a product of the units in the driver activating semantic units, which in turn activate units in the various other sets (as detailed below). During *retrieval*, patterns of activation generated on the semantic units by units in the driver, activate representations in LTM, which are retrieved into the recipient. Propositions in the recipient are available for *mapping* onto propositions in the driver. Active driver units activate corresponding (i.e., semantically similar) units in the recipient, allowing DORA to learn mapping connections between them. Mappings between units in the driver and recipient are the basis of DORA’s ability to *learn new predicate representations* and to *form higher arity structures* from lower arity structures. Finally, DORA can *learn schemas* from mapped propositions in the driver and recipient, which are encoded into the EM, and may subsequently be encoded into LTM and later enter the driver or recipient.

When a proposition in the driver becomes active, role-filler bindings must be represented dynamically on the units that maintain role-filler independence (i.e., POs and semantic units; see Doumas & Hummel, 2005; Doumas et al., 2008; Hummel & Holyoak, 1997, 2003). In DORA, roles are dynamically bound to their fillers by systematic asynchrony of firing. DORA maintains an asynchrony of firing at either the level of POs, which we term *asynchronous binding*, or the level of RBs, which has been termed *synchronous binding*. During asynchronous binding, as a proposition in the driver becomes active, bound roles and objects fire in direct sequence. (e.g., with roles firing directly before their fillers; see Figure 5a). For example, to bind *bigger* to cup and *smaller* to ball (and so represent *bigger* (cup, ball)), the units corresponding to *larger* fire (Figure 5a[i]) directly followed by the units corresponding to cup (Figure 5a[ii]), followed by the units for coding *smaller* (Figure 5a[iii]) followed by the units for ball (Figure 5a[iv]).

**Figure 5.**
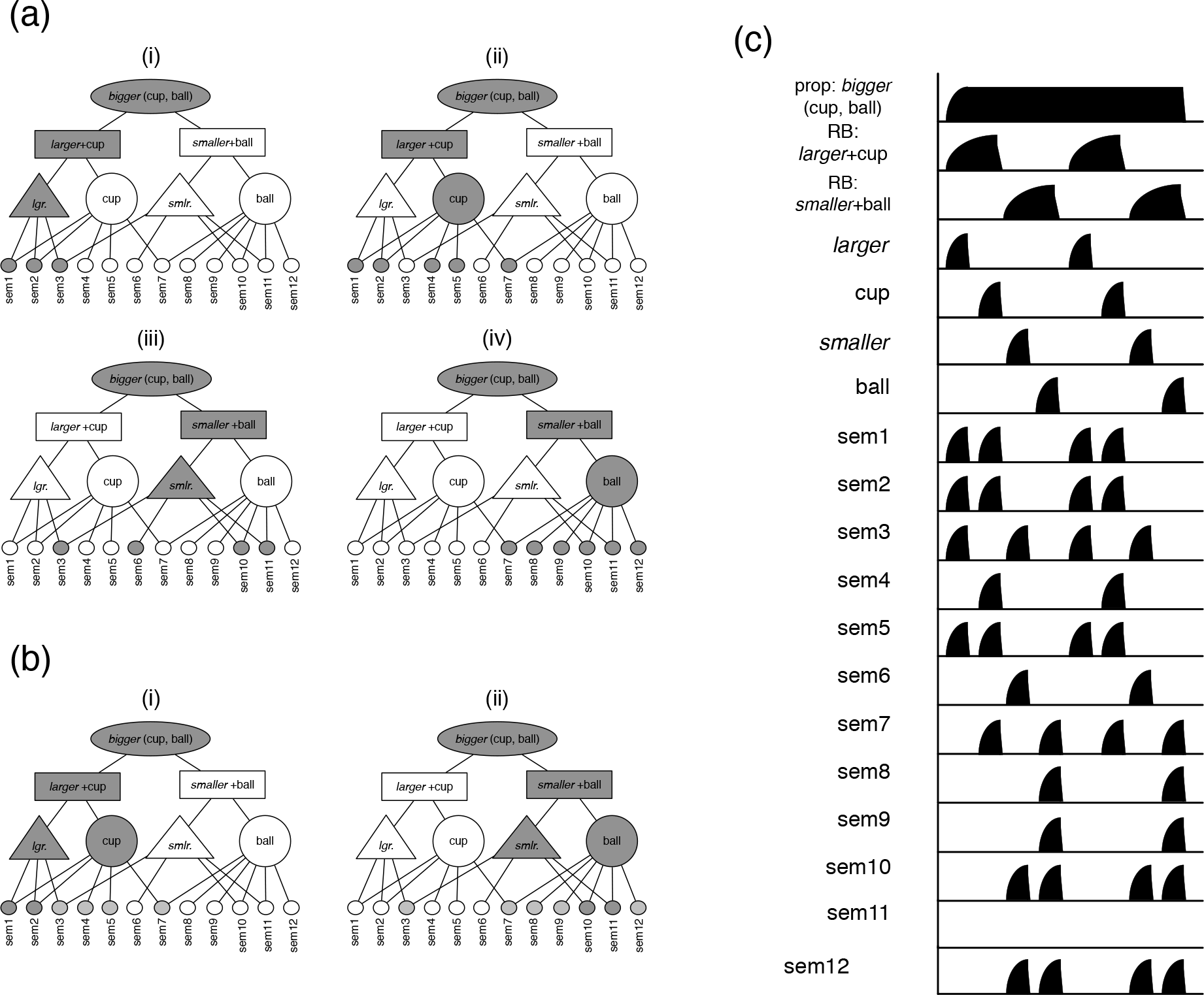
Binding in DORA. Binding in DORA occurs via systematic asynchrony of firing. The asynchrony is maintained at either the level of POs (a) or RBs (b). When the asynchrony is maintained at the level of POs, to represent the binding between a role and a filler, units coding for the role fire directly in sequence with units coding for a filler, and out of synchrony with other role-filler sets. Grey units are active. (a[i-ii]) The binding of *larger* to cup is carried by the sequence of firing of the units representing *larger* a[i] followed by the units representing cup a[ii]. (a[iii-iv]) The binding of *smaller* to ball is carried by the sequence of firing of the units for *smaller* a[iii] followed by the units for ball a[iv]. When the asynchrony is maintained at the level of RBs, to represent the binding between a role and a filler, units coding for one role-filler pair fire out of synchrony with the units coding for another role-filler pair. The binding of *larger* to cup is carried by the firing of the units coding for *larger* and cup firing together (b[i]) and out of sequence with the units coding for *smaller* and ball (b[ii]). (c) A time series illustration of the firing of units during asynchrony-based binding (as in (a)).

In brief, bound role-filler pairs fire as couplets, and role-filler sets from the same proposition fire in sequence. For instance, to represent both *bigger* (cup, ball) and *bigger* (circle star), the units representing *larger* would fire, followed by the units representing cup, then the units representing *smaller* fire, followed by the units representing ball. Next, the units representing *larger* fire, followed by the units representing circle, then the units representing *smaller* fire, followed by the units representing star (see Figure 6). In binding by systematic asynchrony, binding information is carried by when units fire. Role-filler bindings are dynamic and represented explicitly, while role-filler independence is maintained (see, e.g., Doumas et al., 2008). Consequently, there is no need to use different types of units (as in SME; Fakenheiner et al., 1989; Forbus et al., 1995; or STAR; Halford et al., 1998) or different sets of units (as in LISA; Hummel & Holyoak, 1997, 2003) to represent relations/relational roles and objects and their semantic properties. Accordingly, in DORA, roles and objects are both coded by a common pool of semantic units, and, importantly, as detailed below, this capacity allows DORA to learn representations of object properties and relational roles from representations of objects.

**Figure 6.**
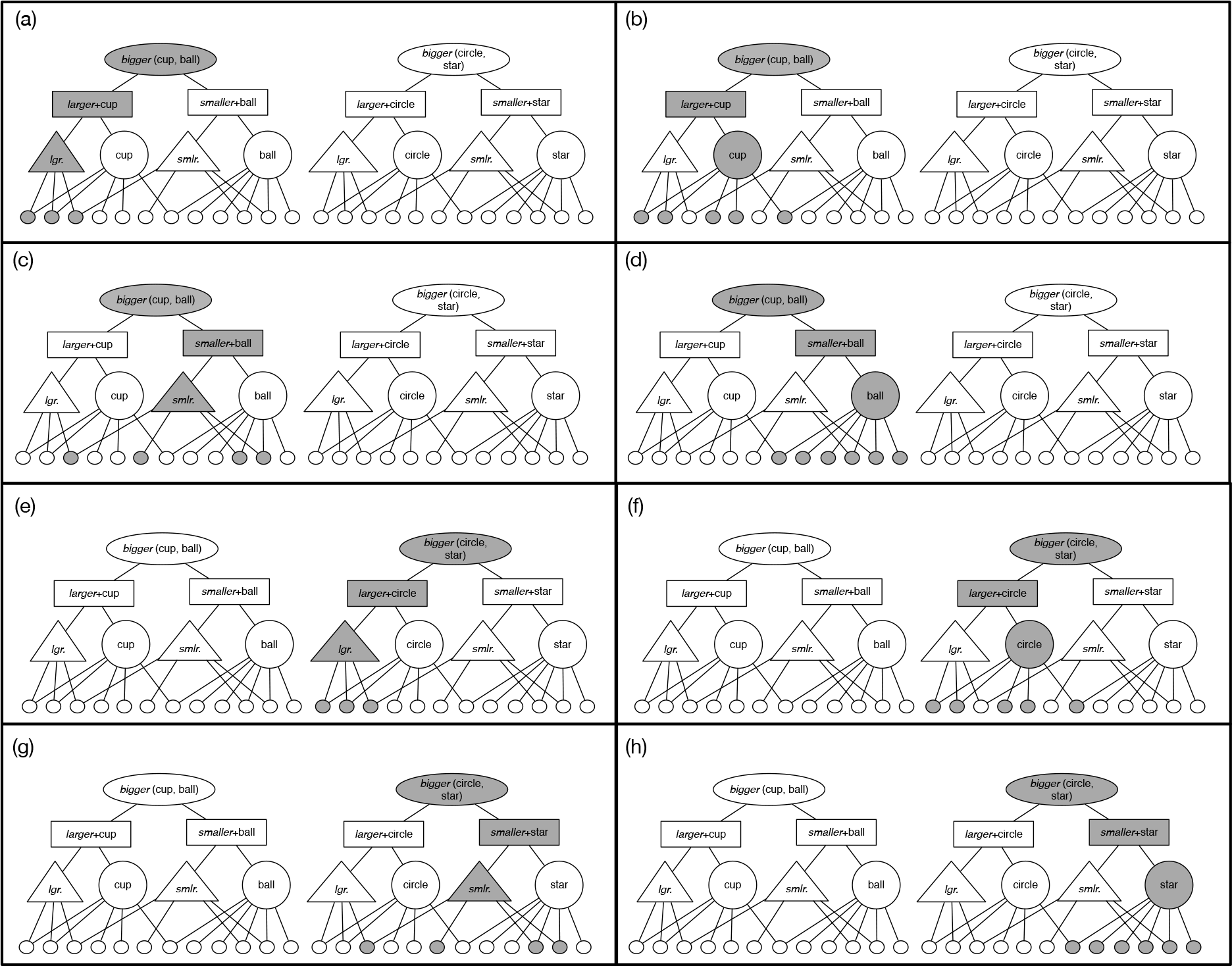
Binding roles to fillers across multiple propositions in DORA. (a-d) DORA represents *bigger*(cup, ball), representing the binding of *larger* to cup (a-b) and *smaller* to ball (c-d). (e-h) DORA represents *bigger* (circle, star), representing the binding of *larger* to circle (e-f), and *smaller* to star (g-h).

DORA can also maintain a systematic asynchrony at the level of RBs, such that bound roles fire in synchrony with their fillers (Figure 5b). This temporal binding signal is the same as that used by LISA (Hummel & Holyoak, 1997, 2003). When role-filler pairs fire in synchrony, their semantic patterns are superimposed. To keep role and filler representations distinct, as is necessary for many forms of learning, including schema induction and relational generalization, and the representation of propositions or sentences (Martin & Doumas, 2017), different pools of units are required to code for the properties of objects and roles, which makes learning relational roles and multi-place relations from object representations much more difficult (see Doumas et al., 2008). However, the resources required to bind role-filler pairs are halved during synchronous binding as opposed to asynchronous binding—specifically, binding a role-filler pair by asynchrony requires that two spikes of unit activation be maintained and kept distinct, but synchrony requires only one spike of unit activation. As such, DORA uses synchronous binding for tasks that do not require that role and filler semantics be differentiated (e.g., retrieval and mapping), and asynchronous binding for tasks that require distinct representations of role and filler semantics (e.g., learning). The model thus makes the prediction that representing a proposition should require more WM (or other processing resources) during learning than during mapping. This prediction appears to hold for humans (see, e.g., Saiki, 2003).

### A process model for invariant feature responses to SRM

As described below, DORA provides a solution to the problem of learning structured representations of invariant signals present in the input. A fundamental problem, however, remains: what are these invariants are and how do we learn to detect them in the signal? One solution is to assume that the perceptual system comes equipped with the capacity to detect (or at least very quickly learn to detect) the necessary invariant properties. For some properties that we learn structured representations of such as colour or absolute values on dimensions, this explanation is perhaps sufficient. However, it is a bit more difficult to imagine what the invariant properties of more abstract concepts like similarity or relative magnitude might be. It is, of course, possible that machinery for the detection of invariant abstract properties is innate, but it would be more satisfying to account for how a system might learn to detect these properties from experience.

Here we present a process model that instantiates the core theoretical claims that comparison facilitates learning an invariant response to instances of SRM (Table 1). In this section, we provide an overview of the model in more conceptual terms. Details of the model are given in Appendix A. There are several algorithms that can be levied to learn these basic responses to SRM. We use a simple algorithm based on unsupervised Hebbian learning (see Appendix B). While perhaps not the most elegant algorithm that might be used, the present descripion has the advantage of being easy to implement and (comparatively) easy to follow. Certainly, more complex algorithms based on reinforcement learning (e.g., Sutton & Barto, 1998), threshold node tuning (e.g., Huang et al., 2006), or free energy minimization (e.g., Friston, 2010), could also be used to solve the problem.

As noted previously, a solution to the problem of learning invariant features that respond to instances of SRM requires the assumption that initially available absolute magnitude information is coded by a direct neural proxy. There is, however, a preponderance of evidence for this assumption (e.g., Engel et al., 1994; Wandell, 1995).

Basic relative magnitude detection is accomplished by comparison. When the model compares two representations with specific magnitude values (e.g., two POs attached to absolute size are present in the driver; Figure 7a, the units coding their absolute magnitudes are co-activated, and the PO units compete via lateral inhibition (Figure 7b). The POs will eventually settle, with either one PO becoming more active and inhibiting the other to inactivity, or, as is the case if both POs code for the same absolute magnitude, with both POs in a steady state of co-activation (Figure 8). Additional semantic units respond to the pattern of firing in the driver POs. Some units are excited by two active POs in the driver, others are excited by a single highly active PO early in firing, or by a single highly active PO late in firing (these regions of excitement are easily learnable via unsupervised learning; a description of this process is given in Appendix B). The active POs learn connections to the active semantic unit by Hebbian learning, or:

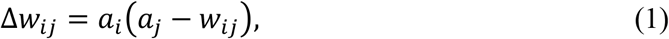

where, *w*_*ij*_ is the weight between PO unit *i* and semantic unit *j, a*_*j*_ is the activation of semantic unit *j*, *a*_*i*_ is the activation of PO unit *i*.

When a single PO unit wins the competition, it becomes active first (Figure 7c). By Eq. 1, hat unit learns connections to any semantic units that are activated early in the SRM competition (Figure 7d). When that active PO becomes inhibited (e.g., due to its yoked inhibitor; Figure 7e), the second PO (the one initially inhibited by the winning PO) will become active (Figure 7f). By Eq. 1, that unit learns connections to the semantics that are activated late in the SRM competition (Figure 7g). In short, the pattern of a unit winning the activity competition and another losing are invariant responses to greater and less-than in a network that compares absolute magnitudes. Any units that respond to these patterns come to code for the qualities of relative “more” and relative “less” respectively (Figure 7h).

**Figure 7.**
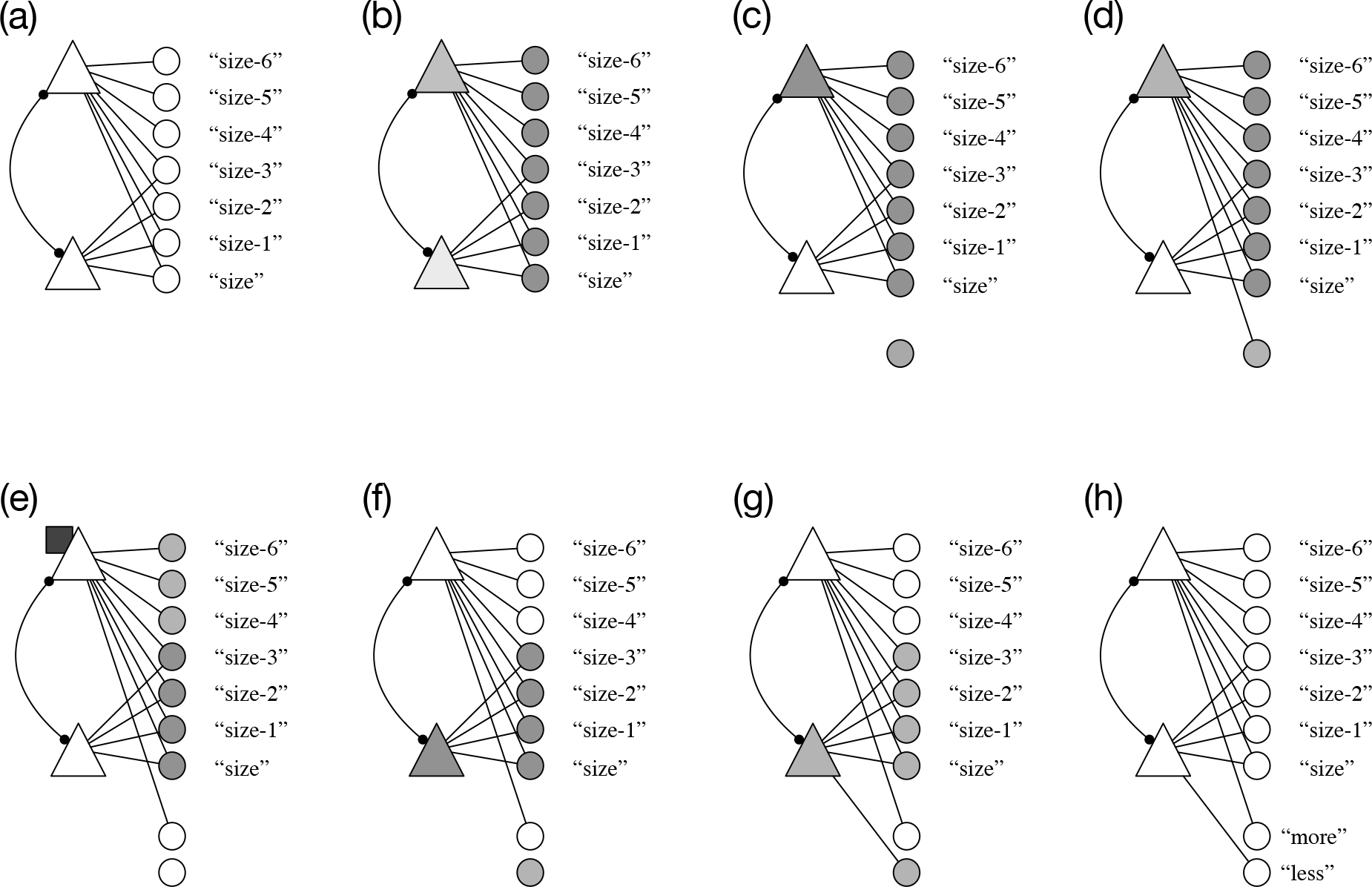
The SRM detector working on POs coding different values on a dimension. For the purposes of clarity, only the predicate POs and their semantics are depicted in this figure. Grey denotes active units. (a) Two POs coding for different sizes are in the driver. (b) The semantics coding for absolute dimensional information become active and the two POs compete via lateral inhibition to become active. (c) The unit coding for the greater value on the dimension (here size-6) becomes active first, and some semantic (dark grey circle belwo the active semantic units) responds to winning the competition between POs. (d) The PO learns a connection to the semantic that responds to winning the SRM competition. (e) The active PO unit’s inhibitor (dark square) becomes active, which inhibits the PO to inactivity. (f) The unit coding for the lesser value on the dimension (here size-3) becomes active next. (g) The active PO is connected to the semantic unit coding for losing the SRM competition. (h) The semantic unit responding to winning units in the SRM competition is an invariant response to relative “more”, while the semantic unit responding to the losing unit in the PO competition is an invariant response to relative “less”.

Similarly, when DORA attends to two representations with the same magnitude value (e.g., two POs attached to the same absolute size are present in the driver together; Figure 8a), the representations of the absolute magnitude semantics are co-activated and the PO units attached to these semantic units compete via lateral inhibition (Figure 8b). The POs will eventually settle, with both POs in a steady state of co-activation (Figure 8c). Semantic units excited by two active POs become active, and the active POs learn connections to these active semantic units by Eq. 1 (Figure 8d). These semantics act as the invariant signal for “same.”

**Figure 8.**
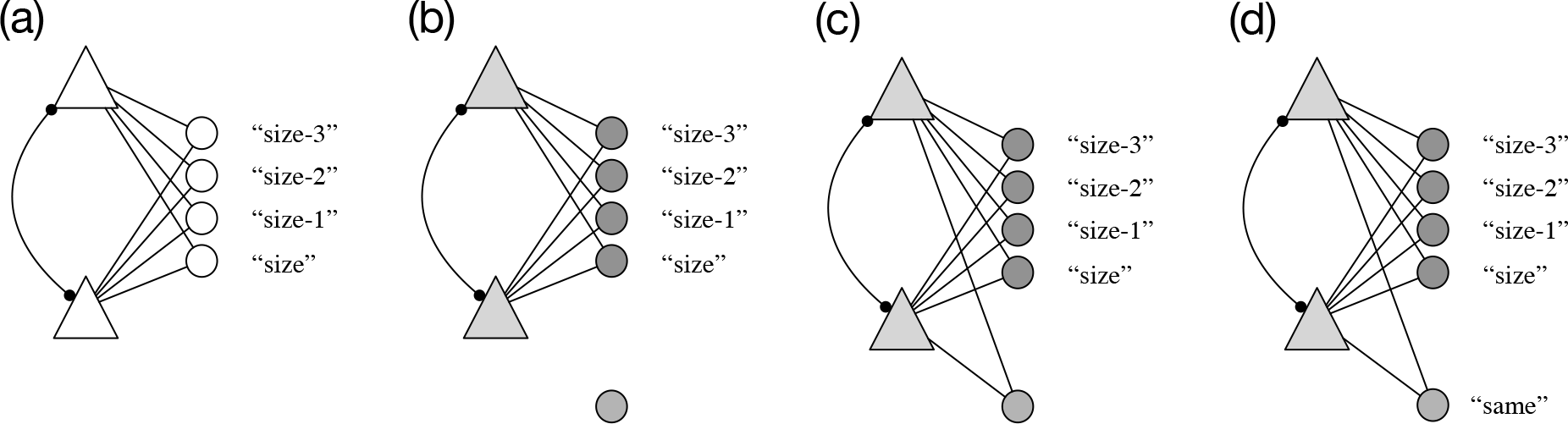
The SRM detector working on POs coding the same values on a dimension. For the purposes of clarity, only the predicate POs and their semantics are depicted in this figure. Grey denotes active units. (a) Two POs coding for the same sizes are in the driver. (b) The semantics coding for absolute dimensional information become active and the two POs compete, via lateral inhibition, to become active. (c) The PO units settle into a steady state of equivalent co-activation. (d) The PO units learn connections to the semantic unit coding for two co-active PO units during the SRM competition, or the invariant of “same”.

In short, comparing different magnitudes in a network in which magnitude information is coded by an absolute proxy (as in the human neural system), produces one of three patterns. (1) Both units settle into a state of similar co-activation—which occurs when two representations of the same magnitude are compared. (2) One unit becomes more active and forces the second unit to inactivity—which occurs when a unit codes for a greater magnitude. (3) One unit becomes active after it has been inhibited by a winning unit—which occurs when a unit codes for a lesser magnitude. Whatever units respond to these patterns naturally or through learning (see Appendix B) become implicit invariant codes for the presence of “sameness”, relative “moreness”, and relative “lessness”, respectively. What is left for the system, then, is to learn explicit representations of these semantic invariants that are not tied to any specific magnitudes and that can take other representations as arguments. This kind of learning is precisely what DORA does.

### Retrieval and mapping

DORA adopts its retrieval and mapping algorithms from LISA (see Doumas et al., 2008; Hummel & Holyoak, 1997, 2003). Retrieval from LTM and analogical mapping are highly related processes. During retrieval, propositions in the driver become active and time-share, due to time-based binding, as described above. Patterns of semantic activation generated by the driver propositions excite token units in LTM. Analogs are retrieved from LTM into the recipient using the Luce (1956) choice axiom:

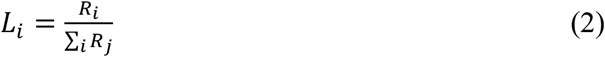

where *L*_*i*_ is the probability that analog *i* will be retrieved into working memory, *R*_*i*_ is the maximum activation of a token unit in *i* reached during the retrieval phase and *j* are all other analogs in LTM.

Analogical mapping is the processes of discovering which elements (objects, relational roles, whole propositions, etc.) of one analog correspond to which elements of another. Mapping is similar to retrieval with two important distinctions. First, activation flows from units in the driver, through the semantic units, to units in the recipient (rather than units in LTM); and second, connections (called *mapping connections*) are established between coactive units in driver and recipient. During mapping, propositions in the driver become active as described above and activate their semantic units. Units in the recipient compete via lateral inhibition to respond to the pattern of active semantic units. Units in the recipient that share the most semantic content with the active units in the driver will tend to become the most active.

At the start of mapping, mapping hypotheses are generated for each unit, *i*, in the driver, and each unit, *j*, of the same type in the recipient (e.g., mapping hypotheses are established between P units in the driver and P units in the recipient, and between RB units in the driver and RB units in the recipient). Mapping hypotheses all start at 0. The mapping process occurs in two steps: In the first step, at each time, *t*, each mapping hypothesis between driver unit *i* and recipient unit *j*, *h*_*ij*_, is updated by a simple Hebbian learning rule:

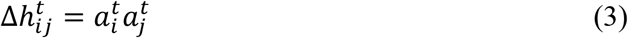

where 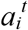 is the activation of driver unit *i* at time *t*, and 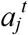 is the activation of recipient unit *j* at time *t*.

In the second step, after each *phase* set—defined as all propositions in the driver firing at least once—mapping connections are updated based on the mapping hypotheses.^2^ Before mapping connections are updated, mapping hypotheses are normalized divisively and subtractively by the equation:

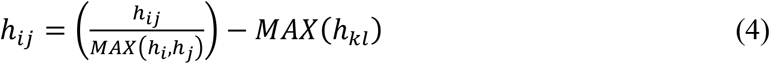

where, *h*_*ij*_ is the mapping hypothesis between units *i* and *j*, *MAX*(*h*_i_, *h*_*j*_) is the largest hypothesis involving either unit *i* or unit *j*, and *MAX*(*h*_*kl*_) is the largest mapping hypothesis where either k=*i* and *l*≠*j*, or *l*=*j* and *k*≠*i*. That is, first, each mapping hypothesis, *h*_*ij*_, is divided by the largest hypothesis involving either unit *i* or *j*, and second, the value of the largest mapping hypothesis involving either *i* or *j* (not including *h*_*ij*_ itself) is subtracted from *h*_*ij*_. The result is that mapping hypotheses are bounded between zero and one (via divisive normalization), and a one-to-one mapping constraint is established as mapping hypotheses involving the same *i* or *j* will compete with one another (via subtractive normalization; see Hummel & Holyoak, 1997). Finally, the mapping weights are updated based on their mapping hypotheses by the equation:

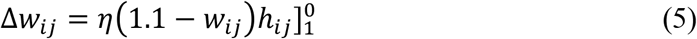

where Δ*w*_*ij*_ is the change in the mapping connection weight between driver unit *i* and recipient unit *j*, *h*_*ij*_ is the mapping hypothesis between unit *i* and unit *j*, *η* is a growth parameter, and Δ*w*_*ij*_ is truncated for values below 0 and above 1. After each phase set, mapping hypotheses are reset to 0. The mapping process continues for three phase sets.

Mapping connections represent correspondences directly between units in the driver and recipient, and allow units in the driver to activate corresponding units in the recipient directly—as opposed only to via shared semantic units. In addition, all driver units transmit a global inhibitory signal to all recipient units of the same type that is proportional to the weight of the largest mapping connection of the driver unit and the recipient unit and the mapping connection between the two units, or:

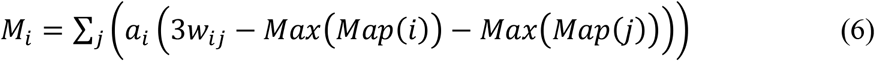

where *j* are token units of the same type as *i* in the driver (e.g., if *i* is a RB unit, *j* is all RB units in the driver), *Max*(*Map*(*i*)) is the highest of all unit *i*’s mapping connections, and *Max*(*Map*(*j*)) is the highest of all unit *j*’s mapping connections. The result is that mappings established earlier in the mapping process will constrain mappings established later. This mapping based inhibitory signal is important for DORA’s other learning routines described below.^3^

### Learning single-place predicates by comparison

DORA uses comparison to isolate shared properties of objects and to represent them as explicit structures. DORA starts with simple feature-vector representations of objects (i.e., a node connected to set of object features; Figure 9a). After mapping, corresponding elements in the driver and recipient will fire together. For example, when DORA compares a cup that is larger than something to a ball that is larger than something, the nodes representing the cup and ball will map, and will fire together (Figure 9a). Any semantic features that are shared by both compared objects (i.e., features common to both the cup and the ball) receive twice as much input and consequently become roughly twice as active as features connected to only one object (here, for example, these features might be “more”—as delivered by the SRM detection procedure, described above—and “size”; Figure 9b).

**Figure 9.**
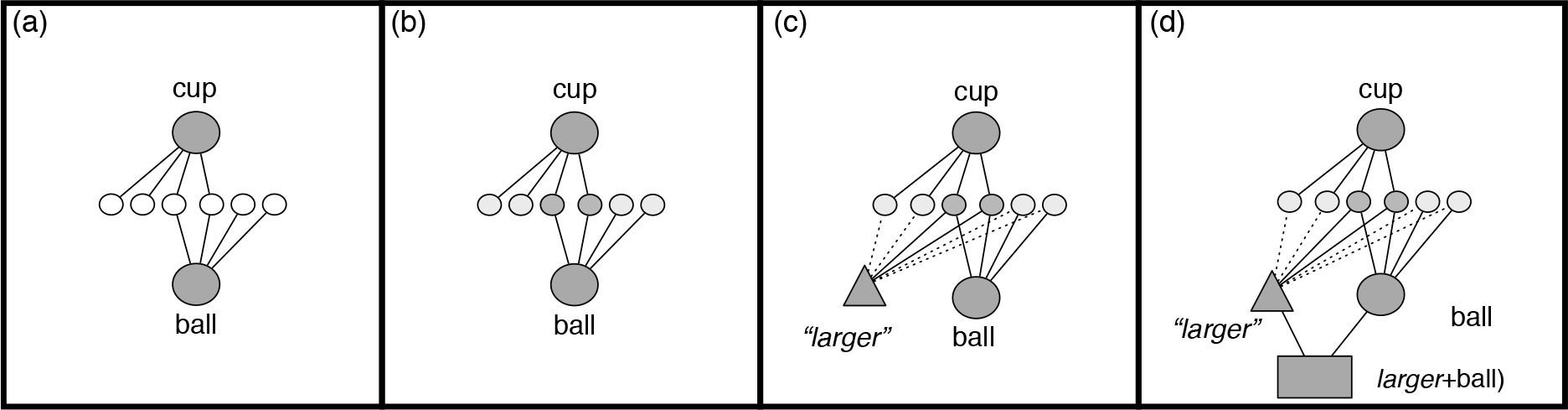
Comparison-based predication in DORA. DORA learns a representation of smaller by comparing a square that is smaller some object to a triangle smaller some object. Darker grey denotes more active units, lighter grey denotes less active units. (a) DORA compares a cup and a ball. Units representing both become active. (b) Features shared by the cup and the ball become more active than unshared features (darker grey). (c) A new unit learns connections to features in proportion to their activation (solid lines indicate stronger connection weights). The new unit codes the featural overlap of the cup and ball (i.e., the role *larger*).

DORA uses this activation based highlighting to bootstrap the explicit predication of shared properties. Whenever two solitary object units are mapped, DORA recruits a PO unit in the recipient, clamps its activation to 1.0, and learns connections between that unit and active semantics via the proportional Hebbian learning equation:

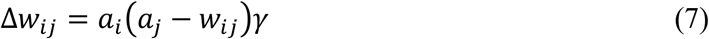

where Δ*w*_*ij*_ is the change in weight between the new PO unit, *i*, and semantic unit, *j*, *a*_*i*_ and *a*_*j*_ are the activations of *i* and *j*, respectively, and *γ* is a growth rate parameter. By Eq. 7, the weight between the recruited PO and a semantic unit will asymptote to that semantic unit’s activation. As semantics shared by the compared objects are roughly twice as active as unshared semantics, and because the strength of connections learned via Hebbian learning is a function of the units’ activations, DORA learns stronger connections between the recruited PO unit and the semantic units shared by the compared objects (Figure 9c). The recruited PO is thus an explicit representation of the featural overlap of the compared objects. In this example, DORA forms an explicit representation of “*larger*” (i.e., the features common to both the cup and ball; Figure 9d). In addition, DORA recruits a RB unit in the recipient, clamps its activation to 1.0, and learns connections between that unit and any active POs (namely, the recruited PO and the compared PO) via Hebbian learning (Fig. 9d). Importantly, because of DORA’s capacity for time-based binding (see above), the new PO will act as an explicit predicate that is dynamically bindable to fillers.

### Predicate refinement

The predicates DORA learns are likely to be initially “dirty” in that they will almost certainly contain extraneous features (e.g., any other features shared by the compared objects). Through repeated iterations of the same learning process, however, DORA forms progressively more refined representations. For example, if DORA learns a representation of *larger* by comparing a cup and a ball, this representation of *larger* might be conflated with the feature “round”. Similarly, when DORA learns a representation of *larger* by comparing a red box and a red bag, this representation of *larger* might be conflated the feature “red.” By comparing these two “dirty” representations, though, DORA can learn a refined representation of *larger*. When DORA compares the two “dirty” representations of *larger* that it has previously learned, it will map them (e.g., mapping a representation of *larger* (cup) to *larger* (box); Figure 10a). Using self-supervised learning (Doumas et al., 2008; Hummel & Holyoak, 2003) DORA recruits units in the emerging recipient that correspond to active units in the driver (Figure 10b). Because of mapping-based inhibition (see above), if any unit, *i*, in the recipient maps to a unit, *j*, in the driver, then *i* will be inhibited all other units, *k* ≠ *j*, in the driver (see Eq. 6). When units in the emerging recipient are recruited to correspond to active driver units, mapping connections are established between the corresponding units in driver and emerging recipient. As a result, when a unit, *k*, in the driver maps to no unit in the emerging recipient (or if the emerging recipient is empty), then when *k* fires, it will inhibit all emerging recipient units just as it excites no units. Any such global mapping-based inhibition is a reliable cue that nothing in the emerging recipient analog corresponds to driver unit *k*. Based on this cue, DORA recruits and activates (activation = 1.0) a unit in the emerging recipient to correspond to *k*. As in comparison-based predication, DORA learns connections between active semantics and recruited POs by Eq. 7, and between active corresponding token units (i.e., between POs and RBs, and between RBs and Ps) by simple Hebbian learning (Figure 10c). So, when DORA compares *larger* (cup) to *larger* (box), the refined representation of the predicate *larger* will have a connection weight of 1.0 to the shared semantics of the two *larger* predicates (i.e., the semantic features corresponding to “larger”), and connections to all extraneous semantics will be roughly halved (e.g., the semantics “round” and “red”). Applied iteratively, this process produces progressively more refined representations of the compared predicates—and eventually whole multi-place relations, as described below—ultimately resulting in “pure” representation of the property or relational role, free of extraneous semantics.

**Figure 10.**
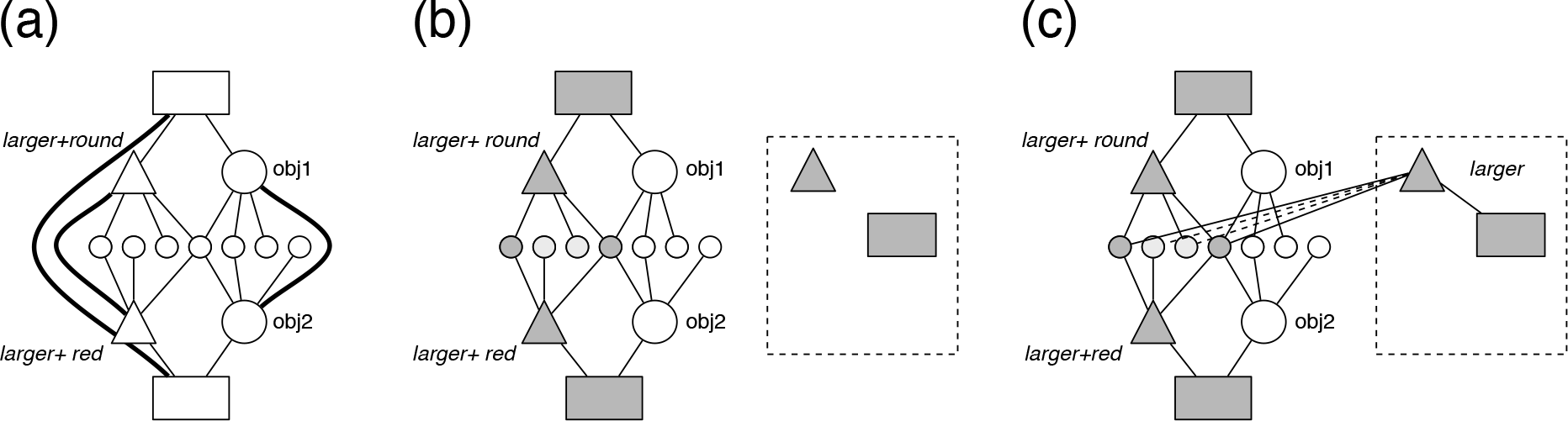
Predicate refinement in DORA. Grey denotes active units. (a) DORA maps two similar predicates. Heavy solid lines denote mapping connections. Solid lines denote connections between semantic and token units. (b) DORA recruits a PO to respond to the active driver units. (c) DORA learns connections between the new PO and RB, and between the new PO and active semantics, with stronger connections learned to more active units.

### Learning multi-place relations

DORA learns representations of multi-place relations by linking sets of constituent role-filler pairs into whole relational structures. DORA exploits the temporal dynamics of binding using time to bootstrap this process. Continuing the previous example, when DORA attends to a cup that is bigger than a ball, and a bowl that is bigger than keys, it will map the representation of *larger* (cup) to *larger* (bowl) and of *smaller* (ball) to *smaller* (keys) (Figure 11a). This process results in a distinct pattern of firing over the units composing each set of propositions: namely, the RB unit coding *larger*+cup fires out of synchrony with the RB unit coding *smaller*+ball, while the RB unit coding *larger*+bowl fires out of synchrony with the RB unit coding *smaller*+keys (Figure 11b-d). This distinct pattern emerges in the model only under two conditions: First, when the model maps sets of role-filler pairs that have already been linked into multi-place relations (i.e., after DORA has learned multi-place relations and happens to map them; e.g., when solving an analogy problem); and second, when DORA encounters multiple structurally similar sets of co-occurring role-filler pairs (i.e., when DORA encounters and then maps similar pairs of roles that are co-occurring in the world). Consequently, the pattern serves as a reliable signal that DORA exploits to bootstrap learning multi-place relations. When this diagnostic pattern emerges, DORA recruits a P unit in the recipient and clamps its activation to 1.0 (Figure 11e). DORA then learns connections between the newly recruited P unit and RB units in the recipient as they become active, via Hebbian learning. Continuing the above example, as the RB unit coding *larger*+bowl becomes active, DORA learns a connection between that unit and the newly recruited P unit (Figure 11f-g). Similarly, when the RB unit coding *smaller*+keys becomes active, DORA learns a connection between that unit and the newly recruited P unit (Figure 11h-i). The result is a P unit linking the RBs in the recipient to form a whole multi-place relational structure, *bigger* (bowl, keys) (Figure 11i).

**Figure 11.**
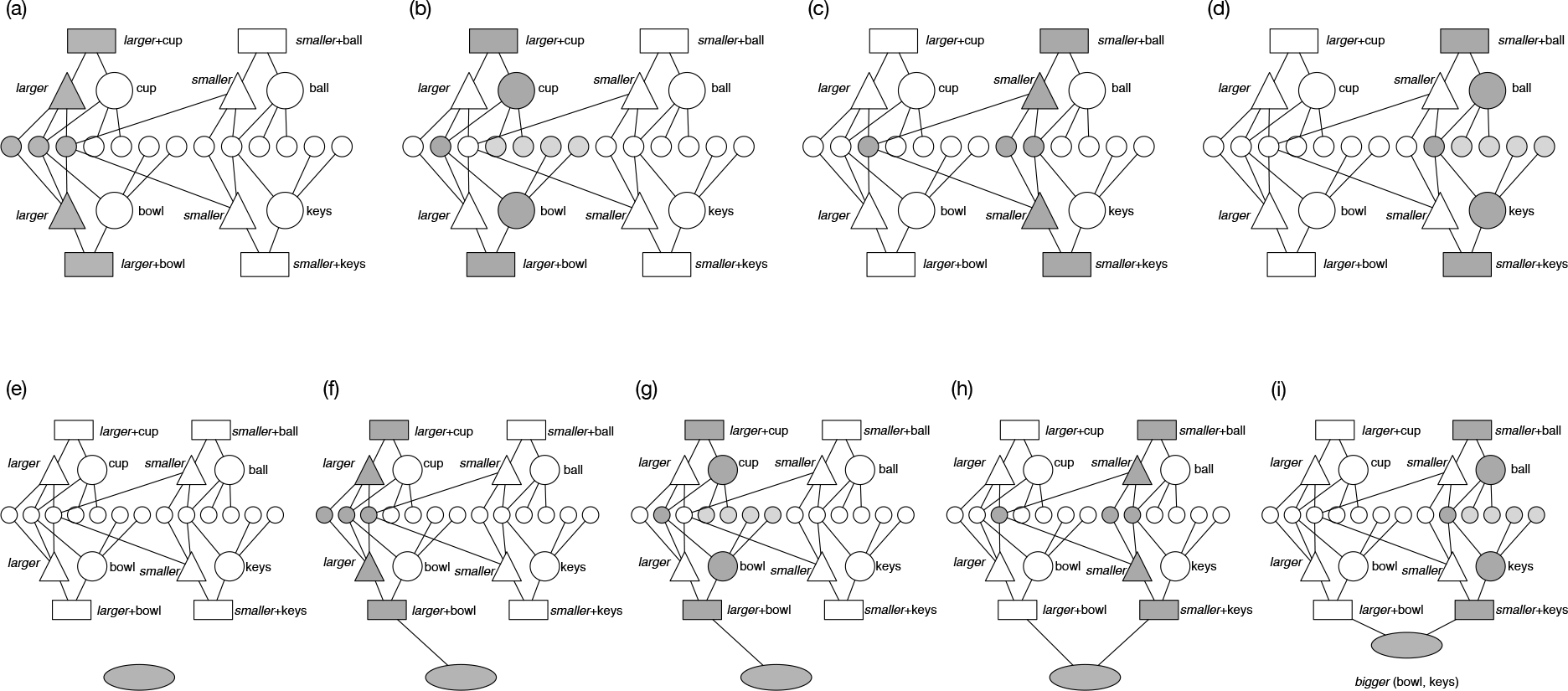
DORA learns a representation of the whole relation *bigger* (bowl, keys) by mapping *larger*(cup) to *larger*(bowl) and *smaller*(ball) *smaller*(keys). Units above the semantic units (small circles) are driver units. Units below the semantic units are recipient units. Grey denotes active units. (a) The units coding *larger* in the driver and recipient fire; (b) the units for cup and bowl fire; (c) the units for *smaller* in the driver and recipient fire; (d) the units for ball and keys fire. (e) DORA recruits a P unit in the recipient. (f-g) DORA learns a connection between the recruited P unit and the active RB unit (the RB coding for *larger*+bowl). (h-i) DORA learns a connection between the recruited P unit and the active RB unit (the RB coding for *smaller*+keys). The result is a structure coding for *bigger* (bowl, keys).

### An example of invariant SRM responding and DORA-based structure learning

Solving the problem of learning structured representations of SRM is accomplished in DORA by using DORA’s predicate learning algorithms to learn explicit structured predicate representations of the invariant signals produced by the SRM detection procedure. In brief, DORA learns explicit representations encoding absolute dimensional information, these values are compared via the SRM detection procedure, creating “valued” representations of dimensions (e.g., *more-size-9*, *less-size-4*, *same-size-6*), and through predicate refinement, connections to the specific values on the dimensions are reduced (e.g., *more-size*, *less-size*, *same-size*), and, finally, full multi-place relational representations of similarities and magnitudes are learned (e.g., *larger* (*x*, *y*), *same-size* (*a*, *b*)). We describe this process in more detail using a simplified but (hopefully) indicative example. Again, full details of the model are given in Appendix A.

Suppose that DORA starts with a set of objects each with a specific absolute size. Object1 and object2 are each size-1, object3 and object4 are each size-3, and object5 and object6 are each size-6. For the sake of simplicity, we assume that these objects are only attached to properties encoding the size dimension and that object’s specific value on that dimension. (As we show in the simulations, this assumption is completely unnecessary for the model to work, but makes the example much simpler to follow.) If DORA compares object1 and object2, it will learn a predicate coding explicitly for their semantic overlap (i.e., size-1). Similarly, if DORA compares object3 and object4, it will learn a predicate coding for size-4, and if DORA compares object5 and object6, it will learn a predicate coding for size-6.

Subsequently, when pairs of the new predicate representations are in the driver together, DORA invokes the SRM detection procedure. For example, consider the case depicted in Figure 7, where predicates coding for size-3 and size-6 are in the driver together. When the representations of the semantic encodings of specific values on the same dimension are present together (here size-3 and size-6; Figure 7a), the SRM detection procedure is invoked, and the two PO units compete to become active (Figure 7b). The unit attached the greater magnitude will ‘win’ the competition (the invariant outcome for the greater magnitude during competition between magnitudes, see above), and is connected to the semantic unit coding for winning the SRM competition—i.e., the semantic unit that serves as the invariant code for “more” (Figure 7c). The unit attached to the smaller magnitude will ‘lose’ the competition (the invariant outcome for the lesser magnitude during competition between magnitudes, see above), and is connected to the semantic unit coding for losing the SRM competition—i.e., the semantic unit that serves as the invariant code for “less” (Figure 7d). The result is a pair of predicates coding for “more”+“size-6” and “less”+“size-3.”

Similarly, when predicates coding for size-3 are in the driver together (Figure 8a), the SRM detection procedure is invoked (Figure 8b). The two PO units compete to become active, but, as both magnitudes are the same, the competition resolves with the activations of both tokens units settling at a consistent activation (Figure 8c), thus marking both values as the “same” (Figure 8d). The result is a pair of predicates coding for “same”+“size-3” and “same”+“size-3.”

After running more such comparisons, DORA will encode several predicates coding values such as “more”+“size-3,” “less”+“size-1,” and “same”+“size-6.” When DORA than compares and maps sets of these predicates coding for similar magnitude directions, but different specific values, for example mapping “more”+“size-6” to “more”+“size-3” (Figure 12), predicate refinement will produce a “value-free” representation of more-size. Specifically, when the two predicates are mapped (Figure 12a), they both activate their respective semantic units. Units shared by both the mapped predicates will get roughly twice as much input and become, therefore, roughly twice as active as any units attached to only one of the two predicates (Figure 12b). When a new unit then learns connections to active semantic units in proportion to their activation (here strong connections to the semantics coding “more” and “size,” and weaker connections to the features coding for specific values of size; Figure 12c), then DORA has learned a more effectively “value-free” representation of *more-size*(*x*). Iterative comparisons would produce progressively more “value-free” representations of *more-size*(*x*), *less-size*(*x*), and *same-size*(*x*).

**Figure 12.**
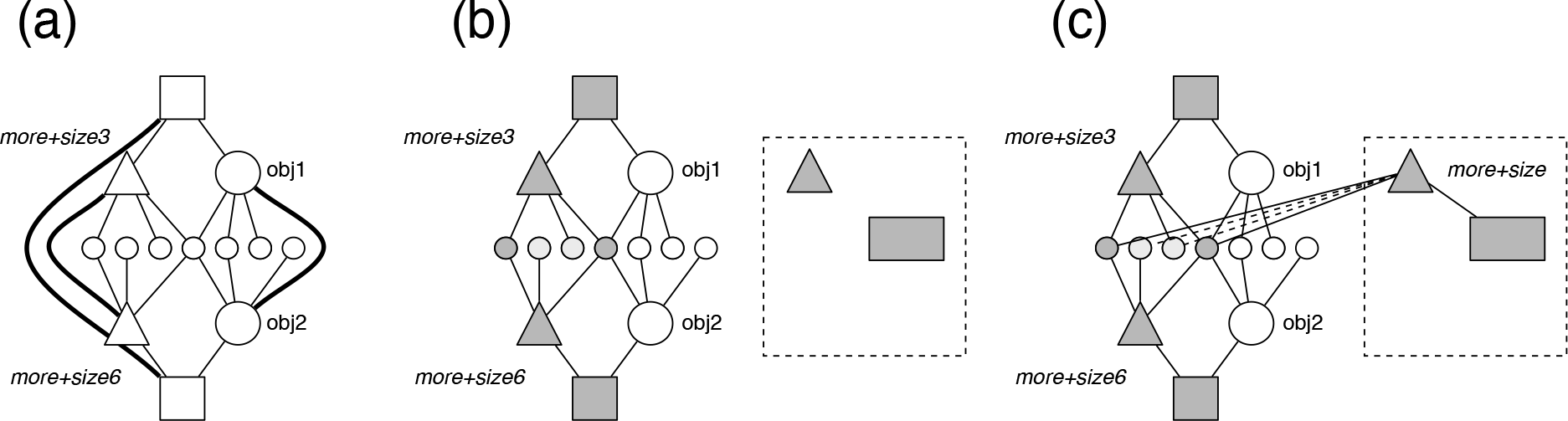
Refining representations of magnitude through predicate refinement learning. Grey denotes active units. (a) Two representations of “more” and a specific size (here size-3 and size-6) are mapped. Heavy solid lines denote mapping connections. Solid lines denote connections between semantic and token units. (b) When the two predicate tokens are coactive, any semantic units attached to both become roughly twice as active as semantic units specific to either (here semantic units coding “more” and “size” are common to both and thus the most active semantics). (c) DORA learns a representation coding for the semantic overlap of the two mapped tokens. Essentially, this new token codes for *more*+*size*, unrelated to any specific size value.

Finally, DORA learns representations of whole multi-place relational structures like *bigger* (*x*, *y*) and *same-size* (*x*, *y*). When two instances where one item is larger than another are compared, or two instances where two objects at the same size are compared, DORA will link the co-occurring role-filler pairs via a P unit to form a whole relational structure. For example, when DORA compares two instances of one object with more size and another with less size, DORA will learn a representation of the *bigger* (*x*, *y*) relational structure (Figure 13).

**Figure 13.**
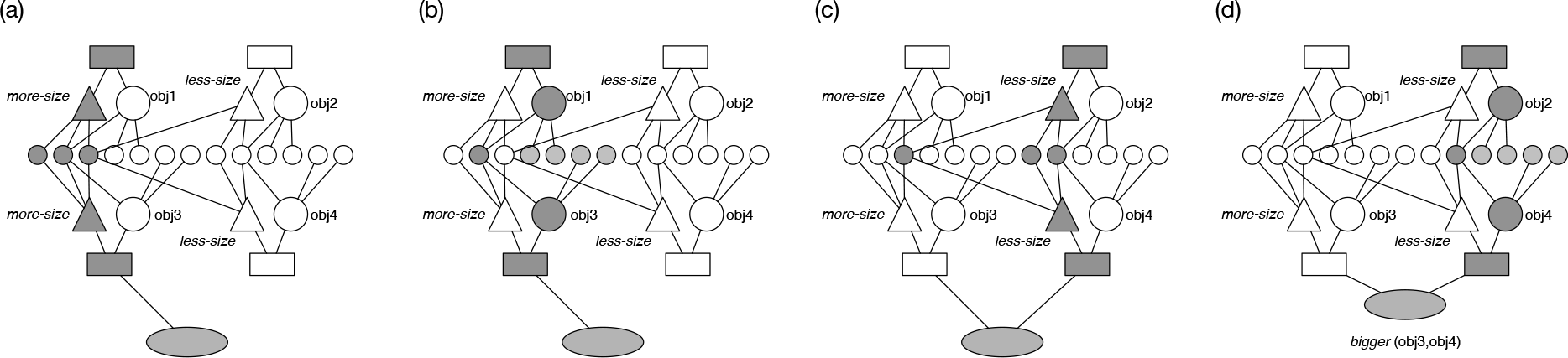
DORA learns a multi-place relational structure coding for *bigger* (obj3, obj4) via relation formation. Grey denotes active units. (a-b) DORA learns a connection between the recruited P unit and the active RB unit (the RB coding for *more-size*+obj3) when the mapped tokens coding for *more-size* fire (a), and when the mapped tokens representing obj 1 and obj3 fire (b). (c-d) DORA learns a connection between the recruited P unit and the active RB unit (the RB coding for *less-size*+obj4) when the mapped tokens coding for *less-size* fire (c), and when the mapped tokens representing obj2 and obj4 fire (d). The end result is a proposition coding for *bigger* (obj3, obj 4).

In brief, when the SRM detection procedure is integrated with DORA, it serves a purpose much like the comparator circuit in the original DORA model (see Doumas et al., 2008). However, unlike with the original comparator circuit, the current model does not require any a priori knowledge of “same” or “more” or “less” as properties. The SRM procedure we describe simply learns to respond to patterns that emerge as invariant properties of comparing magnitudes, and delivers units responding to these patterns as output. A particular pattern emerges whenever two similar values are compared, and another pattern emerges whenever items of different magnitude are compared. The model exploits these different patterns and then learns structured predicate representations of the units that come to respond to these patterns (i.e., structured representations of *same*, *more*, and *less*).

## Simulations

In this section, we describe a series of simulations aimed at demonstrating the basic properties of how humans learn structured representations of similarity and relative magnitude. Table 3 provides a set of general and specific phenomena from the literature. In the following we first provide basic proof-of-concept simulations. These simulations serve to demonstrate the various general points from Table 3. We then simulate several specific findings from the empirical literature to demonstrate the specific phenomena listed in Table 3.

Table 3. General and specific phenomena associated with human relational representation learning.

## Learning structured relational representations

*General phenomena:*

1. Humans learn concepts of similarity and relative magnitude on a number of dimensions, independent of any particular objects.
2. Humans learn structured relational representations from instances in the world that contain large amounts of extraneous information.
3. Humans learn structured representations of multiple relations simultaneously.
4. Humans learn many relational concepts with no explicit training.
5. Humans eventually learn concepts of abstract similarity and relative magnitude independent of even specific dimensions.
6. Human structured representations support analogical mapping.
7. Human relational representations support solving cross-mappings.
8. Human relational representations support mapping similar but non-identical predicates.
9. Human relational representations support mapping objects with no featural overlap, if those objects play similar roles.
10. Human relational representations overcome the n-ary restriction. *Specific phenomena:*
11. Children do not start out with the ability to reason about relative magnitude.
12. Children gradually develop the ability to reason about relative magnitude.
13. Children then use relative magnitude to discriminate between items on some dimension.
14. Children go through a relational shift.
15. Children can make analogies using representations of relative magnitude.
16. Children learn more progressively more powerful representations of relative magnitude that support more proficient relational generalization.
17. Children develop the capacity to integrate multiple relations in the service of reasoning.
18. Children’s relational representations grow more robust with learning, and allow children to overcome more excessive featural distraction.

### Proof of concept: Learning structured representations of similarity and relative magnitude from scratch

These simulations serve to demonstrate that the present model can extract implicit invariant responses of sameness, difference, moreness, and lessness from simple stimuli, and then use these responses to bootstrap learning explicit, structured, and abstract representations of these concepts. We then demonstrate that the resulting concepts meet the requirements of structured relational representations.

#### Simulation 1: Basic shapes

Humans learn structured relational representations that are explicit and independent of any particular objects (Phenomenon 1). We learn these representations from instances in the world that contain large amounts of extraneous information, and from examples that are involved in multiple relations simultaneously (e.g., a ball that is bigger than a toy car, and also lower than a book; Phenomena 2 and 3). In addition, we learn some relational concepts with no formal training (Phenomenon 4).

We used DORA to simulate learning structured representations of SRM from images of different shapes with different greyscale levels and sizes. Our goal was to test whether DORA could learn structured predicate representations of SRM relations starting with information about sets of shapes with properties of absolute values on dimensions. This simulation aims to mirror what happens when a child (or adult) learns from experience without a teacher or guide.

We started with images of basic shapes, each differing in shape, greyscale level, size, width, and height (see Figure 14 for examples of the shape stimuli; see Appendix C for details of shapes). These images were pre-processed with a feedforward neural network trained to deliver absolute shape, grey-scale level, size, width, and height information. Details of the pre-processor are given in Appendix C. When learning in the world, objects have several extraneous properties. To mirror this point, each pre-processed image was also attached to a set of 10 additional features selected randomly from a set of 1000 features. These additional features were included to act as noise, and make learning more realistic. After pre-processing, each shape was randomly paired with another to create pairs of shapes over which relational representations might be learned. So, each shape was represented as a set of semantic features connected to a PO unit, and a pair of objects consisted of two such shapes. We created 100 pairs of objects in this manner and placed them in DORA’s LTM.

**Figure 14.**
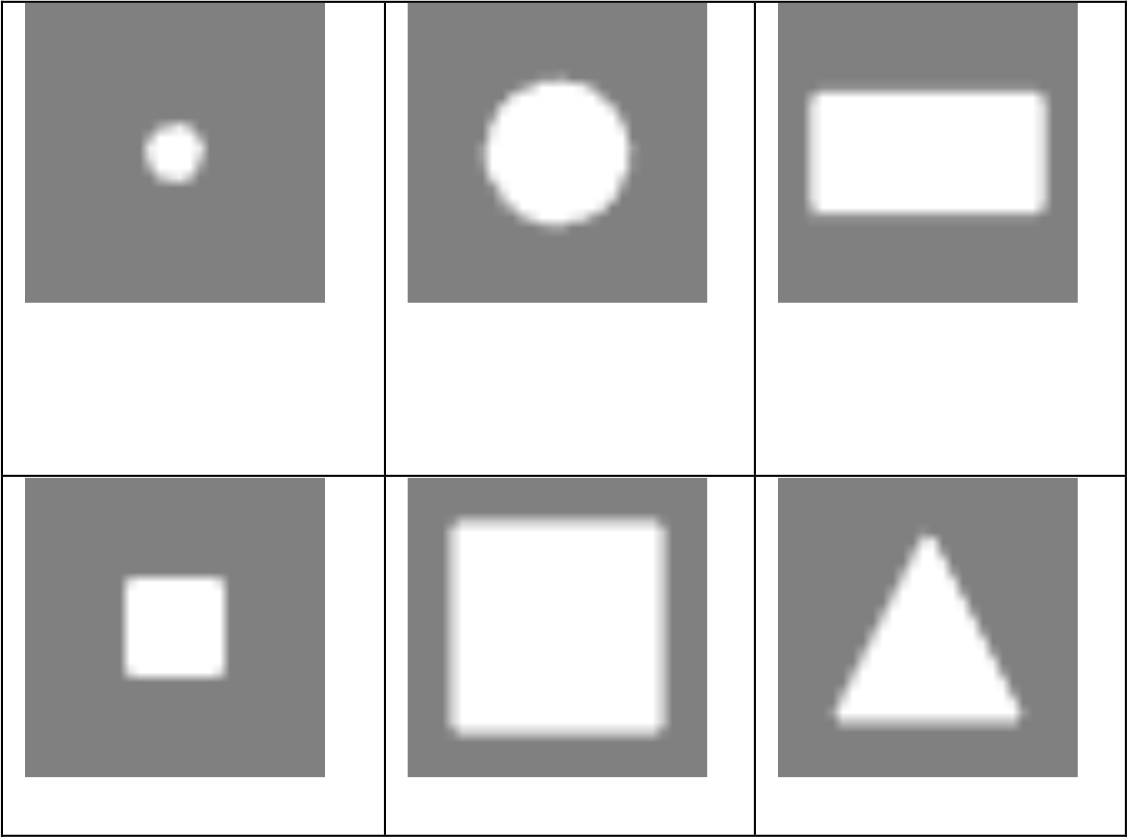
Examples of the shape stimuli used for simulation 1.

We first allowed DORA to attempt to learn from these basic representations. On each learning trial, DORA selected one pair of objects from LTM at random. DORA then ran (or attempted to run) its retrieval, mapping, SDM comparison, predication, multi-place relation learning, and refinement routines, and stored any representations that it learned in LTM. We placed one constraint on DORA’s retrieval algorithm such that more recently learned items were favoured for retrieval. Specifically, with probability .6, DORA attempted to retrieve from the last 100 analogs that it had learned. This constraint followed our assumption that items learned more recently are more salient and more likely to be available for retrieval. This simulation mimicked a child noticing something in the world, and attempting to use previous experience to understand and learn about the current experience. In short, we tested whether unguided learning from simple shape objects was sufficient for DORA to learn structured representations of SRM relations.

We defined a relational quality metric, *Q*_*i*_, for unit *i* as:

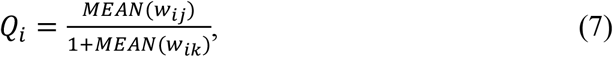

where *w*_*ij*_ are the weights of the connections between unit *i* and all relevant features *j* (i.e., those defining a specific magnitude or similarity relation or role of a said relation), and *w*_*ik*_ are the weights of the connections between unit *i* and all other semantic features, *j*. One was added to the denominator to normalize the quality measure to between 0 and 1. A higher quality denoted stronger connections to the semantics defining a specific relation relative to all other connections (i.e., a more pristine representation of the relation). We then measured the relational quality of the items in DORA’s LTM after each 100 learning trials for 2500 learning trials (where a learning trial is defined as above). Figure 15 shows the quality of the representations that DORA learned.

**Figure 15.**
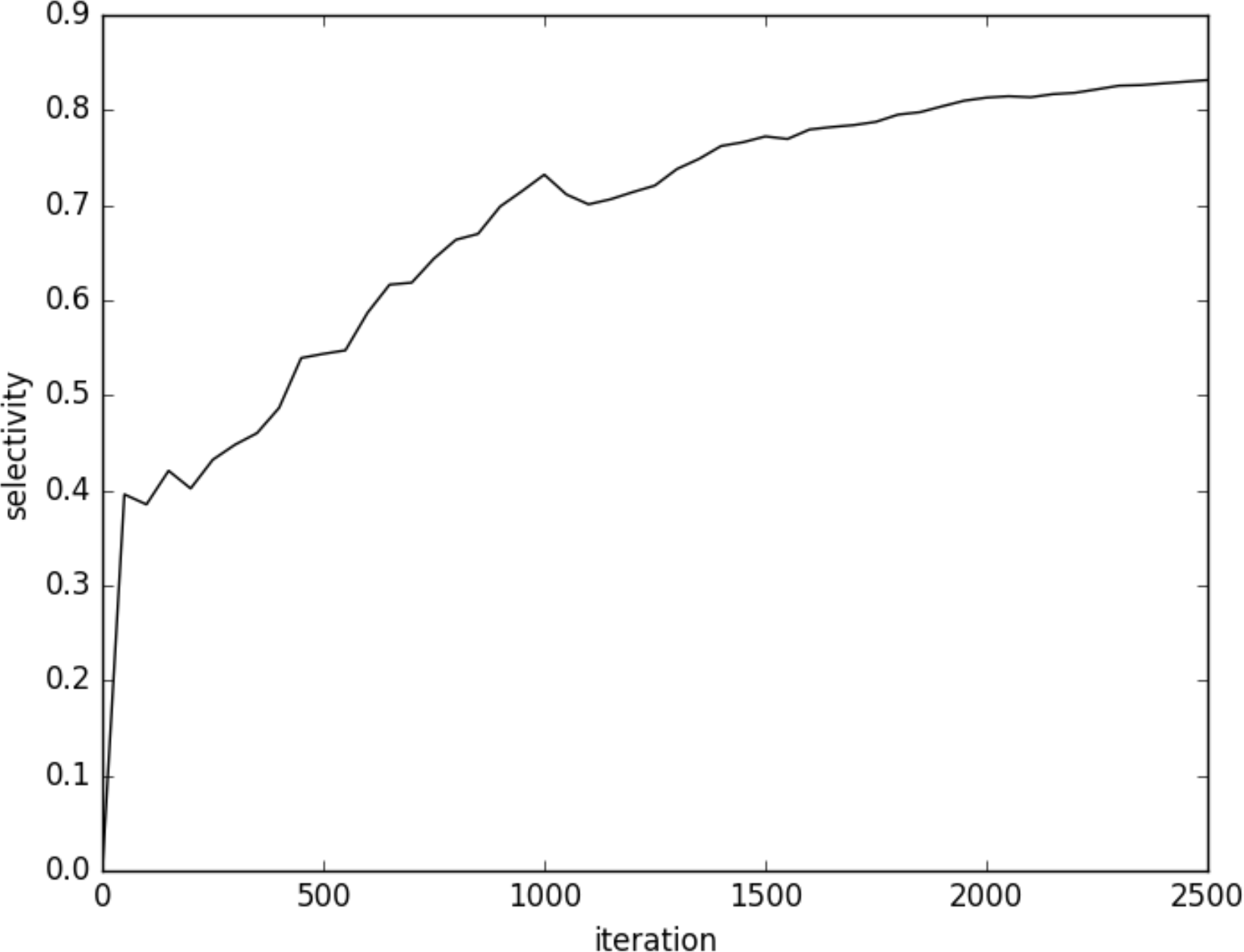
Results of DORA’s learning in simulation 1.

Clearly, these results indicate that DORA learns structured representations of relative magnitude and similarity relations from unstructured (i.e., flat feature vector) representations of objects that include only absolute values on dimensions and extraneous noise features. Figure 16 shows the number of representations of simple objects, single-place predicates, and multi-place relations in DORA’s LTM after each 100 learning trials. During learning, DORA initially learned simple single-place predicate representations encoding absolute values on dimensions. DORA then used these representations for SRM computation, refined these representations, and linked mapped sets of single-place predicates to form whole multi-place relational structures, which it then further refined through additional comparisons. Through comparison-based learning, DORA acquired representations of whole relational structures encoding SRM on all the encoded dimensions. DORA learned representations of *bigger* (where one predicate PO was connected most strongly to the semantics ‘more’ & ‘size’, the other connected to ‘less’ & ‘size’), *wider* (where one predicate PO was connected most strongly to ‘more’ & ‘width, and the other ‘less’ & ‘width’), *taller* (where one predicate PO was connected most strongly to ‘more’ & ‘height, and ‘less’ & ‘height), *same-size* (where both predicate POs were connected most strongly to ‘same’ & ‘size;), *same-width* (where both predicate POs were connected most strongly to ‘same’ & ‘width’), *same-height* (where both predicate POs were connected most strongly to ‘same’ & ‘height’), *same-colour* (where both predicate POs were connected most strongly to ‘same’ & ‘colour’), and *same-shape* (where both predicate POs were connected most strongly to ‘same’ & ‘shape’). Moreover, DORA also learned representations of *greater* (*x*,*y*) and *same* (*x*,*y*) that were independent of any particular dimensions. That is, DORA learned relations that coded strongly for only the invariant features of *more* & *less* and *same*, that were otherwise not strongly connected to any other semantic features (Phenomenon 5). These representations developed after DORA had learned some dimension specific magnitude relations (e.g., *bigger* and *wider*), and compared these representations and learned from the results. This result mirrors the development of abstract magnitude development in children (e.g., Sophian, 2007). The results indicate that DORA can learn structured representations of relative SRM relations from objects that include only absolute values on dimensions and noise.

**Figure 16.**
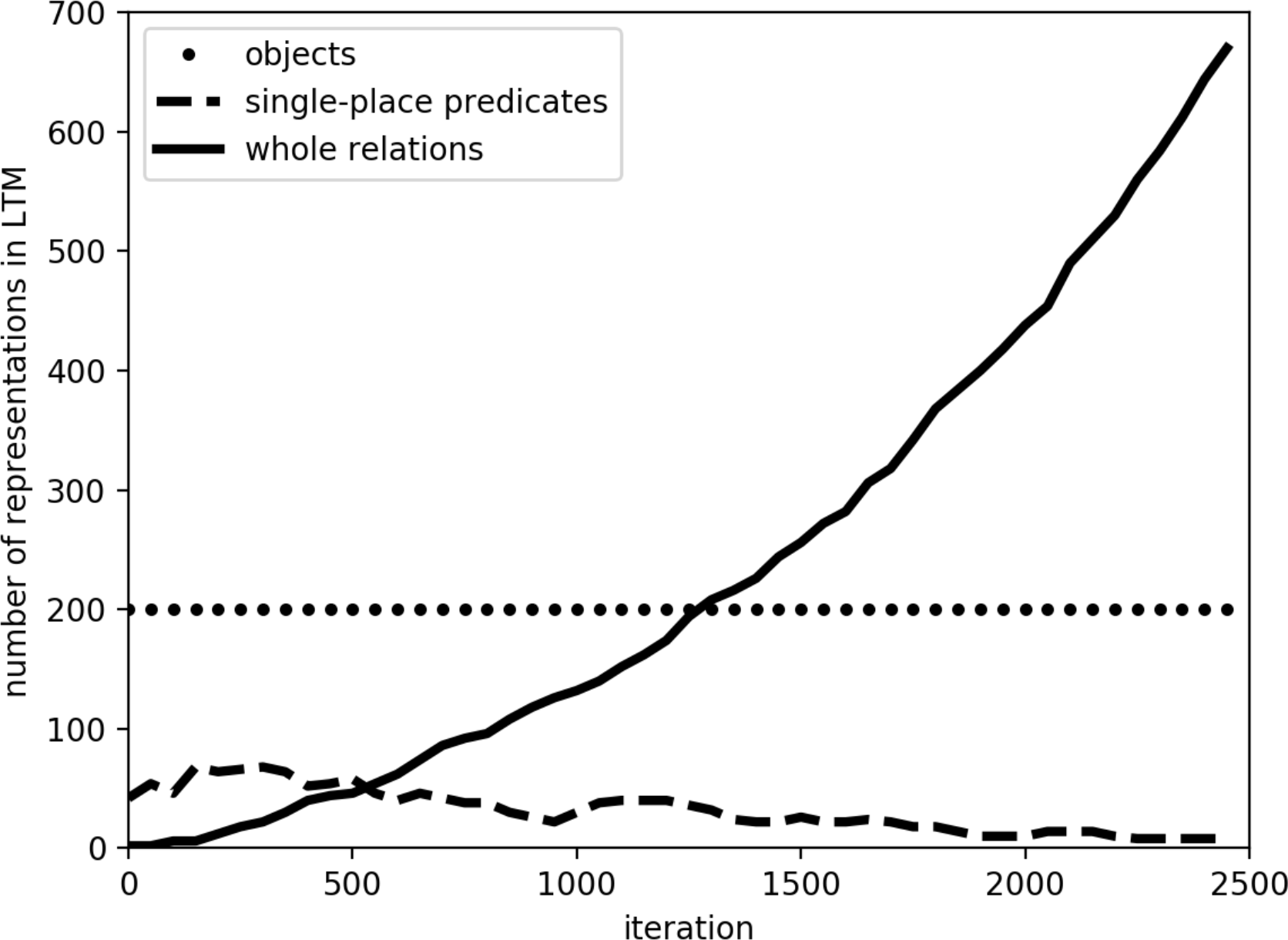
The number of representations of simple objects, single-place predicates, and whole relations in DORA’s LTM after each 100 learning trails.

A crucial question, however, is whether the representations that DORA learns behave like structured relational representations. That is, do the representations DORA learns meet the requirements of human relational representations? While almost any structured representations will support analogical mapping (Phenomenon 6) some more substantive hallmarks of relational representations (see Holyoak, 2012) are that they, (i) form the basis for solving cross mappings (Phenomenon 7); (ii) support mapping similar, but non-identical predicates (Phenomenon 8); (iii) support mapping objects with no featural overlap, if they play similar roles (Phenomenon 9); and (iv) form the basis of overcoming the n-ary restriction (Phenomenon 10).

During a cross-mapping, an object (object1) is mapped to a featurally less similar object (object2) rather than a featurally more similar object (object3) because it (object1) plays the same role as the less similar object (object2). For example, if cat1 chases mouse1 and mouse2 chases cat2, then the structural analogical mapping places cat1 into correspondence with mouse2 because both play the *chaser* role. The ability to find such a mapping is a key property of genuinely relational (i.e., as opposed to feature-based) processing (see, e.g., Gentner, 2003; Holyaok, 2012; Penn et al., 2008). Cross-mappings serve as a stringent test of the structure sensitivity of a representation as they require violating featural or statistical similarity.

We tested the relations that DORA had learned in the previous part of this simulation for their ability to support cross-mappings. We selected two of the refined relations that DORA had learned previously at random, such that both selected representations coded for the same relation (e.g., both coded for *taller*, or both coded for *same-width*). We bound the relations to new objects, creating two new propositions, P1 and P2 such that the agent of P1 was semantically identical to the patient of P2 and patient of P1 was semantically identical to the agent of P2. For example, P1 might be *taller* (square, circle) and P2 might be *taller* (circle, square). DORA then attempted to map P1 onto P2. We were interested in whether DORA would map the square in P1 onto the circle in P2 (the correct relational mapping) or simply map the square to the square and the circle to the circle. We repeated this procedure 10 times (each time with a different randomly-chosen pair of relations). In each simulation, DORA successfully mapped the square in P1 to the circle in P2 and vice-versa (because of their bindings to mapped relational roles). DORA’s success indicates that the relations it learned in the first part of this simulation satisfy the requirement of supporting cross-mapping. DORA successfully solves cross-mappings because the correspondences that emerge between matching predicates and their corresponding RBs, during asynchronous binding force relationally similar objects into correspondence. For example, consider a case when DORA attempts to map *taller* (square, circle) in the driver, and *taller* (circle, square) in the recipient. When the *more-height*+square role-binding becomes active in the driver, because of asynchronous binding, the units coding for *more-height* will become active first, followed by the units coding for square. When *more-height* is active in the driver, it will activate *more-height* and its corresponding RB, *more-height*+circle, in the recipient. When the units coding for square subsequently become active in the driver, the active *more-height*+circle RB unit in the recipient (already in correspondence with the active *more-height*+square RB unit in the driver) will activate the square unit, thus putting circle and square into correspondence, and allowing DORA to map them.

We then tested whether the relations that DORA had learned would support mapping to similar but non-identical relations (such as mapping *taller* to *greater-than*), and would support mapping objects with no semantic over-lap that play similar roles. Humans successfully map such relations (e.g., Bassok & Olseth, 1995; Gick & Holyoak, 1980, Gick & Holyoak 1983; Kubose, Holyoak, & Hummel, 2002), an ability that Hummel and Holyoak (1997, 2003) have argued depends on the semantic-richness of our relational representations. We selected two of the refined relations that DORA had learned during the previous part of this simulation, R1 and R2 (e.g, *taller*(*x*,*y*) or *wider*(*x*,*y*)). Crucially, R1 and R2 both coded for SRM across different dimensions (e.g., if R1 coded *taller*, then R2 coded *wider*). Thus, each role in R1 shared 50% of its semantics with a corresponding role in R2 (e.g., the role *more-height* has 50% of its semantics in common with the role *more-width*). To assure that no mappings would be based on object similarity, none of the objects that served as arguments of the relations had any semantic overlap at all. We repeated this process 10 times, each time with a different pair of relations from DORA’s LTM. Each time, DORA mapped the agent role of R1 to the agent role of R2 and the patient role of R1 to the patient role of R2, and, despite their lack of semantic overlap, corresponding objects always mapped to one another (because of their bindings to mapped roles).

Finally, we tested whether the representations that DORA had learned would support violating the *n-ary restriction*: the restriction that an *n*-place predicate may not map to an *m*-place predicate when *n* ≠ *m*. Almost all models of structured cognition follow the n-ary restriction (namely, those that represent propositions using traditional propositional notation and its isomorphs; see Doumas & Hummel, 2005). However, this limitation does not appear to apply to human reasoning, as evidenced by our ability to easily find correspondences between, say, *bigger* (Sam, Larry) on the one hand and *small* (Joyce) or *big* (Susan), on the other (Hummel & Holyoak, 1997).

To test DORA’s ability to violate the n-ary restriction, we randomly selected a refined relation, R1, that DORA had learned in the previous part of this simulation. We then created a single place predicate (r2) that shared 50% of its semantics with the agent role of R1 and none of its semantics with the patient role. The objects bound to the agent and patient role of R1 each shared 50% of their semantics with the object bound to r2. DORA attempted to map R1 to r2. We repeated this process 10 times, each time with a different relation from DORA’s LTM, and each time DORA successfully mapped the agent role of R1 to r2, along with their arguments. We then repeated the simulation such that r2 shared half its semantic content with the patient (rather than agent) role of R1. In 10 additional simulations, DORA successfully mapped the patient role of R1 to r2 (along with their arguments). In short, in all our simulations DORA overcame the *n*-ary restriction, mapping the single-place predicate r2 onto the most similar relational role of the multi-place relation R1.

Finally, as noted above, DORA also learned representations of *greater* (*x*,*y*) and *same* (*x*,*y*) that were independent of any particular dimensions (i.e., relations that coded strongly for only the invariant features of *more* & *less* and *same*, that were otherwise not strongly connected to any other semantic features). Importantly, these representations also met all the requirements of structured relational representations. We ran the exact same tests for crossmapping, mapping arguments with no semantic overlap based on shared roles, and violating the n-ary restriction that are described above, but using the representations of abstract magnitude (i.e., *greater* (*x*,*y*)) that DORA had learned during training. Just as with DORA’s dimensional SRM representations, DORA’s more abstract SRM representations successfully performed a cross-mapping in 10 out of 10 simulations, mapped arguments with no semantic overlap based only on shared roles in 10 out of 10 simulations, and overcame the n-ary restriction in 10 out of 10 simulations.

#### Simulation 2: Gabor patches

Simulation 2 was identical to simulation 1, with the exception that we used Gabor patches rather than simple shapes (see Figure 17 for examples of the Gabor patches used; see Appendix C for details of Gabor patches). The purpose of this simulation was to replicate simulation 1 with different starting stimuli. Simulation 2 progressed just like simulation 1. The generated Gabor patches were preprocessed using the feedforward network described in Appendix C to deliver absolute frequency, orientation, and bar width information, and extraneous noise features were added. The representations were then paired as described above, and DORA learned from the pairs for 2500 learning trials. Figure 18 shows the qualitys of the representations that DORA learned. Just as in simulation 1, DORA successfully learned representations of *greater-frequency* (one predicate PO connected most strongly to the semantics ‘more’ & ‘frequency, the other connected to ‘less’ & ‘frequency), *more-misoriented* (one predicate PO connected most strongly to ‘more’ & ‘orientation’, and the other to ‘less’ & ‘orientation), *wider* (one predicate PO connected most strongly to ‘more’ & ‘width’, and the other ‘less’ & ‘width), *same-frequency* (both predicate POs connected most strongly to ‘same’ & ‘frequency;), *same-orientation* (both predicate POs connected most strongly to ‘same’ & ‘orientation), and *same-width* (both predicate POs connected most strongly to ‘same’ & ‘width’). Once again, the results indicate that DORA learns structured representations of relative SDM relations from objects that include only absolute values on dimensions and noise. In addition, just as in simulation 1, DORA also learned representations of *greater* (*x*,*y*) and *same* (*x*,*y*) that were independent of any particular dimensions. That is, DORA learned relations that coded strongly for only the invariant features of *more* & *less* and *same*, that were otherwise not strongly connected to any other semantic features.

**Figure 17.**
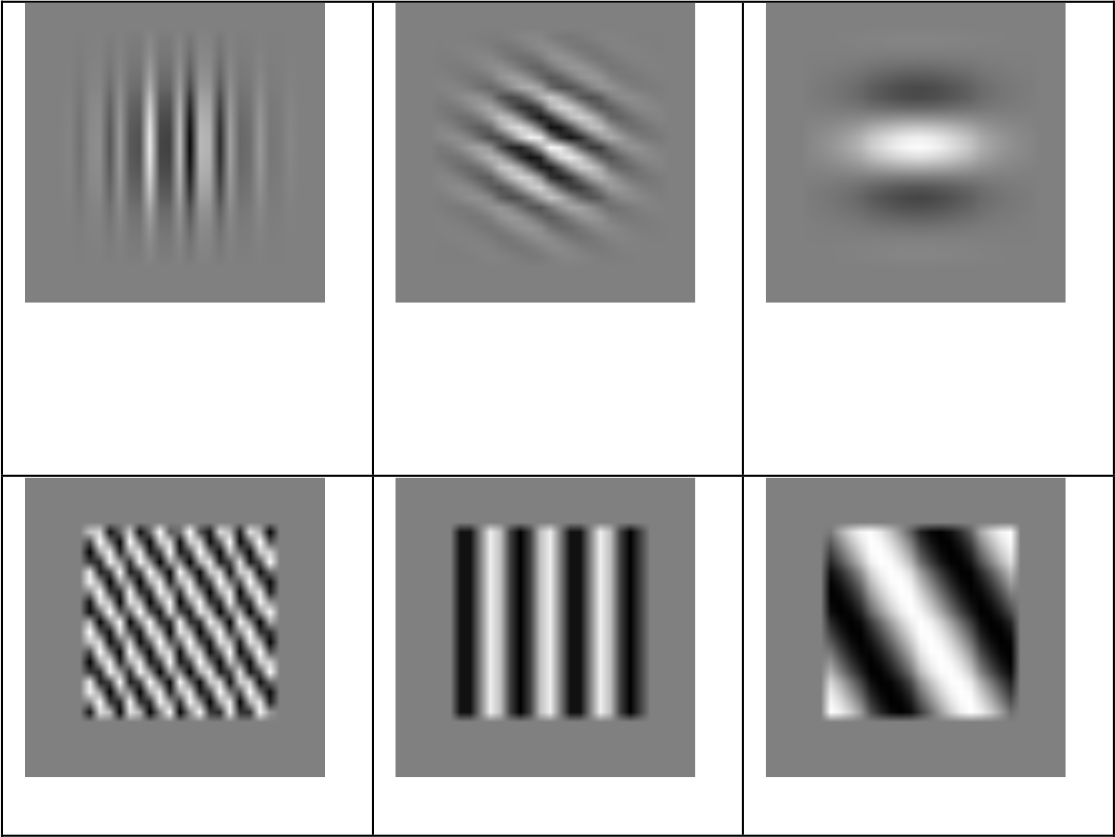
Examples of the Gabor patches used for simulation 2.

**Figure 18.**
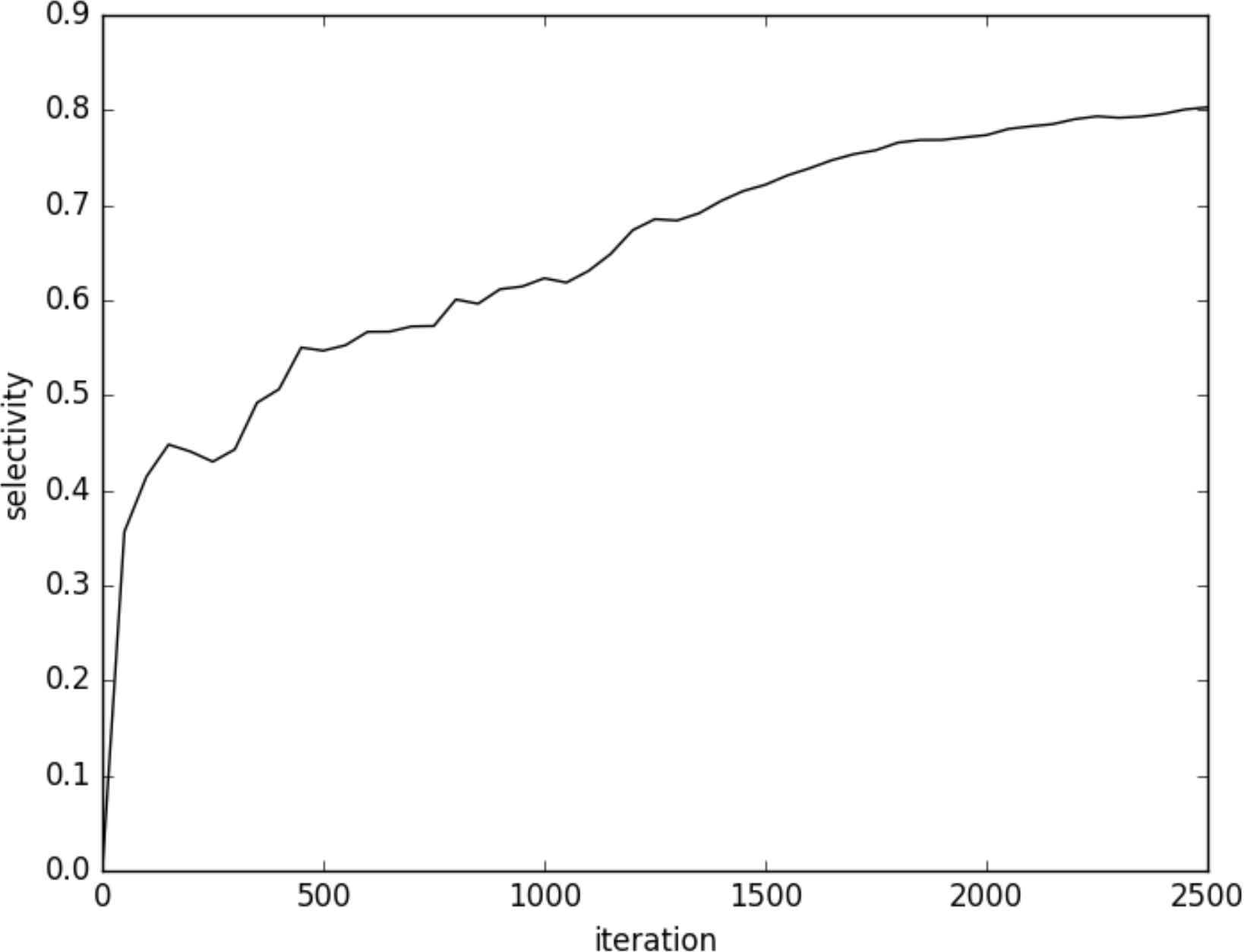
Results of DORA’s learning in simulation 2.

Just as in simulation 1, in simulation 2 we tested the representations that DORA had learned in the first part of the simulation for their capacity to support solving cross mappings, mapping similar, but non-identical predicates, mapping objects with no semantic overlap based on shared roles, and overcoming the n-ary restriction. We conducted the same tests of these capacities described in simulation 1 above. Just as in simulation 1, the representations that DORA learned in the first part of simulation 2 produced unanimous success on tests of cross-mapping, mapping similar but non-identical relations, mapping objects with no semantic overlap based on shared roles, and violation of the n-ary restriction.

In sum, simulations 1 and 2 provide compelling evidence that DORA augmented with SDM detection provides an answer to how structured representations of relative magnitude and similarity can be learned from experience, and successfully accounts for Phenomena 1-10 from Table 3, with very minimal assumptions about innate representations.

### Simulations of specific cognitive phenomena

#### Simulation 3: The development of human-like representations of relative magnitude

Children develop the capacity to reason about SRM. While children do not start out with the ability to reason about SRM (Phenomenon 11; e.g., Smith, 1989), they gradually develop the ability to reason about SRM on a variety of dimensions (Phenomenon 12; e.g., Smith, 1984). Children then use SRM to discriminate between items on some dimension (Phenomenon 13; e.g., Smith & Sera, 1992).

The development of children’s ability to reason about basic magnitudes is well demonstrated in a classic study by Nelson and Benedict (1974). In their experiment, 33 children aged three to six years-old were given a simple identification task. An experimenter presented the child with two pictures of similar objects that differed on some dimensions. The experimenter then asked the child to identify the object with a greater or lesser value on some dimension. For example, the child might be shown pictures of two fences that differed in their height, their size, and their colour, and then asked which of the two fences was taller or shorter. The developmental trajectory was clear: Children between 3-years-10-months and 4-years-4 months (mean age ~48 months) made errors on 34% of trials, children between 4-years-7-months and 5-years-5-months (mean age ~60 months) made errors on 18% of trials, and children aged 5-years-6-months and 6-years-6-months (mean age ~73 months) made errors on only 5% of trials. In short, as children got older, they developed a mastery of simple magnitude comparisons on a range of dimensions.

We simulated the results of Nelson and Benedict (1974) in two parts. In the first part, we allowed DORA to learn representations of relative magnitudes on a range of dimensions using the same procedure described in simulations 1 and 2. DORA started with representations of pairs of objects attached to features describing the object, including absolute values on several dimensions. We then ran DORA’s comparison-based learning routines for 2500 learning iterations. On each learning trial, DORA selected one pair of objects from LTM at random. DORA then ran (or attempted to run) its retrieval, mapping, SDM detection, predication, multi-place relation learning, and refinement routines, and stored any representations that it learned in LTM. The results of this part of the simulation were the same as the results of the simulations described above: DORA initially learned representations of single-place predicates describing absolute dimensional values, used these for SRM computation, learned multi-place relations by linking mapped sets of single-place predicates, and refined these representations through further comparisons.

In the second part of the simulation we simulated Nelson and Benedict’s experiment. To simulate children of different ages we stopped DORA at different points during learning and used the representations that it had learned to that point (i.e., the state of DORA’s LTM) to perform the magnitude reasoning task.

To simulate each trial, we created two objects attached to a set of semantic features. These features included nine random features selected from the pool of 1000 features, and semantic features describing height, width, and size (e.g., one semantic unit describing “size” and other semantics describing absolute size—e.g., nine semantics coding that together encoded “size-9”). We then randomly selected a dimension to serve as the question dimension for that trial (i.e., the dimension on which the question would be based), and sampled at random a representation from DORA’s LTM that was strongly connected to that dimension (with a weight of .95 or higher). If the sampled item was a relation or a single place predicate, we applied it to the objects. For example, if key dimension was size, and we sampled a representation of the relation *bigger* (*x*,*y*), we applied that representation to the objects, binding the larger object to the *larger* role and the smaller object to the *smaller* role. If the we sampled a representation of a single-place predicate *more-size* (*x*), then we bound that predicate to the larger object. Finally, we placed the representations of the two sample objects in the recipient, and a representation of a PO attached to the semantics describing the specific task in the driver. For instance, if the task was to point to the smaller item, the driver PO was attached to the semantics for “less” and “size”. We then fired the PO in the driver, and DORA attempted to map that PO to an item in the recipient. If DORA mapped the driver representation to an element in the recipient, the mapped item was taken as DORA’s response on the task. If DORA failed to find a mapping after 3 phase-sets, an item was chosen from the recipient at random and taken as DORA’s response for that trial (implying that DORA was guessing on that trial). The probability of guessing the correct item by chance was 0.5.

To simulate 4 year-olds, we used the representations in DORA’s LTM after 1300 training trials, to simulate 5 year-olds we used the representations in DORA’s LTM after 1850 training trials, and to simulate 6 year-olds we used the representations in DORA’s LTM after 2300 training trials. We ran 100 simulations with 20 trials at each age level. The results of our simulation and the original results of Nelson and Benedict are presented in Fig. 19.

**Figure 19.**
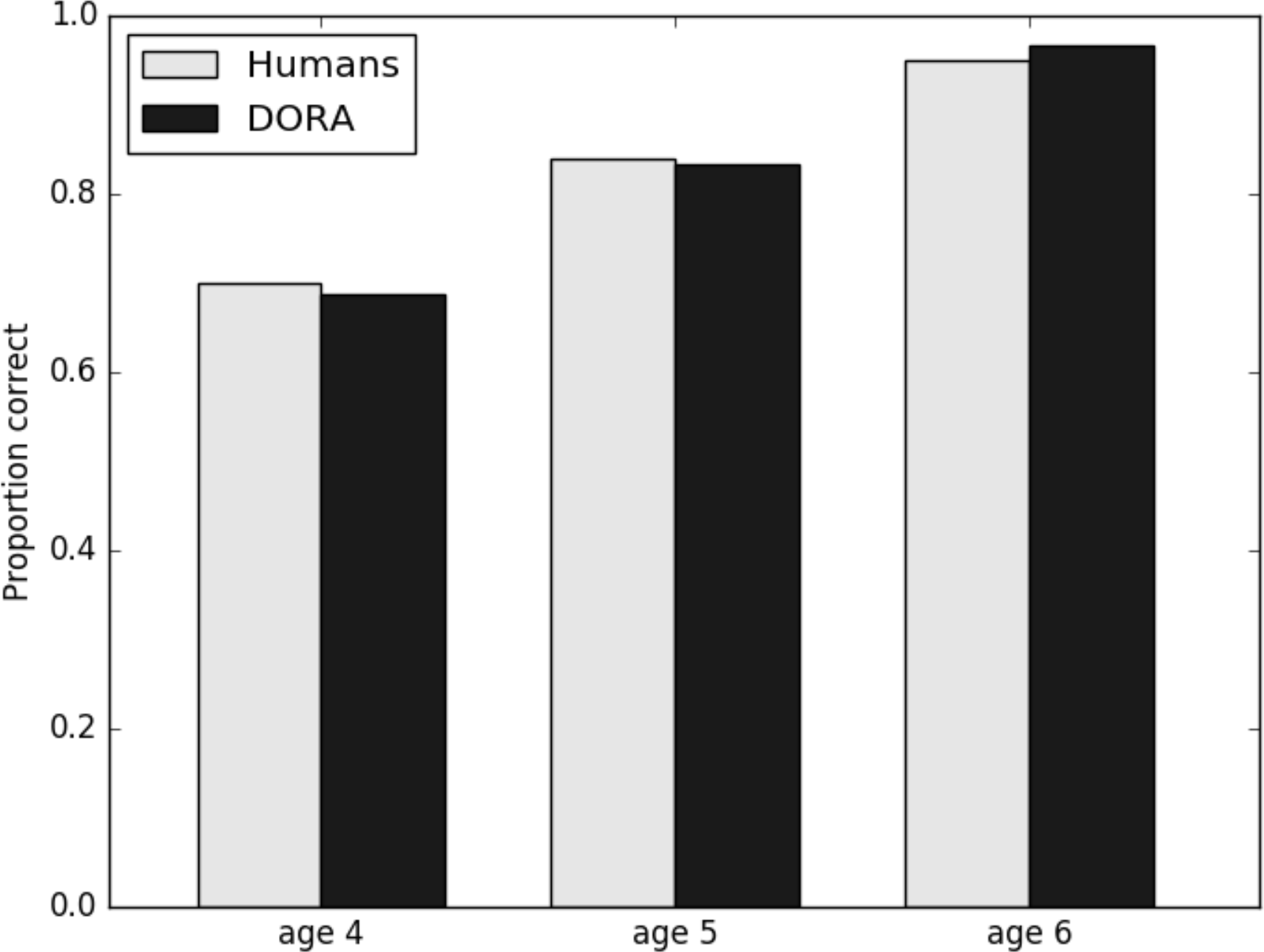
Results of simulation of Nelson and Benedict (1974).

As can be seen in Fig. 19, the qualitative fit between DORA’s performance and the performance of the children in Nelson and Benedict (1974) is very close. Just like the children in the original study, DORA is initially better than chance, but still quite error prone early during learning, but gradually comes to learn representations that support very successful classification of dimensional magnitudes. These simulation results provide evidence that the trajectory of the development of DORA’s representations of relative dimensional magnitude mirrors that of humans, and also that the representations that DORA learns support the same kinds of processing that humans come to excel in.

#### Simulation 4: Relative magnitude representations and the relational shift

One of the key findings from work on the development of analogical reasoning in children, is that children go through a relational shift (Phenomenon 14; see, e.g., Gentner, 2003). The relational shift describes a qualitative change in children’s reasoning wherein they progress from making analogies based on the literal features of things, to making analogies based on the relations that objects are involved in (Phenomenon 15; e.g., Gentner et al., 1995). With development, children learn progressively more powerful representations of SRM relations that support more proficient relational generalization (Phenomenon 16; Smith, 1984). In addition, children develop the capacity to integrate multiple relations in the service of reasoning (Phenomenon 17; e.g., Richland, Morrison, & Holyoak, 2006), and their relational representations grow more robust with learning, and allow them to overcome ever more excessive featural distraction (Phenomenon 18; e.g., Halford & Wilson, 1980; Richland, Morrison, & Holyoak, 2006).

One of the classic examples of the relational shift and the associated phenomena is given in Rattermann and Gentner (1998). In their experiment, Rattermann and Gentner had 3-, 4-, and 5-year-old children participate in a relational matching task. Children were presented with two arrays, one for the child and one for the experimenter. Each array consisted of three items that varied on some relative dimension. For example, the three items in each array might increase in size from left to right, or decrease in width from left to right. The dimensional relation in both presented arrays was the same (e.g., if the items in one array increased in size from left to right, the items in the other array also increased in size from left to right). The items in each array were either sparse (simple shapes of the same colour) or rich (different objects of different colours). The child watched the experimenter hide a sticker under one of the items in the experimenter’s array. The child was then tasked to look for a sticker under the item from the child’s array that matched the item selected by the experimenter. The correct item was always the relational match—e.g., if the experimenter hid a sticker under the largest item, the sticker was under the largest item in the child’s array. Critically, at least one item from the child’s array matched one of the items in the experimenter’s array exactly except for its relation to the other items in its array. To illustrate, if the experimenter might have an array with three squares increasing in size from left to right (Figure 20a). The child might have an array of three squares also increasing in size from left to right, but with the smallest item in the child’s array identical in all featural properties to the middle item in the experimenter’s array (Figure 20b). Thus, each trial created a crossmapping situation, where the relational choice (same relative size in the triad) was at odds with the featural choice (exact object match). The child was rewarded with the sticker if she chose correctly.

**Figure 20.**
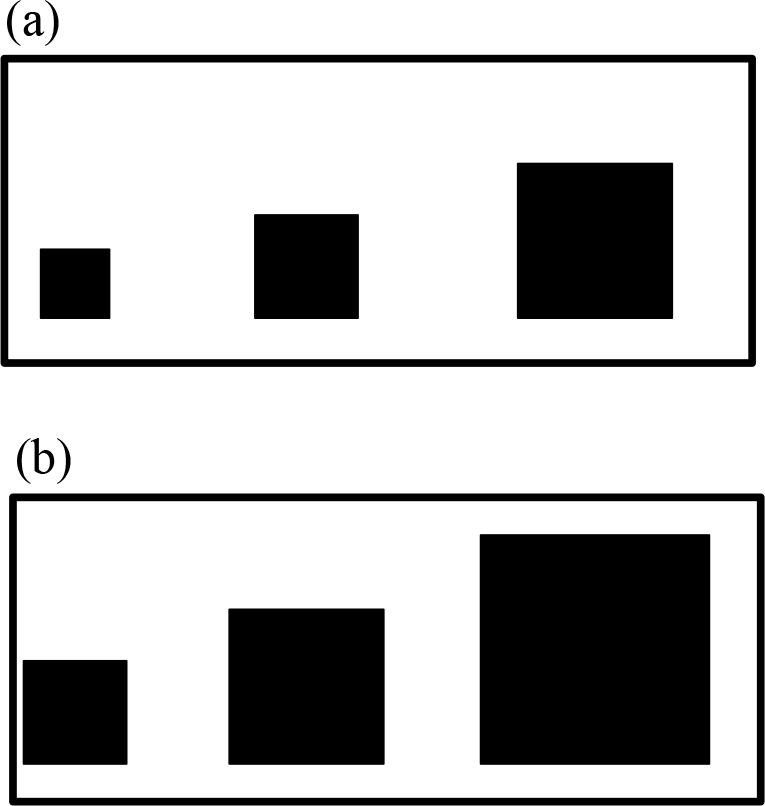
A recreated example of the stimuli used in Rattermann and Gentner (1998).

Rattermann and Gentner found a clear indication of a relational shift. Children between 3 and 4-years-old were very drawn by featural matches, and had trouble systematically making relational matches (making relational matches 32% of the time in the rich condition and 54% of the time in the sparse condition). Children between 4 and 5-years-old were quite good at making relational matches with sparse objects—making relational matches 62% of the time—but still had trouble with rich objects when featural matches were more salient—making relational matches 38% of the time. Children between 5 and 6-years-old were quite good at making relational matches in both the rich and the sparse conditions, with the rich condition providing more trouble than the sparse condition—making relational matches 68% for rich and 95% of the time for sparse stimuli.

Just as in the simulation above, we simulated the results of Rattermann and Gentner (1998) in two parts. In the first part, we allowed DORA to learn representations of relative magnitudes on a number of dimensions using the same procedure described in simulations 1 and 2. DORA started with representations of objects attached to features describing the object, including absolute values on several dimensions. We then ran DORA’s comparison-based learning routines for 2500 learning iterations. On each learning trial, DORA selected one pair of objects from LTM at random. DORA then ran (or attempted to run) its retrieval, mapping, SDM comparison, predication, multi-place relation learning, and refinement routines, and stored any representations that it learned in LTM. The results of this part of the simulation were the same as the results of the simulations described above: DORA initially learned representations of single-place predicates describing absolute dimensional values, used these for SRM detection, learned multi-place relations by linking mapped sets of single-place predicates, and refined these representations through further comparisons.

In the second part of the simulation, we simulated Rattermann & Gentner’s (1998) experiment. To simulate children of different ages we stopped DORA at different points during learning and used the representations that it had learned to that point (i.e., the state of DORA’s LTM) to perform the relational matching task.

To simulate each trial, we created two arrays of three objects. For the sparse trials, each object was attached to a number of semantic feature units: some features describing size (“size” and absolute size), some features describing height (“height” and absolute height), some features describing width (“width” and absolute width), some features describing colour, two features describing shape, one feature describing position in the array (“left”, “center”, “right”), and four features chosen at random from a pool of 1000. The identical objects from both arrays matched on all semantic features. For the rich trials, each object was attached to a number of semantic features: some features describing size (“size” and absolute size), some features describing height (“height” and absolute height), some features describing width (“width” and absolute width), two features describing colour, two features describing shape, four features describing object kind (e.g, “shoe”, “train”, “bucket”), one feature describing position in the array (“left”, “center”, “right”), and 32 features chosen at random from a pool of 1000. The identical objects from both arrays matched on all semantic features. We ordered the objects in both arrays according to some relation (e.g., increasing size, decreasing width), and then sampled four representations from DORA’s LTM that were strongly connected to that dimension (with a weight of .95 or higher). We applied two of the sampled representations to each of the two arrays. If the sampled representation was a relation or a single place predicate, we applied it to the objects. For example, if the key dimension was size, and we sampled a representation of the relation *bigger* (*x*,*y*), we applied that representation to the objects, binding the larger object to the *larger* role and the smaller object to the *smaller* role. If the we sampled a representation of a single-place predicate like *more-size* (*x*), then we bound that predicate to the larger object. As each array consisted of two instances of the key relation (e.g., object1 is *bigger* than object2, and object2 is *bigger* than object3), we applied one of the two sampled items to one of the relations in the array, chosen at random, and the other sampled item to the other relation in the array. For example, if object1 is bigger than object2, and object2 is bigger than object3, and we sampled two representations of *bigger* (*x*,*y*) from DORA’s LTM, then we applied one of the two *bigger* (*x*,*y*) relations to object1 and object2 (i.e., *bigger* (object1, object2)), and the other to object2 and object3 (i.e., *bigger* (object2, object3)).

Finally, we placed the representation of the child’s array in the driver, and the experimenter’s array in the recipient, and allowed DORA to attempt to map the driver and recipient representations. An item from the recipient was chosen at random as the “sticker” item (i.e., the item under which the sticker was hidden). The capacity to ignore features is a function of the salience of those features (Goldstone & Son, 2005), and so richer objects with more features are harder to ignore (Tversky, 1977). To simulate the effect of the rich vs. the sparse stimuli, on each trial we allowed DORA to make a simple similarity comparison before relational processing started. We randomly selected one of the items in the driver, and computed the similarity between that item and the “sticker” item in the recipient. If the computed similarity was above .8, then DORA learned a mapping connection between the two items. Finally, DORA attempted to map the items in the driver to the items in the recipient. If DORA mapped the driver representation to the “sticker” item in the recipient, the mapped item was taken as DORA’s response on the task. If DORA failed to find a mapping after 3 phase-sets, an item was chosen from the recipient at random and taken as DORA’s response for that trial (implying that DORA was guessing on that trial). The probability of guessing the correct item by chance was 0.33.

To simulate 3 year-olds we used the representations in DORA’s LTM after 850 training trials, to simulate 4 year-olds we used the representations in DORA’s LTM after 1550 training trials, and to simulate 5 year-olds we used the representations in DORA’s LTM after 2050 training trials.

We ran 100 simulations each consisting of 20 trials at each age level. The results of the simulation as well as those from the original Rattermann and Gentner experiment with both sparse and rich trials are presented in Fig. 21a and b respectively.

**Figure 21.**
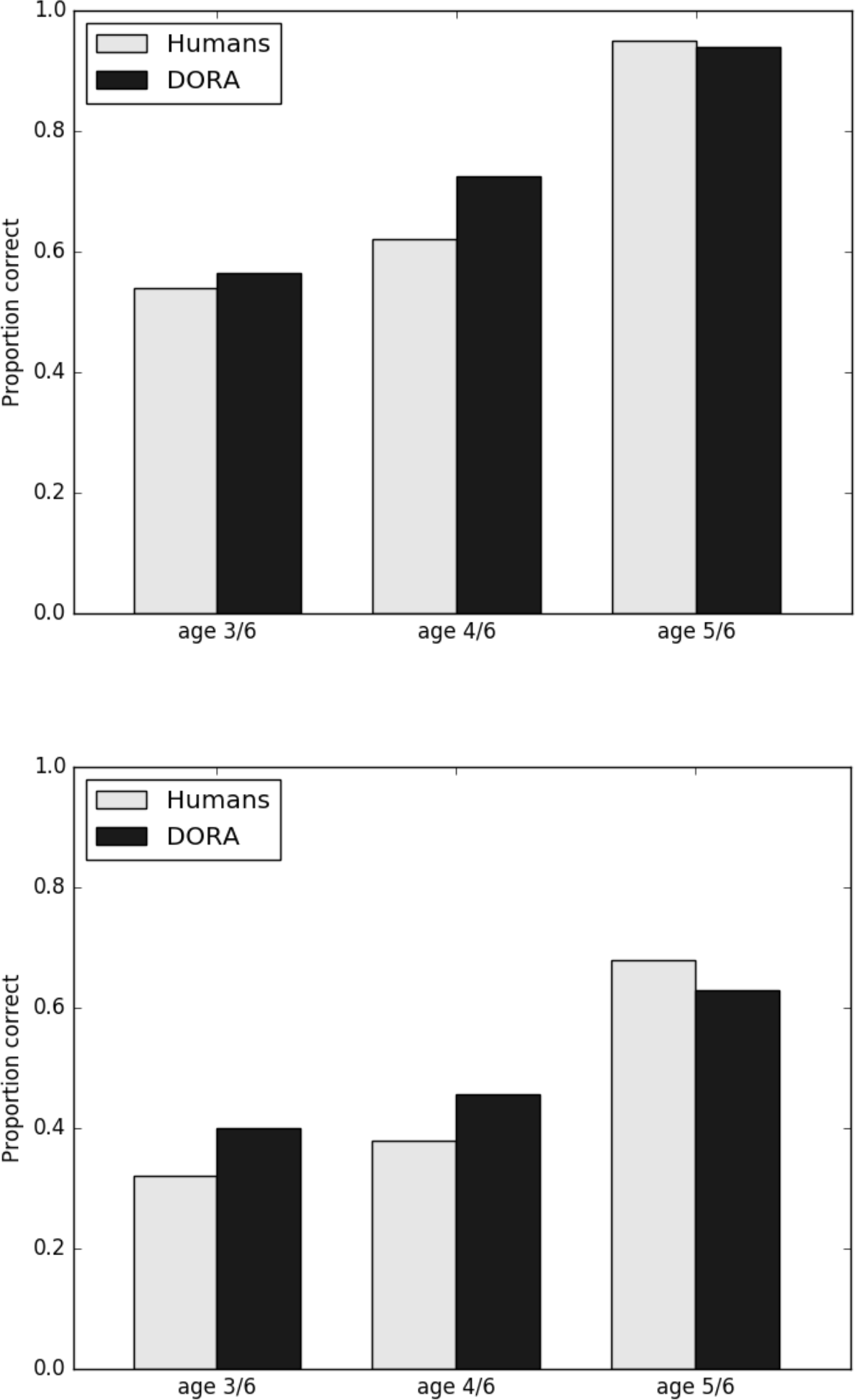
Results of simulation of Rattermann and Gentner (1998). (a) Performance of children and DORA on sparce trials. (b) Performance of children and DORA on rich trials.

As Fig. 21 shows, there is a close qualitative fit between DORA’s performance and the performance of the children in Rattermann and Gentner (1998). Initially, DORA, just as the 3-year-old children in the original study, had some trouble correctly mapping the items in the driver and the recipient, and struggled to solve the cross-mapping. As DORA learned more refined representations (after more training), like the 4-year-old children in the original study, DORA began to solve the sparse problems more successfully, while still struggling with the rich problems. Finally, like the 5-year-old children in the original study, after even more learning, DORA was quite successful at both rich and sparse trials, reaching ceiling level performance on the sparse problems. These simulation results indicate that, like humans, the trajectory of the development of DORA’s representations of relative dimensional magnitude undergoes a relational shift with learning. Additionally, the representations that DORA learns during its development support the same kind of performance on relational matching tasks that is evidenced by human children during their development.

## General Discussion

### Summary and Overview

How a system represents information tightly constrains the kinds of problems it can solve. Humans routinely solve problems that appear to require structured representations of stimulus properties and relations. As such, answering the question of how we acquire these representations has central importance in an account of human cognition.

Addressing the problem of how structured representations can be learned from experience requires solving three interrelated problems. First, the perceptual/cognitive system must learn to detect the basic featural *invariants* that remain constant across instances of a stimulus, object, event, or relation. Second, the system must isolate those invariants from the other properties of the objects engaged in the event or relation to be learned. Third, the system must learn a *predicate* representation of the relational properties (i.e., represent them as an explicit entity that can be bound to arbitrary and novel arguments while remaining independent of those arguments). We have previously (Doumas et al., 2008) proposed a theory for how the latter two problems might be solved. Here we have proposed a theory of how the first problem might be solved, and how the solutions might be integrated in a single system that learns structured representations of object properties and multi-place relations from experience with simple objects in the world.

Our theory is based on a set of 11 core theoretical claims outlined in Tables 1 and 2: (1) The invariant codes for SRM are a property of invariant neural responses that arise as a function of comparison. (2) During comparison compared items are co-activated, and co-activate their constituent distributed representations. (3) The similarity of the two compared items can be established by a measure of the match in the resulting firing pattern of the semantic properties. (4) The cognitive system learns invariant codes (nodes) of basic similarity as well as relative “same” and “different” by exploiting the invariant patterns of firing in (3). (5) The cognitive system detects relative magnitude by directly comparing absolute neural response in the neural system. (6) A specific and invariant firing pattern emerges when two items of greater and lesser magnitude are compared, and a different specific and invariant pattern emerges when two items of the same magnitude are compared. (7) The cognitive system learns invariant codes (nodes) for “same”/“different”, “more”/“less” by exploiting the invariant patterns of firing in (6). (8) Mature human mental representations are formally similar to a role-filler binding system (see Doumas & Hummel, 2005, and below), which reduces the problem of learning structured representations of multi-place relations to the comparatively simpler problems of learning single-place predicates, and then linking sets of corresponding single-place predicates together to form multi-place relational representations. (9) Comparison can lead to the discovery and predication of shared object properties. (10) A common vocabulary of representational primitives forms the basis for both predicates and their arguments. (11) Mapping sets of predicates of smaller arity can lead to the formation of higher arity (i.e., multi-place relational) structures.

We have instantiated our theory in a computational model called DORA. DORA is a symbolic-connectionist architecture based on Hummel and Holyoak’s (1997, 2003) LISA model. DORA is based on traditional connectionist computing principles, but uses time—as well as node activation—to carry information. DORA is, therefore, capable of solving the binding problem (Hummel, 2000). DORA can represent structured (i.e., symbolic) compositional propositions, and learn these representations from experience.

DORA uses Hebbian learning, time-based binding, comparison-based intersection discovery, and analogical mapping to learn to detect and produce invariant responses to instances of SRM, to learn and refine effective single-place predicate representations of object properties and relational roles, and to form whole multi-place relational structures from sets of co-occurring single-place predicates, and then further refine those relational representations. The result is a system that learns structured representations of relations from unstructured flat feature vector representations (i.e., traditional connectionist distributed representations) of objects with absolute properties.

We have shown that DORA accounts for a range of general and specific phenomena in human relational cognition. In particular, we have shown that DORA can learn structured representations of relations involving SRM from pixel images of shapes, and from Gabor patches that have been pre-processed to return absolute magnitude information. We have shown that the resulting representations meet the various requirements of human relational representations such as the capacity to support analogical mapping (including mapping nonidentical relations, and solving cross-mappings). Finally, we used DORA to account for a number of specific phenomena from the literature on the development of children’s reasoning with relations of similarity and relative magnitude, including accounting for the relational shift (see, e.g., Gentner, 2003). In short, we have demonstrated how a system can use the tools of statistical learning to succeed in overcoming the shortcomings of statistical learning, and how an important subset of the representations that account for the power of human cognition might be learned in the first place.

### Generalisability of the findings

In the reported results, we have focused our model on a specific class of relations—namely, those defined by SRM (initially over some dimension). A question that naturally arises is: will DORA’s learning algorithm extend to other kinds of relations (i.e., those non-obviously spatial relations)? The short answer is that it will. Starting with whatever regularities it is given or can calculate from the environment, DORA’s learning algorithm will isolate those invariants, learn structured representations that take arguments (i.e., functional predicates), and, where appropriate, compose them into relational structures. In short, given a set of invariants (or a means to calculate them), DORA’s learning mechanisms will produce explicit predicate and relational representations of those invariants. That is, DORA will learn structured representations of concepts based on their invariant properties, whether the invariants the system detects are instances of stimulus magnitude or romantic love, (see also, Doumas et al., 2008).

However, we might be able to use DORA to say a good deal more about the origins of non-obviously spatial relations. When one considers the relations that fall into the “non-obviously-spatial” set, like *chase*, or *support*, or *love*, it seems that they actually fall on a gradient of how spatial they actually are. For instance, it is quite straightforward to reduce a relation like *chase* to spatial properties: two objects where one is *in-front* of the other (where *in-front* is defined as *more-distance* from an origin point or *less-distance* from an end point), and the configuration is maintained through time. In fact, Michotte (1963) showed that participants would overwhelming interpret chasing occurring in a situation where two dots moved across a screen (or, in Michotte’s original version, two lights moved on a grid of lights) such that one stayed in front of the other as they moved. Relations such as *support* (one object *above* and in *contact* with another object), or *lift* (one object *supports* and *raises* another object), are similarly definable in spatial terms. Even a relation like *love* might reduce to spatial relations, though. In a study by Richardson & Spivey (2002), participants asked to use configurations of objects to represent a relation produced overwhelmingly similar spatial arrangements for relations like *love*, *admire*, and *hate*.

Previously we have proposed a complementary mechanism to DORA’s refinement algorithm that we termed compression (Doumas, 2005). In brief, compression is a form of chunking. During compression, multiple predicates or roles that are attached to the same object in an analog (or situation) fire together, and a recruited PO learns connections to the active semantics by Eq. 7. Compression allow DORA to combine multiple representations about the same object for less memory intensive processing. For example, if DORA encounters multiple situations in which one element is both *bigger* and *occludes* some second object, DORA can compress the roles *larger* and *occluder* about the first object, and compress the roles *smaller* and *occluded* about the second object to form a representation like *cover* (object1, object2). This procedure might, then, serve as a basis for DORA to combine the more spatial relations that it learns to form less obviously spatial relations.

A second question that might arise is whether the role-filler representational system that DORA employs is sufficient for supporting all the relations that humans learn. Again, the short answer is, at least in principle, yes. Formally, any multi-place relation is representable as a linked set of single-place predicates (Mints, 2001). Therefore, a role-filler system can be used to represent the relations that humans represent. The more pressing question, however, is whether a role-filler system fits with what we know about actual human relational representations. Again, the answer seems to be that it does. Certainly, models that employ role-filler type representations have successfully accounted for a large number of phenomena from human analogy making, relational learning, cognitive development, and learning (e.g., Doumas & Hummel, 2010; Doumas et al., 2008; Doumas et al., submitted; Hummel & Holyoak, 1997, 2003; Morrison et al., 2004; Lim, Sinnett, & Doumas, 2012, 2014; Livins, Spivey, & Doumas, 2015; Martin & Doumas, 2017; Morrison et al., 2012; Sandhofer & Doumas, 2008; Son, Doumas, & Goldstone, 2012). Moreover, access to and use of role-based semantic information is quite automatic in human cognition, including during memory retrieval (e.g., Gentner, Ratterman & Forbus, 1993; Ross, 1989), and analogical mapping and inference (Bassok & Olseth, 1995; Kubose, Holyoak & Hummel, 2002; Krawczyk, Holyoak & Hummel, 2008; Ross, 1987). Indeed, the meanings of relational roles influence relational thinking even when they are irrelevant or misleading (e.g., Bassok & Olseth, 1995; Ross, 1989). Role information appears to be an integral part of the mental representation of relations, and role-filler representations provide a direct account for why it is so. Moreover, role-filler systems appear uniquely capable of accounting for human’s abilities to overcome the n-ary restriction (see above).

Additionally, Livins and colleagues (2015) have shown that we can affect the direction of the relation by manipulating which item you look at first, for both obvious SRM relations, and the other kinds of relations. Livins et al. showed participants images depicting a relation that could be interpreted in different forms (e.g., *chase/pursued-by*, *lift/hang*). Before the image appeared on screen, though, a dot appeared on the screen drawing the participants attention to a location that one of the objects involved in the relation would appear. For example, the image might show a monkey hanging from a man’s arm, and the participant might be cued to the location where the monkey would appear. The relation that the participant used to describe the image was strongly influenced by the object that they attended to first. That is, if the participant saw the image of the monkey hanging from the man’s arm, and she was cued to the monkey, they would describe the scene using a *hanging* relation. However, if the participant was cued to the man, she would describe the scene using a *lifting* relation. This result follows directly from a system based on role-filler representations wherein complementary relations are represented by a similar set of roles, but the predicate, or role, that fires first defines the subject of the relation.

### Learning things we don’t already know

As noted above, Bayesian models of concept learning generally follow a learning-by-hypothesis-testing framework (e.g., Goodman et al., 2011; Kemp & Tenenbaum, 2009; Lake et al., 2015; cf Lu et al.’s, 2012). In these models, the system starts with a large set of representations and rules for combining them, and then learns combinations of these elements that best fit a given set of data. In other words, the model might learn a particular configuration of symbols to solve a problem, but this representation is can be generated in the model before any actual learning occurs. In fact, a configuration must be generated in order to enter as a candidate hypothesis in the learning algorithm.

The DORA model starts with representations of objects represented as flat feature vectors. Before learning, there are no functional predicate representations anywhere in the model. That is, at time *t*(0), the model has one type of representation, and not another. During learning, DORA learns new representations. Some of these new representations function like single-place—and eventually multi-place—predicates, and can be bound to arguments. That is, at time *t*(*n*), after learning, the model has another type of representation, one that is of a qualitatively different type from any representation present anywhere in the model at time *t*(0). The result is that the expressive power of the system has increased between *t*(0) and *t*(*n*). Of course, trivially, the capacity to *learn* these representations is present in the model (as it must be). There are architectural assumptions made (all detailed in the main text and in Appendix A), but these assumptions simply produce the capacity to learn predicate type representations. The representations themselves are specified nowhere within the model, and as a result, after learning, once the representations are specified, the model has a capacity to *represent* things that it simply did not have before learning. It is only after learning—including learning both representations that function like predicates and learning the networks necessary for SRM detection—that the structures necessary to represent relational propositions are present in the model. This change stands in stark contrast to models like those of Lake et al. (2015) and Kemp and colleagues (Kemp, 2012; Kemp & Tenenbaum, 2009, 2010), wherein the models start with explicit representations of all the primitive elements and data types necessary to express what is present in the training dataset or environment.

It is certainly possible that models like those proposed by Lake et al. (2015) and Kemp and colleagues (Kemp, 2012; Kemp & Tenenbaum, 2009, 2010) might be augmented with routines to generate the representations that they assume at the onset of learning. Perhaps solutions like DORA and BART might provide the means by which such useful representations can be generated on the first place. At the very least, Lake et al. and Kemp and Tenenbaum’s models might serve as very useful tools for addressing questions about what humans do with the representations that we have learned after they have acquired them.

Fodor (1976) provides a well-known argument for radical nativism, which has come to be known as “Fodor’s puzzle”. The argument—in its simplest form—progresses as follows: (1) concept learning is a process of hypothesis formulation and testing; (2) formulating a hypothesis about some *x*, requires having a concept of that *x*; (3) therefore, concepts are represented before they are learned, i.e., they are innate. In short, the argument holds that because the expressive power of the system at some time *t* determines the hypotheses that can be expressed and tested, the expressive power of a cognitive system is, primarily, fixed at onset. A cognitive system can learn to express combinations of existing concepts, and can learn to weight different combinations as more or less useful, but the constituent elements of these concepts and the rules for combining them are present in the system a priori. This argument has had a great deal of influence in the cognitive science community.

There have been some attempts to address Fodor’s problem. Generally, these accounts take umbrage with the statement that concept learning must be a process of hypothesis formation and testing. Specifically, the argument goes that we can test the utility of concepts via hypothesis testing, but we learn new concepts, by some other means. Such approaches usually propose some process for learning new primitives. While a system might begin with some set of initial representational primitives and means of combining them, by learning new primitives the system can formulate new concepts, which, in turn, can be tested for their utility. There are many such approaches in the literature including tuning (Landy & Goldstone, 2005), Quineian bootstrapping (Carey, 2009, in press), or random generation of representational elements.

An alternative idea is to extend the expressive power of a system by learning new data structures, or data types. For example, if a system that has representations of objects, but no predicate representations, when that system learns a predicate representation—even if that predicate is composed of primitives that already exist in the system—then the expressive power of that system has necessarily increased. Now the system can represent novel statements as a function of being able to represent instances wherein a predicate is *about* objects. The current proposal falls into this later camp.

### Learning degrees of similarity and difference in magnitude

While humans have the capacity to detect, respond to, and learn explicit structured representations of SRM, we also have the capacity to reason about relative differences of SRM. For instance, not only can we detect and represent that one item has more size than another item (i.e., is *larger*), but we can appreciate that the difference between the size of two items can be greater than the difference in size between two other items. That is, humans can represent and reason not only about first-order SRM, but also about second-order SRM (i.e., the similarity and relative magnitudes of different similarities and relative magnitudes). The capacity to detect and respond to second-order SRM falls directly out of the SRM calculation mechanism described above. While we did not describe the process in the main text because it had no bearing on the simulations presented, we describe the process in appendix A, and describe the solution in broad strokes below.

As described above, in DORA’s SRM detection procedure when two magnitude representations are compared and compete to become active, because greater magnitudes are coded by more units, the greater magnitude object will win the competition and the lesser magnitude object will lose. DORA exploits this result to bootstrap learning an invariant response to SRM. Interestingly, other useful invariant patterns also emerge during comparison. Specifically, when two such magnitude representations are compared, the time required for the network to settle into a “winner” and a “loser” varies directly as a function of how close the two magnitudes are. That is, by consequence of the comparison procedure, items that are more different will take less time to settle than items that are less different, and items that are entirely similar will take less time to settle than items that are almost entirely similar. For example, when the two magnitudes like size-2 and size-9 are compared, the unit with size-9 will get much more input and, thus, become much more active than the unit with size-2. By consequence, the unit with size-9 will quickly inhibit the unit with size-2, and “win” the competition to become active. On the other hand, when two magnitudes like size-4 and size-6 are compared, the unit with size-6 will get more input than the unit with size-4, but the difference in input between the two units coding size-6 and size-4 is smaller than the difference between the units coding size-9 and size-2. By consequence, the unit with size-6 will become more active than the unit with size-4, but it will take longer for the size-6 unit to inhibit the unit with size-4, and “win” the competition to become active. The time taken for the SRM circuit to settle on a “winner” is inversely related to the magnitude of the difference of magnitudes (second-order magnitude): Greater differences in magnitude take less time to settle, and smaller differences in magnitude take more time to settle.

The same relationship holds for instances when two items are the “same” (i.e., when the result of the magnitude comparison is two co-active units). When two units with identical magnitudes are compared, the network will quickly settle into a state where the two units are co-active, or quickly conclude that the two magnitudes are similar. By contrast, as the degree of difference becomes larger, the network will take longer to settle.

Importantly, these patterns are consistent across any magnitude comparison (greater magnitudes of difference (or identicality) result in quicker settling). The network can exploit these invariance responses to detect relative differences in the relative differences of magnitudes.^4^ The system can detect (and then respond to) second order SRM information with the same process that it uses to detect and respond simple SRM information. Moreover, this process may help bootstrap responding to magnitudes that are not coded by direct neural proxy. The invariant signal of taking longer to settle during comparison will serve as an invariant for greater difference, and the network will then learn a structured predicate representation of this invariance. This structured predicate representation may then be used (as in the simulations described) to reason about the similarity and relative magnitude of objects without direct analog magnitude codes. This feature of the model makes the interesting prediction that structured SRM representations of dimensions that are coded by analog neural proxy (e.g., visual size, auditory loudness) should be learned earlier than those that are not.

### Limitations and future directions

Humans routinely learn structured representations from experience. We offer an account of this fundamental process that is based on minimal standard assumptions of connectionist systems. Our account is, of course, limited in several ways. Below we outline some of the limitations of the current model, and propose some means of possibly addressing these limitations in future work.

First, the constraints on learning in our system are likely underdetermined; DORA learns when it can, and stores all the results of its learning. We have implemented a crude form of recency bias in our simulations (see above), but future work should focus on development of more principled mechanisms for constraining learning and storage. Such mechanisms might focus on either constraining when learning takes place, or on when the results of learning are stored for future processing. Most likely, though, it will be necessary to account for both.

Constraining when DORA learns amounts, essentially, to constraining when it performs comparison. We have previously proposed a number of possible constraints on comparison such as language (e.g., shared labels) and object salience, and have shown how direction to compare (i.e., instruction) serves as a very powerful constraint on learning (see Doumas et al., 2008; Doumas & Hummel, 2013). These constraints may also serve to limit when the results of learning are stored in memory. DORA might be extended or integrated with existing accounts of language or perceptual (feature) processing in order to implement such constraints (see, e.g., Martin & Doumas, 2017).

Perhaps more satisfyingly, both of these limitations might be successfully addressed by refining the control structure of the system. We see evaluating the quality of comparisons and of the representations that DORA learns as important potential constraints that the control process might impose. Reinforcement learning provides a very useful tool for implementing both of these constraints. Our current work is focused on developing a reinforcement based control structure in DORA. This structure has two primary focuses: (a) it evaluates the utility of a current comparison based on the reward from learning based on similar comparisons in the past. (b) It scores the utility of propositions in LTM based on their retrieval history and the reward from inferences based on these propositions in the past. This utility metric might then be used to prune representations in LTM.

Second, we lack a full account of how known predicates and relations are recognized during real-time processing. Previous work has shown that directing attention to particular features or particular agents has a pronounced effect on what relations are recognized in a scene (Livins & Doumas, 2015; Livins, Spivey, & Doumas, 2015; Livins, Doumas, & Spivey, 2016). For instance, directing attention to the height dimension (e.g., by having the participant move her head up and down) will drastically increase the chance that the participant will recognize a relation on that dimension (e.g., *above* (*x*,*y*); Livens, Doumas, & Spivey, 2016). Furthermore, drawing a participant’s attention to a particular object, unsurprisingly, makes recognizing a relation involving that object more likely, but also increases the probability of recognizing a relation in which that object is the subject (Livins & Doumas, 2015; Livens, Doumas, & Spivey, 2015). It remains an open question, however, how relational recognition is actually implemented in human cognition—although, see Livens et al., (2015) for a potential candidate.

The discovery of invariance has relevance beyond the few problems presented here. For example, detecting invariants in speech and language is a defining and unsolved problem in language acquisition and adult speech processing, including in automatic speech recognition by machines. Similarly, whether the generalization of grammatical rules can be fully accounted for in systems that rely on statistical learning alone remains contentious. The account of learning invariance from experience offered here, combined with principles like the compression of role information (Doumas, 2005), may present new computational vistas on these classic problems in the language sciences (see Martin, 2016 and Martin & Doumas, 2017 for further discussion). Systems with the properties of DORA, augmented by this subroutine and likely others, may offer an inroad to representational sufficiency across multiple domains, built from the same mechanisms and computational primitives.

### Conclusion

We have proposed a theory of how structured (i.e., symbolic) representations of object properties and relations can be learned from experience. We have shown how a computational model based on this theory can learn structured representations of SRM relations from simple pixel images of shapes and Gabor patches, processed to return absolute dimensional values. The resulting representations meet the requirements of human structured relational representations including solving cross-mapping, mapping non-identical relations, and overcoming the n-ary restriction. In addition, we have shown that the resulting model captures several specific phenomena from the literature on the development of children’s cognition about SRM. We have further proposed that relations based on SRM may, in fact, underlie many of the less obviously spatial relations (e.g., *chase*) that humans regularly reason about. As such, we have presented a theory of how what may be the most fundamentally human mental representations (see, e.g., Penn et al., 2008) might be learned from experience, and how these representations might be acquired by human learners during development.

## ACKNOWLEDGEMENTS

We thank Aaron Hamer and Hugh Rabagliati for helpful comments and discussions.

## Appendix A Details of SRM and DORA Operation

Like in LISA (Hummel & Holyoak, 1997), in DORA firing is organized by the *phase set*, or the set of units in the driver that is currently active and firing out of phase. We take the driver to be the focus of DORA’s attention, or what DORA is currently “thinking about”. The general sequence of events in DORA’s operation is outlined below. The details of these steps, along with the relevant equations and parameter values, are provided in the subsections that follow. Importantly, DORA is very robust to the values of the parameters (see Doumas et al., 2008). Throughout the equations in this Appendix, we will use the variable *a* to denote a unit’s activation, *n* its (net) input, and *w*_*ij*_ to denote the connection from unit *i* to unit *j*.

1. Bring an analog into the driver, *D*, (as designated by the user, or selected at random).

2. Initialize the activations of all units in the network to 0.

3. Select the firing order of propositions in *D* to become active. (In all the simulations described here, firing order is either set by the user or at random. However, see Hummel & Holyoak, 2003, for a detailed description of how a system like DORA can set its own firing order according to the constraints of pragmatic centrality and text coherence.)

4. Run SRM.

4.1. Do basic similarity calculation.

4.2. Do basic relative magnitude calculation.

5. Run phase set operations. Repeat the following until each RB in *D* has fired three times if mapping is licensed, or once, otherwise:

5.1. Select the proposition, *P*, at the head of the firing order.

5.1.1. If there are RB units in *P* that have not fired during the current phase set, select the RB, *RB*_*C*_, in *P*, in that is next in the firing order.

5.1.2. Otherwise, if there are PO units in pthat have not fired during the current phase set, select the PO unit, *PO*_*C*_, in *P*, that is next in the firing order.

5.2. Update the network in discrete time steps until the global inhibitor fires. On each time step *t* do:

5.2.1. Set input to *RB*_*C*_ or *PO*_*C*_ to 1.

5.2.2. Update modes of all P units in the recipient set, *R*. (We do not use higher-order relations in any of the simulations described in the text so the mode of P units is always at 1, however, we include this step for completeness.)

5.2.3. Update inputs to all token units in driver.

5.2.4. Update input to the PO inhibitors.

5.2.5. Update input to the RB inhibitors.

5.2.6. Update the local inhibitor.

5.2.7. Update the global inhibitor.

5.2.8. Update input to semantic units.

5.2.9. Update input to all token units in the recipient, *R*, and the emerging recipient, *N*.

5.2.10. Update activations of all units in the network.

5.2.11. Update all mapping hypotheses (if mapping is licensed).

5.2.12. Run retrieval (if retrieval is licensed).

5.2.13. Run comparison based learning (if comparison based learning is licensed).

5.2.13.1. If there are no active RB units in *P*, then run comparison-based predication.

5.2.13.2. If there are active RB units in *P*, then run refinement learning:

5.2.13.2.1. Run relational generalization.

5.2.13.2.2. Run relation formation.

5.2.13.2.3. Run predicate refinement.

6. Update mapping connections (if applicable).

7. Store results of learning (if any).

## Step 4.1: Do basic similarity calculation

Similarity calculation is performed when two PO units are compared and co-activated across the driver and recipient. During comparison, the activation of the compared POs is set to 1, and they pass activation to their constituent semantic units (see description of 5.2.8 below). This process continues for 25 iterations. Subsequently, similarity is calculated by the equation:

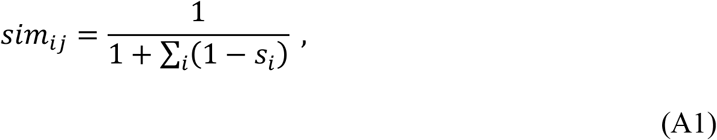

where, *sim*_*ij*_ is the 0 to 1 normalised similarity of PO unit *i* and PO unit *j*, and *s*_*i*_ is the activation of semantic unit *i*.

## Step 4.2: Do basic relative magnitude calculation

Magnitude calculation is performed when two POs from the same analog that both code some dimension are present in the driver. First, the activation of the two POs is set to 1, and they pass activation to the semantics coding the dimension they have in common (e.g., height), and any invariant SRM units (i.e., the semantics coding “same”, “different”, “more”, and “less”; initially POs are not connected to any of these semantics) to which they are connected. Input and activation of the semantic units is updated for 25 iterations (see steps 5.2.8 and 5.2.10, respectively).

*Step* 2: If no SRM unit is active above threshold (=.7), then the activation of the active magnitude semantics is clamped at 1, and the compared PO units compete to respond to the pattern of activation on the semantic units, otherwise, skip to *Step 4* (see below). Until the compared PO activation settles—remaining unchanged for 4 iterations—the network is updated in discrete time steps. On each time step, the input and activation of compared POs is updated. During magnitude calculation PO units are receiving input only from semantic units. Consequently, input to PO unit *i* is calculated using the equation,

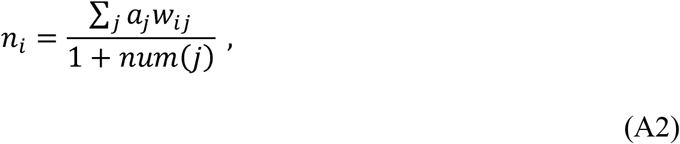

where *j* are semantic units connected to *i*, *w*_*ij*_ is the weight between units *i* and *j*, and *num(j*) is the number of semantic units *j*. PO units update their activation by the leaky integrator function:

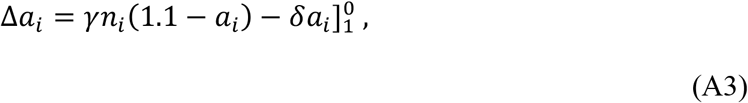

where Δ*a*_*i*_ is the change in activation of unit *i*, *a*_*i*_ is the activation of unit *i*, *γ* is a growth parameter, *n*_*i*_ is the net input to unit *i*, and *δ* is a decay parameter.

*Step* 3: Upon settling, the network is updated in discrete time steps until the local inhibitor fires. On each time step, the local inhibitor is updated (see step 5.2.6), the input and activation of compared PO units is updated (by Eqs. A2 and A3, respectively), and active POs learn connections to semantic units by Eq. 1 in the main text. Connection weights between POs and semantic units are limited to values between 0 and 1. If, upon settling, both compared POs are active, the “same” semantic unit becomes active until the local inhibitor fires. Othwerwise, if, upon settling, only one compared PO is active, the “more” semantic unit becomes active and the network is updated in discrete time steps (as above) until the local inhibitor fires. After the local inhibitor fires, the “less” semantic unit becomes active and the network is updated in discrete time steps (as above) until the local inhibitor fires.

*Step* 4: If any SRM units are active above threshold (=.7), these units compete via lateral inhibition, with the “more” semantic inhibiting “less” and “same”, and the “same” semantic inhibiting “more” and “less” (see Appendix B). After the SRM units settle, activation of the most active SRM unit is clamped to 1, and the network is updated in discrete time steps (as above) until the local inhibitor fires. If the “more” semantic unit was previously active, the “less” semantic unit becomes active and is clamped to 1. Again, the network is updated in discrete time steps (as in *Step 3*) until the local inhibitor fires.

During relative magnitude calculation, additional invariant patterns emerge. The relative magnitude calculation procedure described above also provides information about the relative differences of the relative differences in magnitude—or, second-order relative magnitude information (e.g., the distance between A and B is greater than the distance between C and D). More specifically, by consequence of the comparison procedure, items that are more different will take less time to settle than items that are less different, and items that are entirely similar will take less time to settle than items that are almost entirely similar. The network can exploit these invariance responses—in much the same way it exploits the different patterns of settling when items of different and similar magnitude are compared—to detect relative differences in the relative differences of magnitudes. That is, the system can detect (and then respond to) second order relative magnitude information with the same process that it uses to detect and respond simple relative magnitude information.

When two POs coding for some magnitude are compared as described directly above, second-order magnitude difference (i.e., the magnitude of the magnitude difference) can be approximated by the equation:

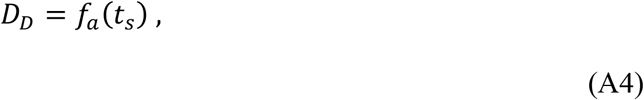

where, *D*_*D*_ is the second-order magnitude difference, *f*_*a*_ is a function, and *t*_*s*_ is a measure of the time to settling during the comparison process. For present purposes, *t*_*s*_ is a count of the number of iterations that the relative magnitude calculation required to settle.

The function *f*_*a*_ maps a value of *t*_*s*_ to a second-order magnitude difference, *D*_*D*_. Any number of functions, *f*_*a*_, will produce reasonable results, including a simple linear function. We do not claim to have an optimised *f*_*a*_ at present, but for present purposes we define the function as:

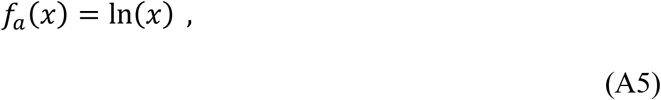

where, ln is the natural logarithm.

The same relative magnitude calculation procedure that produces responses to analog magnitudes also responds to values of *D*_*D*_ in precisely the same manner. As such, the same magnitude calculation method will produce implicit invariant responses to second-order magnitude differences, and explicit predicate representations of these responses can be learned as described in the main text and in the sections below.

## Step 5.2.2. Update mode of all P units in the recipient set

P units in all propositions operate in one of three modes: Parent, child, and neutral, as described by Hummel and Holyoak (1997, 2003). The idea of units firing in “modes” may sound “non-neural”, but Hummel, Burns & Holyoak (1994) describe how it can be accomplished with two or more auxiliary nodes with multiplicative synapses. A P unit in parent mode is operating as the linking element of a multi-place relational proposition. Parent P units excite and are excited by RBs to which they are downwardly connected (i.e., RB units that they link into multi-place relational propositions). In child mode, a P unit is acting as the argument of a higher-order proposition. Child P units excite and are excited only by RBs to which they are upwardly connected. In neutral mode, P units take input from all RBs to which they are upwardly and downwardly connected. The mode of P units in the driver are set at the beginning of each run by the rule given in the order of operations outline above. Each P unit *i* in *R* updates its mode, *m*_*i*_, according to:

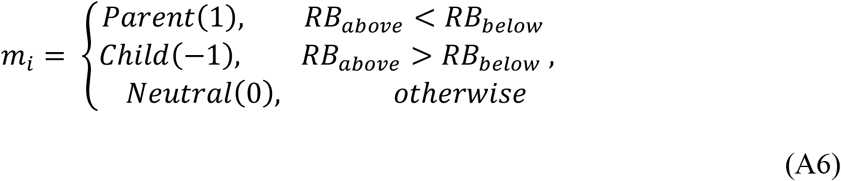

where *RB*_*above*_ is the summed input from all RB units to which *i* is upwardly connected (i.e., relative to which, *i* serves as an argument) and *RB*_*below*_ is the summed input from all RB units to which it is downwardly connected.

## Steps 5.2.3. Updating input to all token units in the driver

## P units

Each P unit *i* in *D* in parent mode updates its input as:

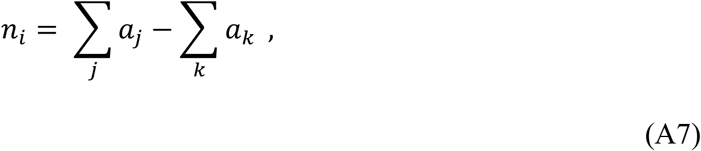

where *j* are all RB units below P unit *i* to which *i* is connected and *k* are all other P units in *D* that are currently in parent mode. P units in *D* in child mode update their inputs by:

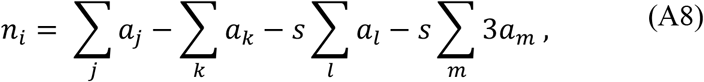

where *j* are RB units to which *i* is upwardly connected, *k* are other P units in the driver that are currently in child mode, *l* are all PO units in the driver that are not connected to the same RB as *i*, and *m* are all PO units that are connected to the same RB (or RBs) as *i*. When DORA is operating in binding-by-asynchrony mode, *s* = 1; when it is operating in binding-by-synchrony mode (i.e., like LISA), s = 0.

## RB units

RB units in the driver update their inputs by:

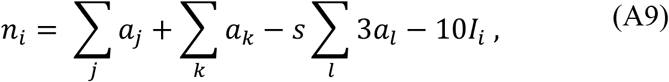

where *j* are all P units in parent mode to which RB unit *i* is upwardly connected, *k* are all PO units connected to *i*, *l* are all other RB units in D, *and I*_*i*_ is the activation of the RB inhibitor yoked to *i*.

## PO units

PO units in the driver update their input by:

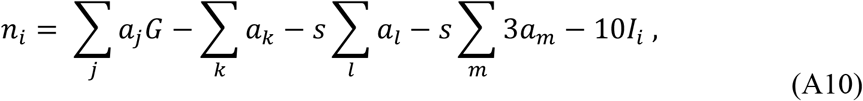

where *j* are all RB units to which PO unit *i* is connected, *G* is a gain parameter attached to the weight between the RB and its POs (POs learned via DORA’s comparison based predication algorithm—and thus with mode=1; see section 5.2.13.1 below—have *G*=2 and 1 otherwise), *k* are P units in *D* that are currently in child mode and not connected to the same RB as *i*, *l* are all PO units in the driver that are not connected to the same RB as *i*, *m* are PO units that are connected to the same RB (or RBs) as *i*, and *I*_*i*_ is the activation of the PO inhibitor yoked to *i*. When DORA is operating in binding-by-asynchrony mode, *s* = 1; when it is operating in binding-by-synchrony mode (i.e., like LISA), s = 0.

## Steps 5.2.4 and 5.2.5. Update input to the PO and RB inhibitors

Every RB and PO unit is yoked to an inhibitor unit *i*. Both RB and PO inhibitors integrate input over time as:

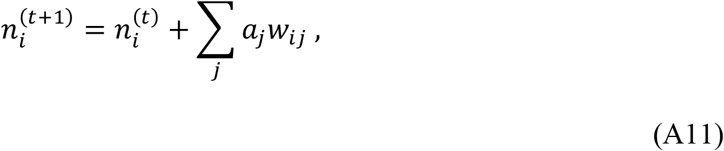

where *t* refers to the current iteration*, j* is the RB or PO unit yoked to inhibitor unit *i*, and *w*_*ij*_ is the weight between RB or PO inhibitor *i* and its yoked RB or PO unit (set to 1). Inhibitor units become active (*a*_*i*_ = 1) when *n*_*i*_ is greater than the activation threshold (=220). RB inhibitors are yoked only to their corresponding RB. PO inhibitors are yoked both to their corresponding PO and all RB units in the same analog. As a result, at any given instant, PO inhibitors receive twice as much input as RB inhibitors, and reach their activation threshold twice as fast. POs, therefore, oscillate twice as fast as RBs. PO and RB inhibitors establish the time-sharing that carries role-filler binding information and allows DORA to dynamically bind roles to fillers. All PO and RB inhibitors become refreshed (*a*_*i*_ = 0 and *n*_*i*_ = 0) when the global inhibitor (Γ_G_; described below) fires.

## Steps 5.2.6 and 5.2.7. Update the local and global inhibitors

The local and global inhibitors, Γ_L_ and Γ_G_ respectively (see e.g., Horn and Usher, 1990; Horn et al., 1992; Usher and Nieber, 1996; von der Malsburg and Buhman, 1992), serve to allow units in the recipient to keep pace with firing of units in the driver. The local inhibitor is inhibited to inactivity (Γ_L_ = 0) by any PO in the driver with activation above Θ_L_ (= 0.5), and becomes active (Γ_L_ = 10) when no PO in the driver has an activity above Θ_L_. During asynchronous binding, the predicate and object POs time-share. There is a period during the firing of each role-filler pair after the one PO fires and before the other PO becomes active when no PO in the driver is very active. During this time the local inhibitor becomes active and inhibits all PO units in the recipient to inactivity. Effectively, Γ_L_ serves as a local refresh signal, punctuating the change from predicate to object or object to predicate firing in the driver, and allowing the units in the recipient to keep pace with units in the driver.

The global inhibitor works similarly. It is inhibited to inactivity (Γ_G_ = 0) by any RB in the driver with activation above Θ_G_ (= 0.5), and becomes active (Γ_G_ = 10) when no RB in the driver is active above threshold. During the transition between RBs in the driver there is a brief period when no driver RB are active above Θ_G_. During this time Γ_G_ inhibits all units in the recipient to inactivity, allowing units in the recipient to keep pace with units in the driver.

## Step 5.2.8. Update input to semantic units

Semantic units update their input as:

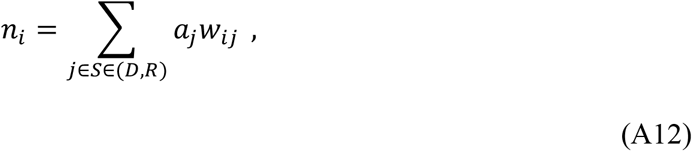

where *j* is all PO units in *S*, which is the set of propositions in the driver, *D*, and recipient *R*, and *w*_*ij*_ is the weight between PO unit *j* and semantic unit *i*.

## Step 5.2.9. Update input to token units in the recipient and the emerging recipient

Input to all token units in the recipient and emergent recipient are not updated for the first 5 iterations after the global or local inhibitor fires. This delay in update occurs in order to allow units in the recipient and emergent recipient to respond to the pattern of activation imposed on the semantic units by the driver PO unit that wins the competition to become active after an inhibitor fires.

## P units

P units in parent mode in the recipient update their inputs by:

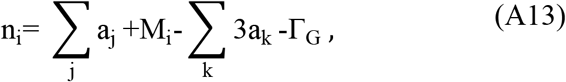

where *j* are all RB units to which P unit *i* is downwardly connected, *k* are all other P units in the recipient currently in parent mode and *M*_*i*_ is the mapping input to *i*:

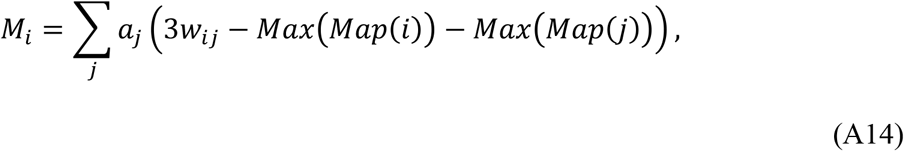

where *j* are token units of the same type as *i* in the driver (e.g., if *i* is a RB unit, *j* is all RB units in the driver), *Max*(*Map*(*i*)) is the highest of all unit *i*’s mapping connections, and *Max*(*Map*(*j*)) is the highest of all unit *j*’s mapping connections. As a result of Eq. A14, an active token unit in the driver will excite any recipient unit to which it maps, and inhibit all recipient units of the same type to which it does not map.

P units in child mode in the recipient update their inputs by:

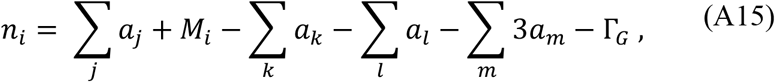

where *j* are all RB units to which *i* is upwardly connected, *M*_*i*_ is the mapping input to *i*, *k* are all other P units in the recipient currently in child mode, *l* are POs in the recipient that are not connected to the same RB (or RBs if *i* is connected to multiple RBs) as *i*, and *m* are PO units connected to the same RB (or RBs) as *i*.

## RB units

RB units in the recipient update their input by:

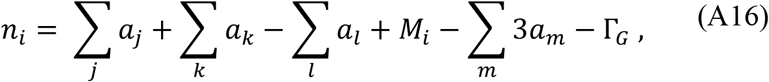

where *j* are P units currently in parent to which RB unit *i* is upwardly connected, *k* are P units currently in child mode to which *i* is downwardly connected, *l* are PO units to which unit *i* is connected, *M*_*i*_ is the mapping input to *i*, and *m* are other RB units in the recipient.

## PO units

PO units in the recipient update their input by:

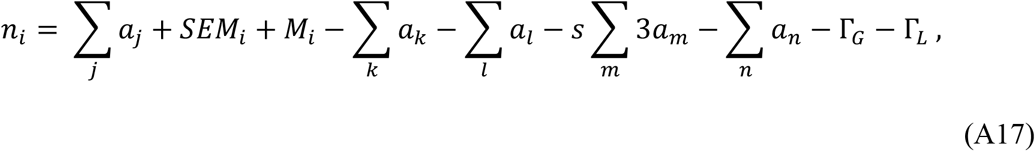

where *j* is RB units to which PO unit *i* is connected (input from *j* is only included on phase sets beyond the first), *SEM*_*i*_ is the semantic input to unit *i, M*_*i*_ is the mapping input to unit *i, k* is all PO units in the recipient that are not connected to the same RB (or RBs if unit *i* is connected to multiple RBs) as *i, l* is all other P units in the recipient currently in child mode that are not connected to the same RB (or RBs) as *i, m* is PO units connected to the same RB (or RBs) as *i*, and *n* is RB units in the recipient to which unit *i* is not connected. When DORA is operating in binding-by-asynchrony mode, *s* = 1; when it is operating in binding-by-synchrony mode (i.e., like LISA), *s* = 0. *SEM*_*i*_, the semantic input to *i*, is calculated as:

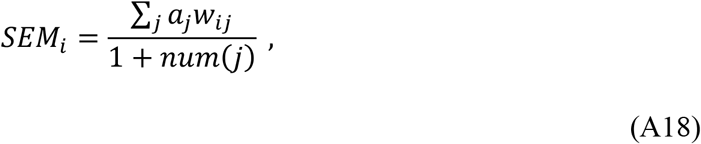

where *j* are semantic units, *w*_*ij*_ is the weight between semantic unit *j* and PO unit *i*, and num(*j*) is the total number of semantic units *i* is connected to with a weight above *θ* (=0.1). Semantic input to POs is normalized by a Weber fraction so that the PO unit that best matches the current pattern of semantic activation takes the most semantic input, and semantic input is not biased by the raw number of semantic features that any given PO is connected to (see Hummel & Holyoak, 1997, 2003; Marshall, 1995).

## Step 5.2.10. Update activations of all units in the network

All token units in DORA update their activation by Eq. A3.

The value of growth parameter, γ, is 0.3, and the value of decay parameter, δ, is 0.1.

As noted above, semantic unit activations are divisively normalized, and semantic units update their activation by the equation:

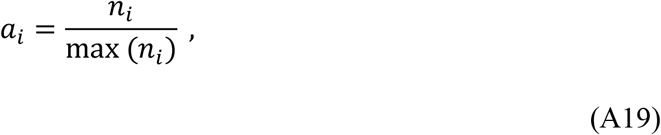

where *a*_*i*_ is the activation of semantic unit *i*, *n*_*i*_ is the net input to semantic unit *i*, and max(*n*_*i*_) is the maximum input to any semantic unit. There is physiological evidence for divisive normalization in the feline visual system (e.g., Albrecht & Geisler, 1991; Bonds, 1989; Heeger, 1992) and psychophysical evidence for divisive normalization in human vision (e.g., Foley, 1994; Thomas & Olzak, 1997).

RB and PO inhibitors, *i*, update their activations according to a threshold function:

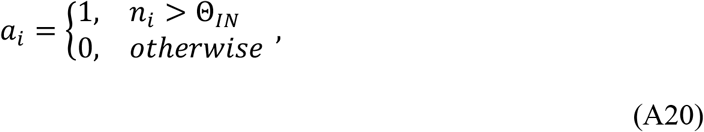

where Θ_*IN*_ = 220.

## Step 5.2.11. Update all mapping hypotheses

DORA’s mapping algorithm is adopted from Hummel and Holyoak (1997, 2003). During the mapping process, DORA learns mapping hypotheses between all token units in the driver and token units of the same type in the recipient (i.e., between P units, between RB units and between PO units in the same mode [described below]). Mapping hypotheses initialize to zero at the beginning of a phase set. The mapping hypothesis between an active driver unit and a recipient unit of the same type is updated by Eq. 3 in the main text (i.e., mapping hypotheses update via a Hebbian learning rule).

## Step 4.2.12 Run retrieval

DORA uses a variant of the retrieval routine described by Hummel and Holyoak (1997). During retrieval propositions in the driver fire as described above for one phase set. Units in the dormant/LTM set become active in response to the patterns of activation imposed on the semantics by active driver POs. After all RBs in the driver have fired once, DORA retrieves propositions from LTM probabilistically using the Luce choice axiom:

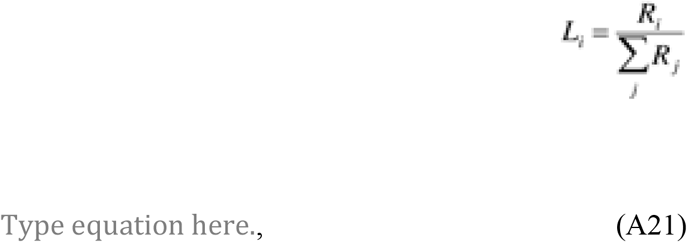

where *L*_*i*_ is the probability that P unit *i* will be retrieved into working memory, *R*_i_ is the maximum activation P unit *i* reached while during the retrieval phase set and *j* are all other P units in LTM. If a P unit is retrieved from LTM, the entire structure of tokens (i.e., RBs, POs, and P units that serve as arguments of the retrieved P unit) are retrieved into working memory.

## Step 5.2.13. Run comparison based learning

In the current version of the model, learning is licensed whenever 70% of the driver token units map to recipient items (this 70% criterion is arbitrary, and in practice 100% of the units nearly always map; in the general discussion section of the main text, we discuss some additional potential constraints on learning). If learning is licensed DORA invokes either comparison-based-predication or refinement learning. If the driver contains single objects, not yet bound to any predicates (i.e., each RB in the driver is bound only to a single PO), then comparison based predication is licensed. Otherwise, refinement learning is licensed.

## Step 5.2.13.1. Comparison-based predication

As detailed in the text, during comparison-based predication (CBP) for each PO in the driver that is currently active, and maps to a unit in the recipient with a mapping connection above the threshold Θ_MAP_ (=0.5), DORA recruits a new PO unit (i.e., a PO connected to no semantic features) in the recipient. The mode of the existing PO units in both the driver and recipient is set to 0 and the mode of the newly inferred PO is set to 1. The mode of PO units is used to distinguish POs acting as objects from POs acting as predicates (i.e., those POs learned via comparison-based learning). Learned predicate POs have a higher gain on their connection weights to RBs than do object POs (see section 5.2.3 above). This gain allows predicates to fire before objects during asynchronous binding, and reflects our assumptions that humans favour things they learn when making inferences (see, e.g., Goldstone et al., 1991). The mode of POs is is also important for assuring mappings from predicates to other predicates and from objects and other objects when DORA uses LISA-like synchrony binding (i.e., when roles and their fillers fire in synchrony). Although we do not use synchrony based binding for the reported simulations, we mention it here and implement it in our code for the purposes of completeness. DORA learns connections between the new PO and all active semantics by the Eq. 7 in the main text. During CBP, DORA also infers a new RB unit in the recipient. The activation of each inferred unit is set to 1, and remains at 1 until Γ_G_ or Γ_L_ fires. DORA learns a connection with a weight of 1 between corresponding active token units (i.e., between P and RB units. and between RB and PO units) that are not already connected.

## Step 5.3.13.2. Refinement Learning

During refinement learning DORA first runs its relation formation routine then its predicate refinement routine.

*Step 5.2.13.2.1: Relational generalisation*. Relational generalisation is the process by which relational inferences are made in DORA. The relational generalisation algorithm is adopted from that used in Hummel and Holyoak’s (2003) LISA model. The relational generalisation algorithm in the current model is the same as the one used in Doumas et al. (2008). While we do not used relational generalisation for any of the simulations included in the current paper, we have used the algorithm in previous simulations (e.g., Doumas et al, 2008; Doumas, Morrison, & Richland, submitted), and so it is included here for the purposes of completeness.

*Step 5.2.13.2.2: Relation formation*. As described in the main text, when DORA successfully maps sets of role-filler bindings in the driver to sets of role-filler bindings in the recipient, the resulting pattern of firing on the recipient RB units is exactly like what would emerge from RB units joined by a common P unit (i.e., mapped RBs in the recipient fire out of synchrony but in close temporal proximity, and within each mapped RB, mapped POs fire out of synchrony but in close temporal proximity). During relation formation DORA exploits this temporal pattern to link the recipient RBs (along with their respective POs) into a full proposition—i.e., a multi-place relation. This process is accomplished as a case of SSL. When an RB in the recipient becomes active, if no P units are active in the recipient, then a P unit is recruited in the recipient via SSL. The newly recruited P unit remains active (activation=1) until the end of the phase set and learns connections to active RBs: The P unit learns connections of 1 to RBs active above threshold(=.8). When the phase set ends, connection weights between the new P unit *i* and any RBs to which it has connections, *j*, are updated by the equation:

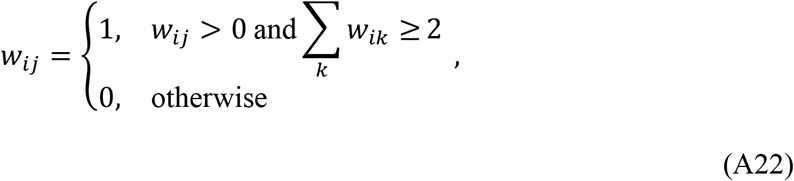

where *w*_*ij*_ is the connection weight between P unit *i* and RB unit *j*, and *w*_*ik*_ is the connection weight between *i*, and RB unit *k* where *k* is all RB units (including *j*) in the recipient. Essentially, if the new P has at least two connections to RB units (and so sum over *k* of *w*_*ik*_ is greater than or equal 2), then DORA retains the connections between the recruited P and all RBs to which it has learned connections; if the sum is less than two, then it discards the connections (along with the P unit). This convention ensures that DORA does not learn superfluous P units (i.e., Ps that connect only to a single RB). The exact same result is accomplished by learning connections between P and RB units by Hebbian learning, and then truncating all connections if the sum of connections to the newly recruited P unit is not greater than 1.

*Step 5.3.13.2.3: Predicate refinement*. As detailed in the text, during predicate refinement DORA learns a refined representation of mapped propositions or role-filler sets. For each PO in the driver that is currently active, and maps to a unit in the recipient with a mapping connection above the threshold Θ_MAP_ (=0.7), DORA infers a PO unit connected to no semantic features in the emerging recipient with a mapping connection to the active driver unit. DORA learns connections between the new PO and all active semantics by Eq. 7 in the main text. DORA also licenses self-supervised learning (SSL). During SSL, DORA infers token units in the emerging recipient that match active tokens in *D* (the driver). Specifically, DORA infers a structure unit in the emerging recipient in response to any unmapped token unit in *D*. If unit *j* in *D* maps to nothing in the emerging recipient, then when *j* fires, it will send a global inhibitory signal to all units in the emerging recipient (Eq. A14). This uniform inhibition, unaccompanied by any excitation in the recipient us a signal that DORA exploits, and infers a unit of the same type (i.e., P, RB, PO) in the emerging recipient. Inferred PO units in the emerging recipient have the same mode as the active PO in the driver. The activation of each inferred unit in the emerging recipient is set to 1. DORA learns connections (weight=1) between corresponding active tokens in the emerging recipient (i.e., between P and RB units. and between RB and PO units). To keep DORA’s representations manageable (and decrease the runtime of the simulations), at the end of the phase set, we discard any connections between semantic units and POs whose weights are less than 0.1.

## Step 6. Update mapping connections

Mapping connections are updated at the end of each phase set. First, all mapping hypotheses are normalized by Eq. 4 in the main text. That is, each mapping hypothesis is normalised divisively: Each mapping hypothesis, *h*_*ij*_ between units *i* and *j*, is divided by the largest hypothesis involving either unit *i* or *j*. Next each mapping hypothesis is normalized subtractively: The value of the largest hypothesis involving either *i* or *j* (not including *h*_*ij*_ itself) is subtracted from *h*_*ij*_. The divisive normalization keeps the mapping hypotheses bounded between zero and one, and the subtractive normalization implements the one-to-one mapping constraint by forcing mapping hypotheses involving the same *i* or *j* to compete with one another (see Hummel & Holyoak, 1997). Finally, the mapping weights between each unit in the driver and the token units in the recipient of the same type are updated by Eq. 5 in the main text.

## Appendix B Network for neural coding of invariant magnitude response

As noted in the main text, there are several ways to instantiate a neural response to the invariant signal produced during similarity and magnitude comparison. Here we present one solution, but—also as noted in the main text—we make no strong theoretical claims about the veracity of any particular instantiation. Rather, our claim is that the neural system learns to respond to the invariant pattern of firing that occurs when similar and different magnitudes are compared, and uses this response as the basis for learning invariant semantic features that support implicit SRM processing, and can serve as the basis for learning structured (i.e., predicate) explicit SRM representations.

Figure B1 depicts the magnitude response network. The network instantiates the magnitude comparison operation described in Appendix A, and behaves equivalently to that description. This network is neutrally plausible—based on classical connectionist computing principles—and is learnable via unsupervised learning. In the following we first describe the behavior of the network, then a basic algorithm for learning this network.

**Figure B1.**
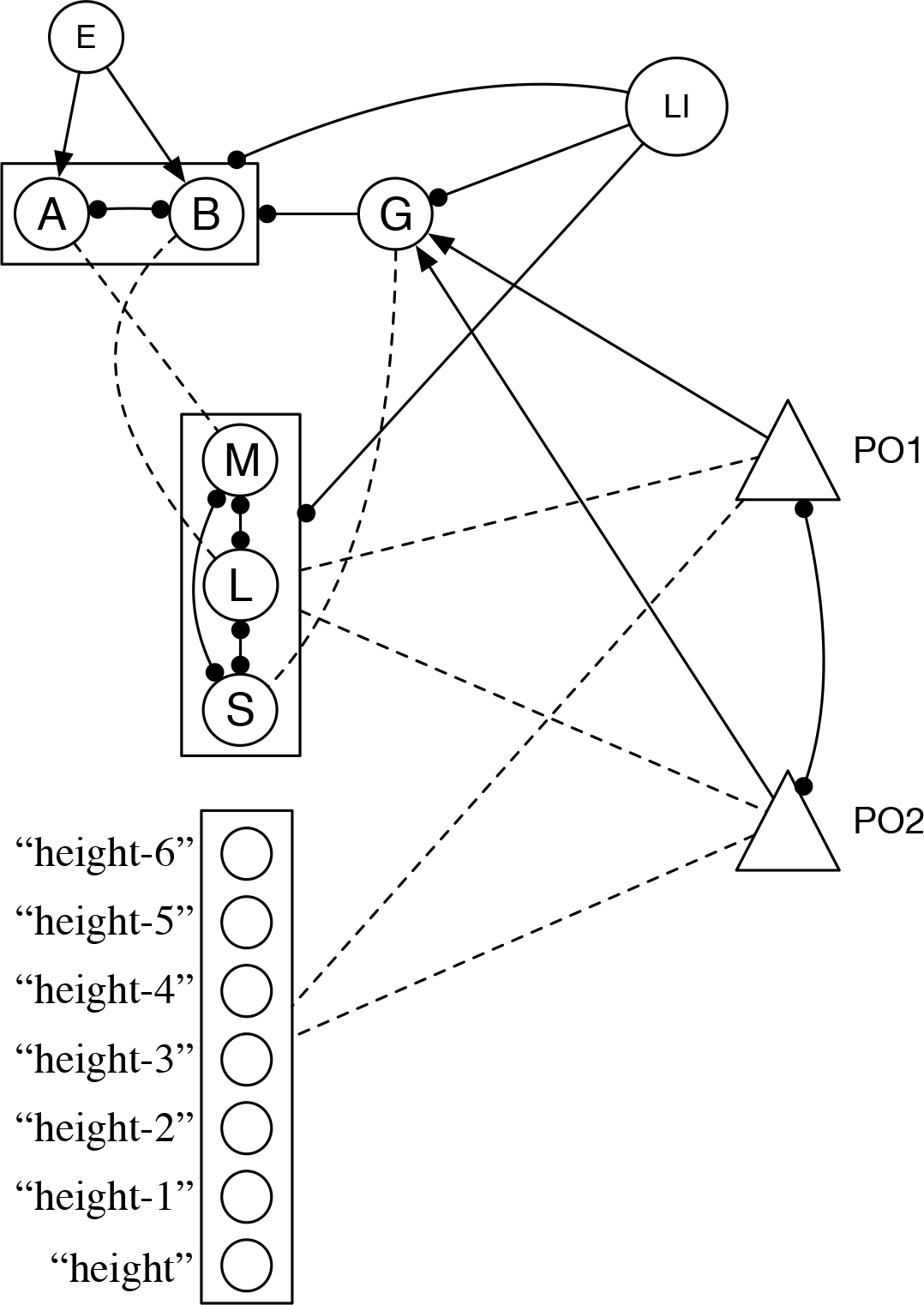
A network that responds to the invariant patterns of firing that occur during magnitude comparison. Node (A) marks early in comparison process. Node (B) marks later in the comparison process. Node (G) is a gating unit that fires only when receiving input from multiple units, and marks two active POs. Node (E) fires once the activation of the PO units (triangles) has settled (see main test and Appendix A). Node (M) comes to stand as the invariant of ‘more’; node (L) comes to stand as the invariant of ‘less’; node (S) comes to stand as the invariant of ‘same’. LI=local inhibitor. Solid lines with circles at either endpoint specify two-way inhibitory connections. Solid lines with a circle at only one endpoint specify one-way inhibitory connections where the node at the circle end is inhibited. Solid lines with arrows at one endpoint specify one-way excitatory connections where the node at the arrow end is excited. Dashed lines indicate excitatory connections between modified via unsupervised Hebbian learning, as described in the text. Where nodes are contained in a box connections to and from that box apply to all units in that box.

During magnitude comparison, PO units (nodes labelled PO1 and PO2 in Fig. A1) compete to become active. PO units are connected to semantic units indicating their absolute magnitude, with greater magnitudes encoded by larger numbers of units (see main text). As a consequence, when the two PO units code different absolute magnitudes, the PO unit connected to the greater magnitude will win the completion to become active, and inhibit the PO unit connected to the lesser magnitude.

When the units settle, node E in Fig. A1 fires. As can be seen in Fig. A1, node E passes activation to nodes A and B. Node A is randomly more strongly connected to E and becomes active more quickly, inhibiting node B to inactivity. Node A also activates to the “more” semantic, and the active PO unit learns a connection to the “more” semantic (see main text). When the inhibitor on the active PO fires, the active PO unit is inhibited to inactivity, and the local inhibitor (LI) fires (see main text and Appendix A). The LI inhibits unit A, allowing node B to become active as the next PO unit also becomes active. Node B inhibits node A, and passes activation to the “less” semantic. The active PO unit learns a connection to the “less” semantic.

When both PO units code for the same semantic, they settle into a stable state of mutual activation. Two active PO units will activate gating node G. Node G inhibits nodes A and B, and passes activation the “same” semantic. The active PO units learn connections to the “same” semantic.

The magnitude circuit consists of nodes A, B, G, and the magnitude semantics. This circuit is learned via an unsupervised learning algorithm. Nodes A and B follow the activation function:

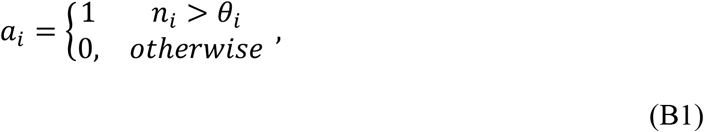

where *a*_*i*_ is the activation of threshold unit *i*, *n*_*i*_ is the input to unit *i*, and *θ*_*i*_ is the threshold of unit *i*. The input to unit *i* is calculated by:

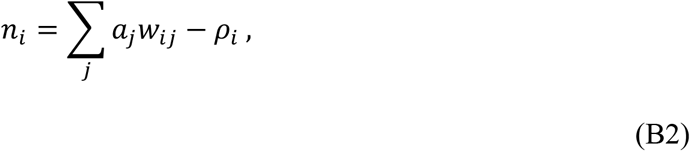

where, *n*_*i*_ is the input to unit *i*, *a*_*j*_ is the activation of unit *j* connected to unit *i*, *W*_*ij*_ is the connection weight between units *i* and *j*, and *ρ*_*i*_ is the refraction of unit *i*. Node G is a gating unit (i.e., it has a threshold greater than 1). Refractions for A and B units have a base level, *θ*_*b*_, are adjusted as a function of time since last firing by:

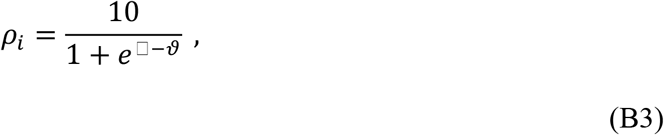

where, *x* is the number of iterations since unit *i* has fired, and *ϑ* is the threshold of PO inhibitors (=110). Connection weights between units A and B and unit E, and connection weights between units A, B, G and semantics are initially set to a random value between 0 and 1. PO units and semantic units behave as described in Appendix A with the exception that semantic units M, L, and S divisively and subtractively normalise their input.

During learning, connection weights between units A, B, G and semantic units are adjusted through Eq. 6 in the main text. Learning begins 3 iterations after POs settle. In short, after learning, the unit most strongly connected to unit E will become the A unit, and the unit with the next strongest connection to E will become the B unit. Similarly, the semantic unit that ends up the most strongly connected to A becomes the semantic invariant for “more”, the semantic unit that ends up the most strongly connected to B becomes the semantic invariant for “less”, and the semantic unit that ends up the most strongly connected to G becomes the semantic invariant for “same”. Connections between semantic units and active PO units are updated by Eq. 6 in the main text.

## Appendix C Image pre-processor

## Geometrical figures

A 3-layer feedforward neural network was trained to recognize 36 geometrical figures. The figures were 30 × 30 pixel images that varied across five dimensions: (1) shape: circle, rectangle, triangle, square. (2) Width: nine different levels. (3) Height: 9 different levels, with the rectangle always having height = width/2 and the rest of the shapes having height = width. (4) Area: 9 different levels, with the width and height constraints described yielding 27 possible values of area across shapes.; (5) color: 5 different levels.

The feedforward neural network had 900 input units, 150 hidden units and 54 output units. The output layer was composed of 4 groups of units: the first group had four units coding for shape, the second group had nine units coding for with, the third group had 14 units coding for height, and the fourth group had 27 units coding for area.

The network was trained using back-propagation. The cost function used was cross-entropy with no regularization. When evaluating the accuracy of the network, for each group of output units the network’s output was taken as 1 for the unit with the largest activation and zero for all other units. The network was trained for 2400 epochs with a learning rate of 0.1 and batch size of 36, yielding a classification accuracy of 1.

## Gabor patches

A 3-layer feedforward neural network was trained to recognize 48 Gabor patches. This images corresponded to 30 × 30 pixel grayscale images that varied across 3 dimensions: (1) Frequency of stripes: 4 levels from 2 to 8 (increasing in steps of 2). (2) Orientation of stripes: 6 levels from 0 to 180 (increasing in steps of 30 degrees). (3) Window: square window, gaussian window.

The feedforward neural network had 900 input units, 30 hidden units and 12 output units. The output layer was composed of three groups of units: the first group had four units coding for frequency of stripes, the second group had six units coding for orientation of stripes, and the third group had two units coding for window.

The network was trained using back-propagation. The cost function used was Cross-Entropy with no regularization. When evaluating the accuracy of the network, for each group of output units the network’s output was taken as 1 for the unit with the largest activation and cero for all other units. The network was trained for 300 epochs with a learning rate of 0.2 and batch size of 30, yielding a classification accuracy of 1.

## Footnotes

1 While we use labels for semantic units, the specific content of the units coding for a property are unimportant to DORA (in fact, the model is actually unaware of the labels given to semantic units). So long as there is something common across the units representing a set of objects, DORA can learn an explicit representation of this commonality. That is, for the purposes of DORA’s learning algorithm, all that matters is there is something invariant across instances of a *container* (which there must be for us to learn the concept), and that the perceptual system is capable of responding to this invariance (which, again, there must be for us to respond similarly across instances of containment in the world; for a more complete discussion of the role of invariance in perception see, e.g., Biederman, 1987; Hummel, Kellman, & Burke, 1999).

2 While we describe mapping connections as instantiated as weights between units for the purposes of expositional clarity, we are not committed to this instantiation. Mapping connections might also be instantiated as units with fast modifiable connections (as they are in the current model). For the purposes of the simulations described in this paper, the particular instantiation (weighted connections vs. units) does not have an effect. See Knowlton et al. (2012) for a more comprehensive discussion.

3 Following from LISA (Hummel & Holyoak, 2003), we adopt a basic strategy during the mapping process. Previous work has shown that people place different weights on featural and relational similarity during different kinds of tasks. For example, Medin, Goldstone, and Gentner (1994) had participants perform a similarity-judgment task in which participants aligned structurally complex objects (e.g., insects with many parts). Medin and colleagues found that participants placed greater emphasis on features (and ignoring relations) early in processing, but that their emphasis shifted strongly to relations (and ignoring surface features) later in mapping and similarity judgment. More generally, participants differentially emphasized features and relations during different parts of a task, and attention to features or relations showed an inverse relationship (e.g., as relational emphasis went up, attention to surface features went down). Accordingly, Hummel and Holyoak (2003) developed a strategic relational processing routine for LISA. When multiple relational propositions are placed in LISA’s WM (i.e., the driver) simultaneously, LISA ignores the semantic features of the object units. The routine is based on the assumption that people default to considering relations singly, and think about multiple relations simultaneously only to make inferences based on higher-order structures (i.e., relations). This constraint allowed LISA to simulate multiple findings on relational integration in human analogy making (Hummel & Holyoak, 2003). We adopt the same constraint in DORA’s mapping routine. While this constraint is not important for any of the simulations described herein, we include it for the purposes of completeness.

4 Similarly, when any two representations are over-laid (i.e., compared), the extent to which one matches the other (calculable by Hamming distance, or by some more complex metric like the one described in Appendix A) serves as another implicit signal for detection of more and less.

